# Molecular basis of nick ligation in the nucleosome by DNA Ligase IIIα

**DOI:** 10.64898/2026.03.31.714456

**Authors:** Daniel J. Boesch, Nadia I. Martin, Chantal A. Kontor, Ashlee G. Nguyen, Alan E. Tomkinson, Bennett Van Houten, Natacha M. Gillet, Emmanuelle Bignon, Amy M. Whitaker, Tyler M. Weaver

## Abstract

Eukaryotic genomic DNA is packaged into chromatin through a fundamental repeating unit known as the nucleosome core particle. Within this chromatin context, genomic DNA is constantly exposed to endogenous and exogenous stress that result in the formation of DNA damage, which must be effectively repaired to maintain genome stability. Single-strand breaks (SSBs) are among the most prevalent forms of DNA damage that arise via the oxidation-induced disintegration of the sugar-phosphate backbone or as repair intermediates during base excision repair. DNA ligase IIIα (LigIIIα) is one of the primary enzymes responsible for repairing SSBs containing an intact 5′-phosphate and 3′-OH (nick) during the terminal step of single-strand break repair (SSBR) and base excision repair (BER) pathways. To date, a complete mechanistic description for how LigIIIα processes nicks within chromatin remains elusive. Here, we use a combination of biochemical assays, molecular dynamics simulations, and cryogenic electron microscopy (cryo-EM) to define the molecular basis of nick ligation in the nucleosome by LigIIIα. Quantitative enzyme kinetics reveal that the LigIIIα ligation rate is highly dependent on the translational position of the nick in the nucleosome, where nicks near the nucleosome entry/exit site are ligated with moderate efficiency and nicks near the nucleosome dyad are refractory to ligation. Cryo-EM structures of LigIIIα bound to nicks at four unique translational positions in the nucleosome reveal the structural basis for this position-dependent catalytic activity, identifying that local steric constraints imposed by the histone octamer prevent LigIIIα from readily adopting a ligation-competent conformation. Further biochemical and structural analysis demonstrates that the scaffolding protein XRCC1, which forms a heterodimer with LigIIIα, does not substantially alter the ability of LigIIIα to bind or ligate nicks in the nucleosome. Together, this work provides foundational insight into the processing of nicks in the nucleosome during the terminal step of SSBR/BER.

## Introduction

Genomic DNA is organized into chromatin by a fundamental repeating structural unit known as the nucleosome core particle (NCP). The NCP is a protein-nucleic acid complex composed of an octameric assembly containing two copies each of histones H2A, H2B, H3, and H4 wrapped by ∼147 bp of nucleosomal DNA^1,2^, which functions as structural scaffold and a regulatory barrier that controls access to genomic DNA. Chromatinized genomic DNA is continuously exposed to a variety of endogenous and exogenous stressors that generate DNA damage. This DNA damage occurs ubiquitously throughout chromatinized and non-chromatinized regions of genomic DNA^3-14^ and must be faithfully repaired to maintain genomic stability. How DNA repair enzymes overcome the nucleosome barrier to repair DNA damage and maintain genome stability remains poorly understood.

Single-strand breaks (SSBs) are among the most prevalent forms of genomic DNA damage^15,16^, and unrepaired SSBs result in mutagenesis, genome instability, and/or cell death^16-19^. SSBs arise from multiple independent mechanisms and are repaired via distinct pathways^16^. One common mechanism for generating SSBs is through oxidation-induced disintegration of the sugar-phosphate backbone, and these SSBs are repaired via the single-strand break repair (SSBR) pathway. During SSBR, the ends of the SSB are initially processed by one or more end processing enzymes (e.g., APE1, PNKP, TDP1, TDP2, APTX, FEN1) that generate a ligatable SSB containing an intact 5′-phosphate and 3′-OH (nick)^19-21^. In addition, SSBs can also arise indirectly as intermediates during the base excision repair (BER) of oxidative and alkylative DNA base damage through the combined actions of a DNA glycosylase, APE1, and DNA Polymerase β, which also generates a ligatable nick containing an intact 5′-phosphate and 3′-OH^22-24^. Though the upstream processing steps differ between SSBR and BER, repair is ultimately completed through a common terminal step where the nick intermediate containing a 5′-phosphate and 3′-OH is ligated by DNA Ligase I or a heterodimeric complex of X-ray repair cross complementing protein 1 (XRCC1) and DNA Ligase IIIα (LigIIIα)^25^.

LigIIIα is one of the primary enzymes responsible for sealing the nick intermediate containing a 5′-phosphate and 3′-OH to complete BER and SSBR^25,26^. Similar to the other human DNA ligases^27,28^, LigIIIα possesses a catalytic core made up of three domains including a DNA binding domain (DBD), a nucleotidyl transferase (NTase) domain, and a oligonucleotide/oligosaccharide-fold binding domain (OBD) that facilitate nick recognition and ligation^25,29^. In addition, LigIIIα uniquely contains a N-terminal zinc finger (ZnF) motif that contributes to DNA binding^29-34^, as well as a C-terminal BRCT domain that mediates complex formation with XRCC1^35-37^. Over the past several decades, mechanistic insight into how LigIIIα binds and ligates nicks within non-chromatinized DNA has come from robust structural and biochemical studies that identified a multi-step mechanism for nick ligation^28,29,33,38^. LigIIIα initially catalyzes the transfer of AMP from ATP to an active-site site side chain (K421), generating an adenylylated LigIIIα intermediate. The adenylylated LigIIIα then engages the Nick-DNA substrate, which is mediated by encircling of the duplex DNA by the DBD, NTase, and OBD of the catalytic core. Following engagement of the Nick-DNA substrate, LigIIIα catalyzes the transfer of AMP to the 5′-phosphate of the nick, generating an adenylylated DNA intermediate. In the final step, the 3′-OH of the nick performs nucleophilic attack on the adenylylated 5′-phosphate, resulting in phosphodiester bond formation and release of AMP. Though these prior studies have provided comprehensive molecular insight into how LigIIIα catalyzes nick ligation within non-chromatinized DNA, how LigIIIα catalyzes nick ligation within chromatin remains poorly understood. Prior *in vitro* studies established that the ligation efficiency of LigIIIα is partially inhibited by nucleosome structure^39-43^, suggesting that nick ligation may represent the rate limiting step of chromatin-based SSBR/BER^39,43^. Despite these initial observations, the molecular mechanism underlying the reduced ligation efficiency of LigIIIα in the nucleosome remains unknown. Here, we use a combination of biochemical assays, molecular dynamics simulations, and cryogenic electron microscopy (cryo-EM) to define the molecular basis of nick ligation in the nucleosome by LigIIIα.

## Results

To explore the molecular basis of nick ligation in the nucleosome by LigIIIα, we initially designed four 601 strong positioning DNA sequences containing a single ligatable nick with a 5′-phosphate and 3′-OH^44^. Each nick is located in a solvent-exposed rotational orientation at SHL−6, SHL−4, SHL−2, and SHL0 (Fig. 1a,b), herein referred to as Nick-NCP−6, Nick-NCP−4, Nick-NCP−2, and Nick-NCP0, respectively. These nick positions were chosen as solvent-exposed locations in the nucleosome are more prone to direct single-strand breaks^45,46^. In addition, nicks arising as BER intermediates also have a strong propensity to form at solvent-exposed positions in the nucleosome, as upstream BER enzymes (i.e., DNA glycosylases, APE1, and DNA Polymerase β) have robust preferences for processing solvent-exposed nucleosomal DNA damage^46-50^. Each Nick-NCP was then reconstituted using recombinant human histones (Supplementary Fig. 1a,b), and the nucleosome substrates used to dissect the molecular basis of nick recognition and ligation in the nucleosome by LigIIIα.

**Fig. 1:**
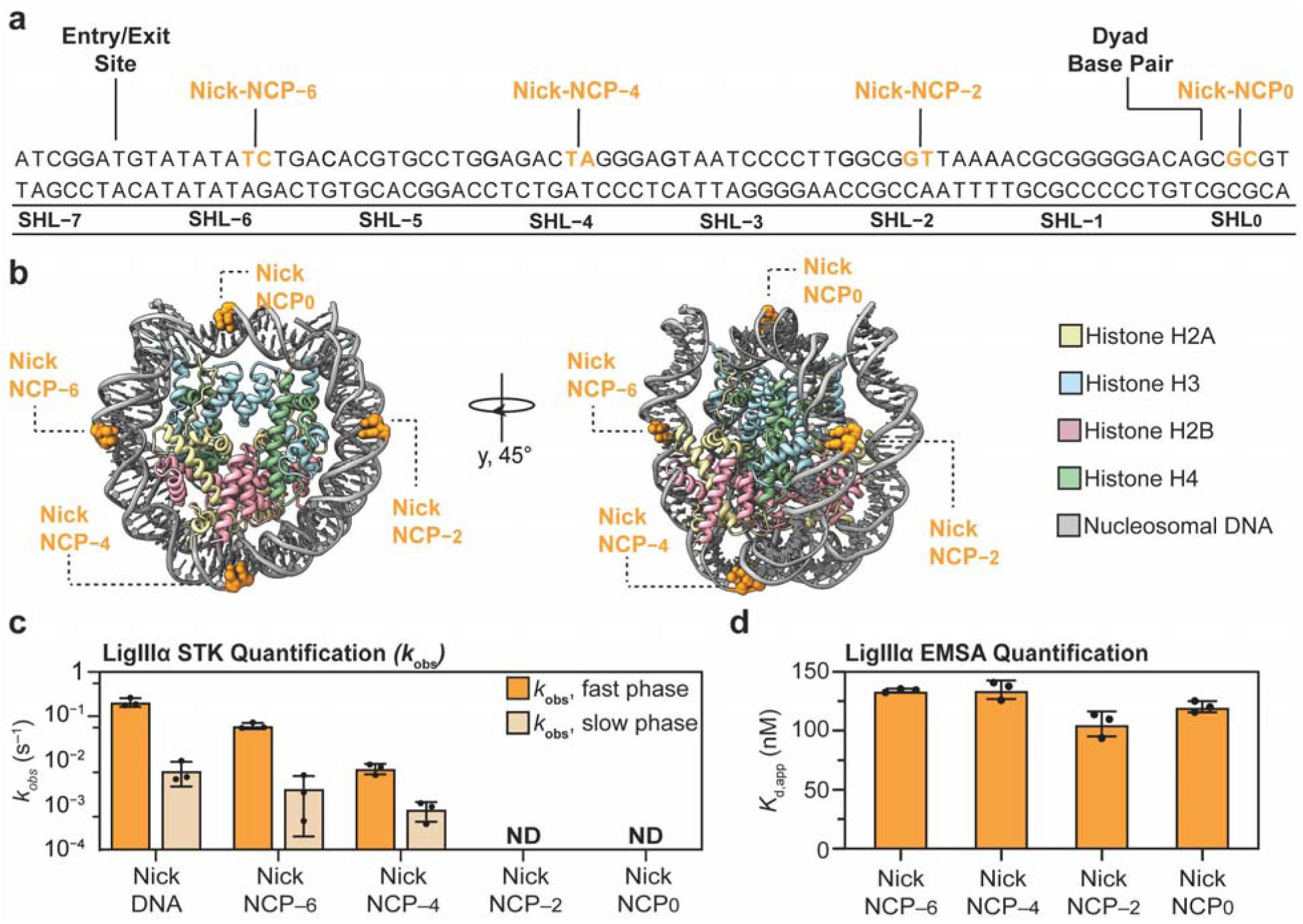
LigIIIα preferentially ligates nicks at the nucleosome entry/exit site. **a,** Diagram of the 601 strong positioning sequence with the location of each individual nick highlighted in orange. **b,** Nucleosome core particle (adapted from PDB:9DWF) shown in two orientations highlighting the nick locations at SHL−6, SHL−4, SHL−2, and SHL0. **c,** LigIIIα ligation rates (*k*_obs_) for Nick-DNA, Nick-NCP−6, Nick-NCP−4, Nick-NCP−2, and Nick-NCP0 obtained from the single-turnover kinetic analysis. Ligation rates were not determined (ND) for Nick-NCP−2 and Nick-NCP0 due to minimal product formation (>10%) during the kinetic time course. The data points represent the mean ± standard error of the mean from three independent replicate experiments. See Supplementary Fig. 2 for associated data. **d,** Apparent binding affinities (*K*_d,app_) for the interaction of LigIIIα with Nick-NCP−6, Nick-NCP−4, Nick-NCP−2, and Nick-NCP0 obtained from the EMSA analysis. The data points represent the mean ± standard deviation from three independent replicate experiments. See Supplementary Fig. 3 for associated data. All source data in this figure are provided as a Source data file.

### LigIIIα preferentially ligates nicks at the nucleosome entry/exit site

To determine the ligation kinetics of LigIIIα in the nucleosome, we utilized single-turnover (pre-steady-state) conditions to monitor the ability of LigIIIα to convert a nicked substrate to a ligated product with an intact phosphate backbone in the nucleosome. Under these conditions, the concentration of LigIIIα is in vast excess of the concentration of each Nick-NCP, ensuring the kinetic time course represents a single catalytic turnover. The resulting kinetic time course is fit to an exponential to determine the rate constant *k_obs_*, which represents the apparent rate limiting step(s) of the reaction up to and including catalysis. Of note, the positioning of the 6-fluorescein label on the nucleosomal DNA precludes the ability to distinguish the adenylyl transfer from the nick sealing step of the ligation reaction^38^. Therefore, the *k_obs_* reported here reflects the overall ligation rate within the nucleosome. To determine how the nucleosome impacts the ligation kinetics of LigIIIα, we performed the single-turnover kinetic analysis on the Nick-NCP−6, Nick-NCP−4, Nick-NCP−2, and Nick-NCP0 substrates, and compared the ligation rates to those obtained with a 50 bp Nick-DNA substrate (Fig. 1c, Supplementary Fig. 2, and Supplementary Table 1). The single-turnover kinetic analysis on the 50 bp Nick-DNA substrate revealed biphasic kinetics, consistent with two observable LigIIIα ligation phases with *k_obs_*values of 0.21 ± 0.027 s^−1^ (fast phase) and 6.1×10^−3^ ± 2.0×10^−3^ s^−1^ (slow phase). Importantly, the ligation rate of the fast phase is almost identical to the ligation rate previously determine for LigIIIα and a 28 bp Nick-DNA substrate under similar buffer conditions^38^. Similar to the 50 bp Nick-DNA substrate, biphasic behavior was also observed for Nick-NCP−6 and Nick-NCP−4, consistent with two observable LigIIIα ligation phases at both positions in the nucleosomal DNA (Fig. 1c). The ligation rates (*k*_obs_) of LigIIIα for Nick-NCP−6 were 0.061 ± 5.6×10^−3^ s^−1^ (fast phase) and 2.4×10^−3^ ± 1.3×10^−3^ s^−1^ (slow phase), which represent a 3.4-fold and 2.5-fold decrease in ligation rate compared to the 50 bp Nick-DNA control, respectively. The ligation rates (*k*_obs_) of LigIIIα for Nick-NCP−4 were 6.8×10^−3^ ± 1.8×10^−3^ s^−1^ (fast phase) and 8.1×10^−4^ ± 2.2×10^−4^ s^−1^ (slow phase), which represent a 31-fold and 7.5-fold decrease in ligation rate compared to the 50 bp Nick-DNA control, respectively. In contrast to Nick-NCP−6 and Nick-NCP−4, we observed minimal product formation during the kinetic time course for Nick-NCP−2 and Nick-NCP0 (Fig. 1c and Supplementary Fig. 2), indicating that LigIIIα exhibits severely impaired ligation kinetics for solvent-exposed nicks at translational positions in proximity to the nucleosome dyad. Together, these results demonstrate that LigIIIα preferentially ligates nicks at nucleosome entry/exit site^43^, albeit at reduced rates compared to non-nucleosomal DNA.

To exclude the possibility that the position-dependent ligation rates of LigIIIα are the result of a differential ability to engage nicks at the unique translational positions in the nucleosome, we performed electrophoretic mobility shift assays (EMSAs) to determine the apparent binding affinity (*K*_d,app_) of LigIIIα for the four Nick-NCPs (Fig. 1d, Supplementary Fig. 3, and Supplementary Table 1). Importantly, the EMSAs were performed in the absence of Mg^2+^ to prevent adenylyl transfer and ligation. The EMSA analysis revealed LigIIIα binds Nick-NCP−6 with a *K*_d,app_ of 120 ± 5 nM, Nick-NCP−4 with a *K*_d,app_ of 135 ± 8 nM, Nick-NCP−2 with a *K*_d,app_ of 106 ± 11 nM, and Nick-NCP0 with a *K*_d,app_ of 134 ± 2 nM, indicating that LigIIIα has a similar apparent binding affinity for each Nick-NCP. Notably, we observed apparent cooperativity for the interaction of LigIIIα with each Nick-NCP, with Hill coefficients ranging from 1.5 to 3 (Supplementary Fig. 3 and Supplementary Table 1). Although the precise mechanism underlying this cooperativity is unclear, it may result from the ability of LigIIIα to simultaneously engage the blunt DNA ends of the nucleosomal DNA^33,51^ and/or due to the ability of LigIIIα to cooperatively engage nicks using the ZnF and the catalytic core^33^. Ultimately, the similar apparent binding affinities for LigIIIα and the four Nick-NCPs strongly suggest that the position-dependent differences in ligation kinetics are not attributable to differences in substrate binding affinity.

### Nicks have minimal impact on nucleosome stability, structure, and dynamics

One plausible explanation for the position-dependent ligation kinetics of LigIIIα in the nucleosome is that the unique translational position of each nick may differentially impact nucleosome stability, structure, and/or dynamics^52,53^. To evaluate if the unique translational position of each nick differentially impacts nucleosome stability, we performed nucleosome stability assays on each Nick-NCP by monitoring the conversion of intact nucleosome to free DNA following incubation at 37 °C for 24 hours (Fig 2a,b and Supplementary Fig. 12a)^54^. After the 24 hour-incubation, we observed 87 ± 3%, 82 ± 3%, 87 ± 3%, and 78 ± 6% of intact nucleosome remaining for Nick-NCP−6, Nick-NCP−4, Nick-NCP−2, and Nick-NCP0, respectively, which is similar to the 85 ± 4% of intact nucleosome remaining for a non-damaged NCP (ND-NCP). This data indicates that the solvent-exposed nicks at each unique translational position in the nucleosomal DNA have minimal impact on overall nucleosome stability.

**Fig. 2:**
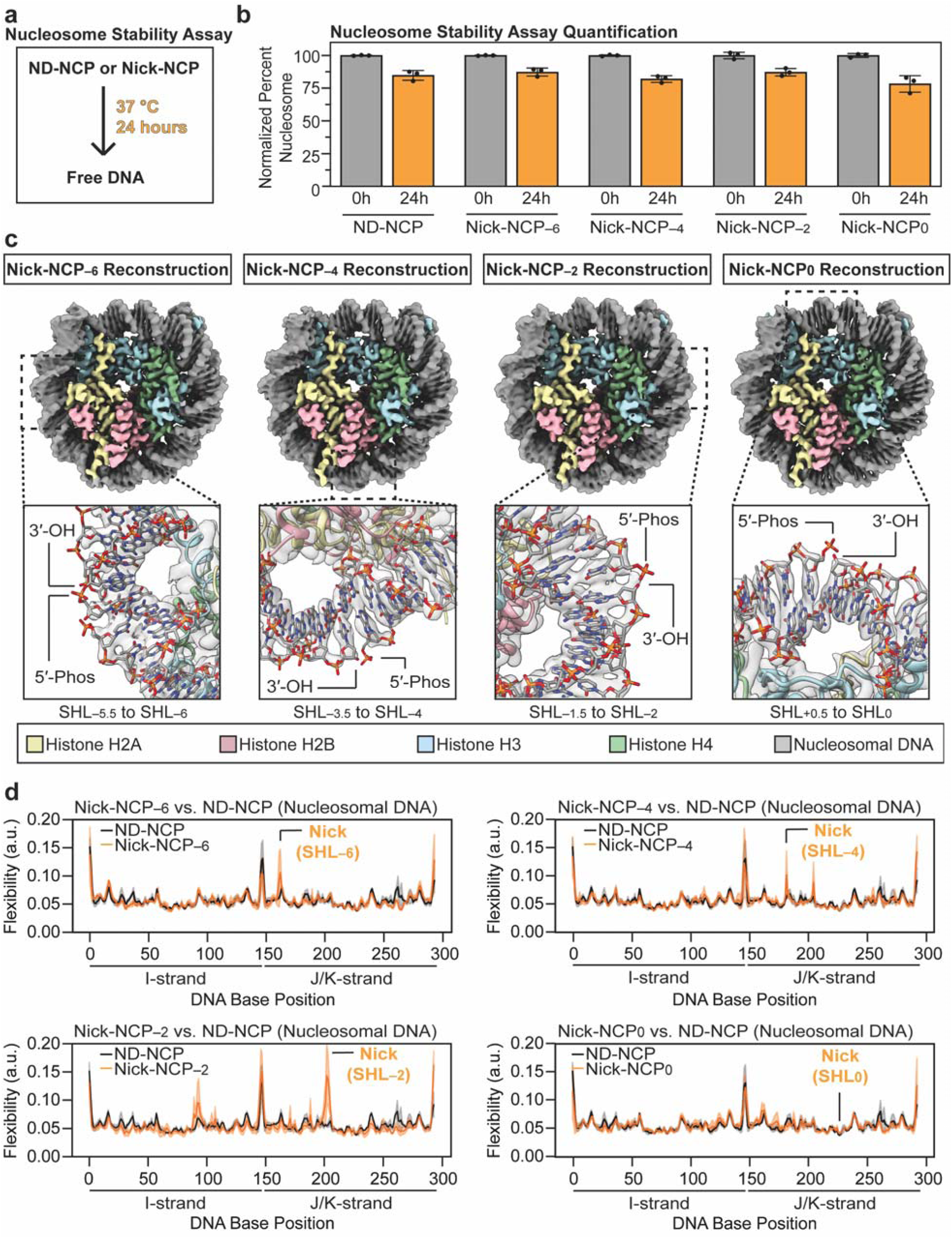
Nicks have minimal impact on nucleosome stability, structure, and dynamics. **a,** Diagram outlining the nucleosome stability assays. **b,** Quantification of the nucleosome stability assays for ND-NCP, Nick-NCP−6, Nick-NCP−4, Nick-NCP−2, and Nick-NCP0. The data points represent the mean ± standard deviation from three independent replicate experiments. **c,** Cryo-EM reconstructions of Nick-NCP−6 (left), Nick-NCP−4 (middle left), Nick-NCP−2 (middle right), and Nick-NCP0 (right) with a with a focused view of the nick site shown as an inset. Cryo-EM density for the nucleosomal DNA and histones are shown as a transparent gray surface. **d,** PCA-based flexibility analysis comparing the dynamics of the nucleosomal DNA in the ND-NCP and Nick-NCP−6 (top left), Nick-NCP−4 (top right), Nick-NCP−2 (bottom left), and Nick-NCP0 (bottom right) trajectories. The I-strand (non-damaged) and J/K-strands (nicked strands) are represented by DNA base positions 1-147 and 148-295, respectively. The average flexibility and standard deviation from the triplicate ND-NCP trajectories are represented by a black and gray line, respectively. The average flexibility and standard deviation from the triplicate Nick-NCP trajectories are represented by an orange and light orange line, respectively. All source data in this figure are provided as a Source data file.

To determine if the unique translational position of each nick differentially impacts nucleosome structure, we obtained cryo-EM reconstructions of the ND-NCP, Nick-NCP−6, Nick-NCP−4, Nick-NCP−2, and Nick-NCP0 at overall resolutions of 2.6 Å, 2.8 Å, 2.7 Å, 2.9 Å, and 2.8 Å, respectively (Fig. 2c and Supplementary Figs. 4-11). Structural comparison of each Nick-NCP to the ND-NCP structure revealed an overall RMSD of 0.15 Å for Nick-NCP−6, 0.15 Å for Nick-NCP−4, 0.16 Å for Nick-NCP−2, and 0.10 Å for Nick-NCP0 (Supplementary Fig. 12b), indicating that nicks do not substantially alter the conformation of the core histone octamer or the nucleosomal DNA. Of note, we did observe subtle changes within the nucleosomal DNA at each nick site (Supplementary Fig. 12c), consistent with localized fraying around the nick site. However, these localized changes at the nick site were observed at all four translational positions, suggesting that localized DNA fraying cannot explain the position-dependent ligation kinetics of LigIIIα in the nucleosome. Together, this structural analysis indicates that solvent-exposed nicks at unique translational positions in the nucleosomal DNA do not substantially alter nucleosome structure.

To probe whether the unique translational position of each nick differentially impacts nucleosome dynamics, we performed 1 µs MD simulations on the ND-NCP, Nick-NCP−6, Nick-NCP−4, Nick-NCP−2, and Nick-NCP0. These simulations provide direct insight into dynamics of damaged nucleosomes on the microsecond timescale (e.g., DNA fraying)^52,55,56^, but the relatively short timescale of the MD simulations precludes the ability to probe how DNA lesions impact nucleosome dynamics that occur on longer timescales (e.g., unwrapping of the nucleosomal DNA at the entry/exit site)^57-59^. Initial inspection of the MD simulations confirmed the overall stability of the nucleosome containing nicks at each unique translation position throughout the trajectories, as only subtle differences in the overall conformation of the histone octamer and the nucleosomal DNA were observed (Supplementary Fig. 13a). However, these subtle differences include elevated DNA dynamics at the nick sites for Nick-NCP−6, Nick-NCP−4, and Nick-NCP−2 (Fig. 2d), which correspond to localized fraying events that separate the 5′-phosphate and 3′-OH (Supplementary Fig. 13b). Of note, these elevated DNA dynamics at the nick site were not observed for Nick-NCP0 (Fig. 2d and Supplementary Fig. 13b), consistent with the enhanced stability of the nucleosomal DNA adjacent to the nucleosome dyad. Despite these observations, the elevated DNA dynamics at the nick site for Nick-NCP−2, Nick-NCP−4, and Nick-NCP−6 do not correlate with the position-dependent differences in LigIIIα ligation kinetics in the nucleosome (Fig. 1c). Together, the MD simulations indicate that nicks have minimal impact on overall nucleosome structure, and only a subtle impact on microsecond timescale nucleosome dynamics at the nick site (i.e., nick fraying). Collectively, these data indicate that the unique translational position of each nick has minimal impact on nucleosome stability, structure, and fast timescale dynamics, and therefore cannot fully account for the position-dependent ligation kinetics of LigIIIα in the nucleosome.

### Structural basis of nick ligation in the nucleosome by LigIIIα

To further probe the mechanistic basis for the position-dependent LigIIIα ligation kinetics in the nucleosome, we obtained cryo-EM reconstructions of LigIIIα bound to the four NCPs containing a nick at SHL−6, SHL−4, SHL−2, and SHL0 (Fig. 3). This was accomplished by generating complexes of adenylylated LigIIIα bound to Nick-NCP−6, Nick-NCP−4, Nick-NCP−2, and Nick-NCP0 that were stabilized via glutaraldehyde crosslinking. Of note, these complexes were formed in the absence of Mg^2+^ to ensure a pre-catalytic conformation was captured prior to adenylyl transfer and ligation. The complexes were then subjected to single particle analysis, which resulted in cryo-EM reconstructions of the LigIIIα-Nick-NCP−6, LigIIIα-Nick-NCP−4, LigIIIα-Nick-NCP−2, and LigIIIα-Nick-NCP0 complexes at overall resolutions of 2.8 Å, 2.9 Å, 3.0 Å, and 2.5 Å, respectively (Supplementary Figs. 5-8, 14-17 and Supplementary Tables 2-4). Importantly, the local resolution of LigIIIα in the LigIIIα-Nick-NCP−6 and LigIIIα-Nick-NCP0 reconstructions were of sufficient quality to dock a previously determined X-ray crystal structure of LigIIIα bound to non-nucleosomal Nick-DNA^29^ and manually build most amino acid side chains throughout the LigIIIα DNA binding interface and active site (Supplementary Fig. 14,17). In contrast, the local resolution of LigIIIα in the LigIIIα-Nick-NCP−4 and LigIIIα-Nick-NCP−2 reconstructions was not sufficient to observe amino acid side chains throughout most of the LigIIIα DNA binding interface and active site (Supplementary Fig. 15,16). Therefore, the LigIIIα side chain residues within these structures generally reflect their position within the higher resolution LigIIIα-Nick-NCP0 structure.

**Fig. 3:**
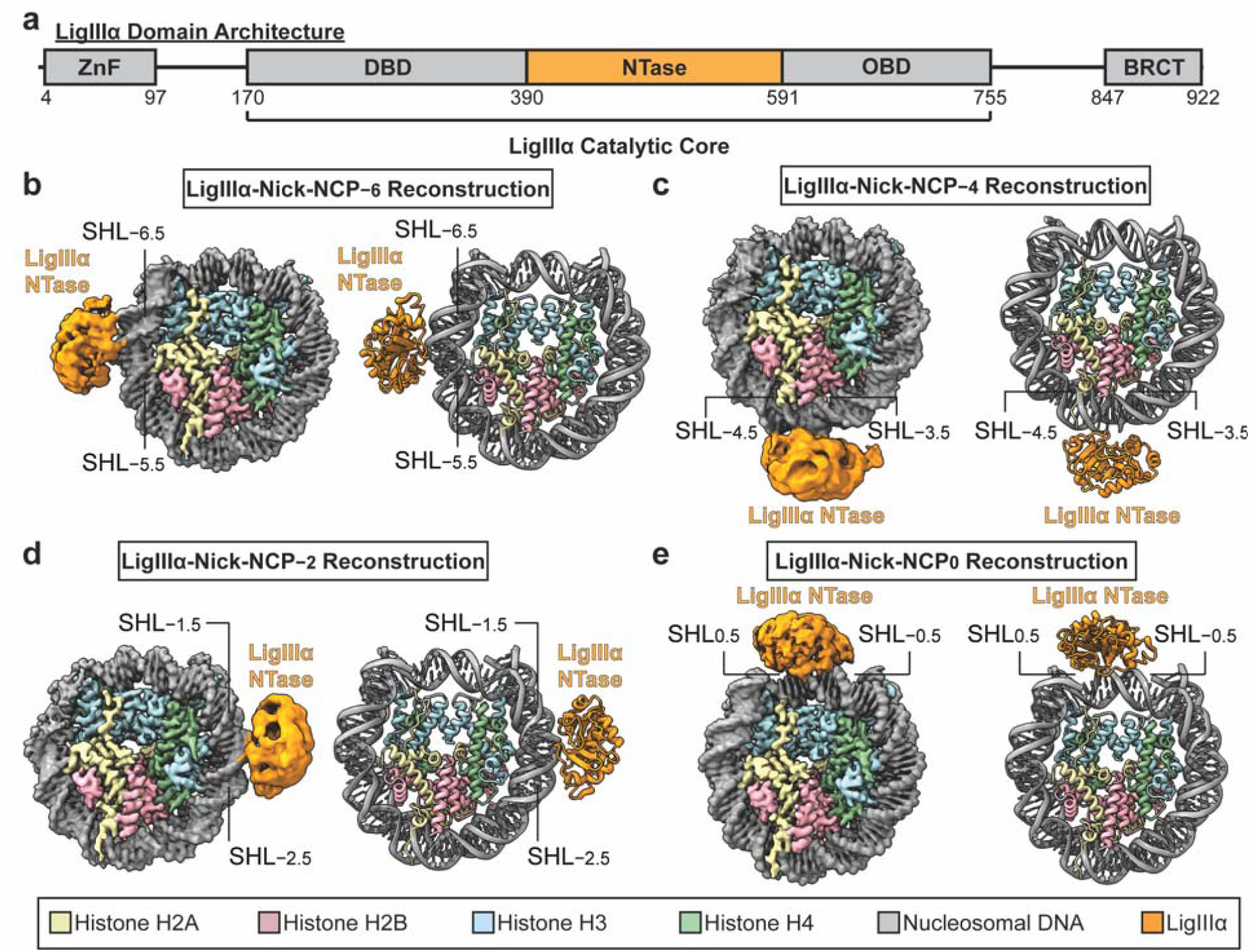
Cryo-EM reconstructions of LigIIIα-Nick-NCP complexes. **a,** Diagram showing the domain architecture of LigIIIα. **b,** Composite cryo-EM map and model of the LigIIIα-Nick-NCP−6 complex. **c,** Composite cryo-EM map and model of the LigIIIα-Nick-NCP−4 complex. **d,** Composite cryo-EM map and model of the LigIIIα-Nick-NCP−2 complex. **e,** Cryo-EM map and model of the LigIIIα-Nick-NCP0 complex.

Prior structural studies of LigIIIα bound to non-nucleosomal Nick-DNA identified that nick recognition is mediated by the coordinated action of the DNA-binding domain (DBD), nucleotidyltransferase domain (NTase), and oligosaccharide-fold binding domain (OBD), which completely encircle the DNA to engage the nick^29,33^. Interestingly, of the three domains that contribute to nick recognition in non-nucleosomal DNA (DBD, NTase, and OBD), we only observed clear density for the LigIIIα NTase domain in the reconstructions of LigIIIα bound to a nick in the nucleosome at SHL−6, SHL−4, SHL−2, and SHL0 (Fig. 3). Closer inspection of each reconstruction after applying a low-pass filter (8 Å) revealed additional density adjacent to the LigIIIα NTase (Supplementary Fig. 18), though the substantial structural heterogeneity precludes the ability to assign this density to other domains within the catalytic core (i.e., the DBD or OBD), the ZnF, or the BRCT domain.

The structures of LigIIIα bound to nicks in the nucleosome at SHL−6, SHL−4, SHL−2, and SHL0 provide molecular insight into nick recognition in the nucleosome. In all four structures, the LigIIIα NTase engages the nucleosomal DNA with an ∼6 bp footprint that encompasses the nick site (Fig. 4). Importantly, LigIIIα only interacts with the nucleosomal DNA and does not make discernible contacts with the histone octamer core. The interaction between the LigIIIα and the nucleosome is primarily driven by non-specific electrostatic interactions with the phosphate backbone of the damaged nucleosomal DNA strand (Fig. 4). On the 3′ side, Lys444, His449, and Lys450 coordinate the phosphate backbone of 4 bp of nucleosomal DNA. On the 5′ side of the nick, Arg441, Arg584, and Lys588 make direct contacts with the 5′-phosphate. Notably, structural comparison of the LigIIIα DNA binding interfaces at SHL−6, SHL−4, SHL−2, and SHL0 reveals a consistent mode of DNA binding at each unique translational position (Supplementary Fig. 19), indicating that LigIIIα recognizes nicks at different translational positions in the nucleosome using a similar general mechanism.

**Fig. 4:**
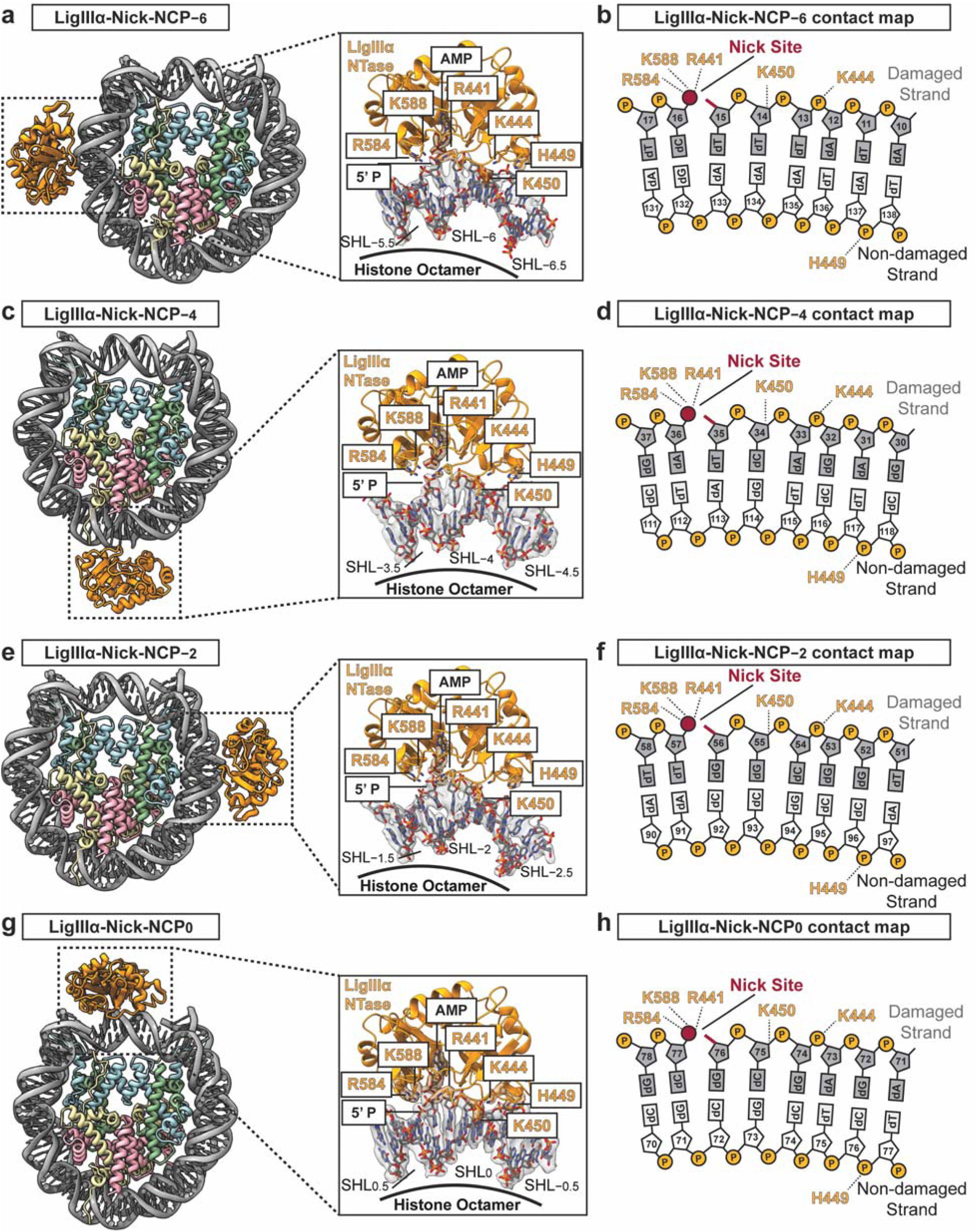
The LigIIIα NTase domain mediates nick recognition in the nucleosome. **a,** A model of the LigIIIα-Nick-NCP−6 complex with a focused view of the LigIIIα-nucleosomal DNA binding interface from SHL−5.5 to SHL−6.5 shown as an inset. Segmented cryo-EM density for LigIIIα and the nucleosomal DNA are shown as a transparent gray surface. **b,** Contact map showing the interaction between LigIIIα and the nucleosomal DNA at SHL−6. LigIIIα residues that interact with the nucleosomal DNA are labeled in orange. **c,** A model of the LigIIIα-Nick-NCP−4 complex with a focused view of the LigIIIα-nucleosomal DNA binding interface from SHL−3.5 to SHL−4.5 shown as an inset. Segmented cryo-EM density for LigIIIα and the nucleosomal DNA are shown as a transparent gray surface. **d,** Contact map showing the interaction between LigIIIα and the nucleosomal DNA at SHL−4. LigIIIα residues that interact with the nucleosomal DNA are labeled in orange. **e,** A model of the LigIIIα-Nick-NCP−2 complex with a focused view of the LigIIIα-nucleosomal DNA binding interface from SHL−1.5 to SHL−2.5 shown as an inset. Segmented cryo-EM density for LigIIIα and the nucleosomal DNA are shown as a transparent gray surface. **f,** Contact map showing the interaction between LigIIIα and the nucleosomal DNA at SHL−2. LigIIIα residues that interact with the nucleosomal DNA are labeled in orange. **g,** A model of the LigIIIα-Nick-NCP0 complex with a focused view of the LigIIIα-nucleosomal DNA binding interface from SHL−0.5 to SHL0.5 shown as an inset. Segmented cryo-EM density for LigIIIα and the nucleosomal DNA are shown as a transparent gray surface. **h,** Contact map showing the interaction between LigIIIα and the nucleosomal DNA at SHL0. LigIIIα residues that interact with the nucleosomal DNA are labeled in orange.

Prior structural studies of LigIIIα bound to non-nucleosomal Nick-DNA established the structural basis of nick ligation^29,33^. To catalyze nick ligation in non-nucleosomal Nick-DNA, the LigIIIα NTase, DBD, and OBD completely encircle the DNA duplex and introduce a sharp kink at the nick site (Supplementary Fig. 20a). This ultimately transitions the DNA on the 3′ end of the nick from B-form to A-form and shifts the 3′ nucleotide from a C2′ endo conformation to a C3′ endo conformation, which properly organizes the nick within the active site for catalysis (Supplementary Fig. 20a). Importantly, our inability to resolve the DBD and OBD in the reconstructions of LigIIIα bound to nicks at SHL−6, SHL−4, SHL−2, and SHL0 indicates that these domains are unable to encircle the nucleosomal DNA in a stable manner (Fig. 3), suggesting that these structures may represent a conformation of LigIIIα that is incompatible with nick ligation. To further explore this possibility, we compared the structures of LigIIIα bound to nicks in the nucleosome at SHL−6, SHL−4, SHL−2, and SHL0 to the apo Nick-NCP structures (Supplementary Fig. 20b). This structural comparison revealed that LigIIIα induces subtle rearrangements in the nucleosomal DNA upon nick recognition at each of the four translational positions, which are largely localized to the 5′-phosphate and involve a modest ∼5 Å to ∼7 Å movement of the 5′-phosphate toward the active site (Supplementary Fig. 20b). Despite clear stabilization of the 5′-phosphate within the NTase domain, the nucleosomal DNA on the 3′ end of the nick remains in B-form and the 3′ nucleotide remains in the C2′-endo conformation (Fig. 5a-d and Supplementary Fig. 20f-i). The inability of LigIIIα to encircle the nucleosomal DNA, transition the nucleosomal DNA on the 3′ end of the nick to A-form, and shift the 3′ nucleotide into a C3′ endo conformation indicate these structures represent LigIIIα in a conformation that is incompatible with nick ligation. Importantly, these observations provide a structural rationale for the severely reduced ligation kinetics at SHL0 and SHL−2 and the moderate ligation kinetics observed at SHL−4 and SHL−6 (Fig. 1c). At all four nick locations in the nucleosome, encircling and kinking the nucleosomal DNA for ligation would generate substantial clashes between the LigIIIα OBD/DBD and the core histone octamer (Fig. 5e-h). However, the nucleosomal DNA at the entry/exit site is prone to spontaneous unwrapping from the histone octamer^58-60^, which could provide an opportunity for LigIIIα to encircle and deform the nucleosomal DNA for ligation at SHL−6 and SHL−4 without steric conflict with the histone octamer (see discussion). Together, this structural analysis indicates that LigIIIα is readily able to engage nicks at multiple translational positions within the nucleosomal DNA but is unable to efficiently encircle and deform the nucleosomal DNA to properly organize the active site for nick ligation when the nucleosomal DNA remains engaged with the histone octamer.

**Fig. 5:**
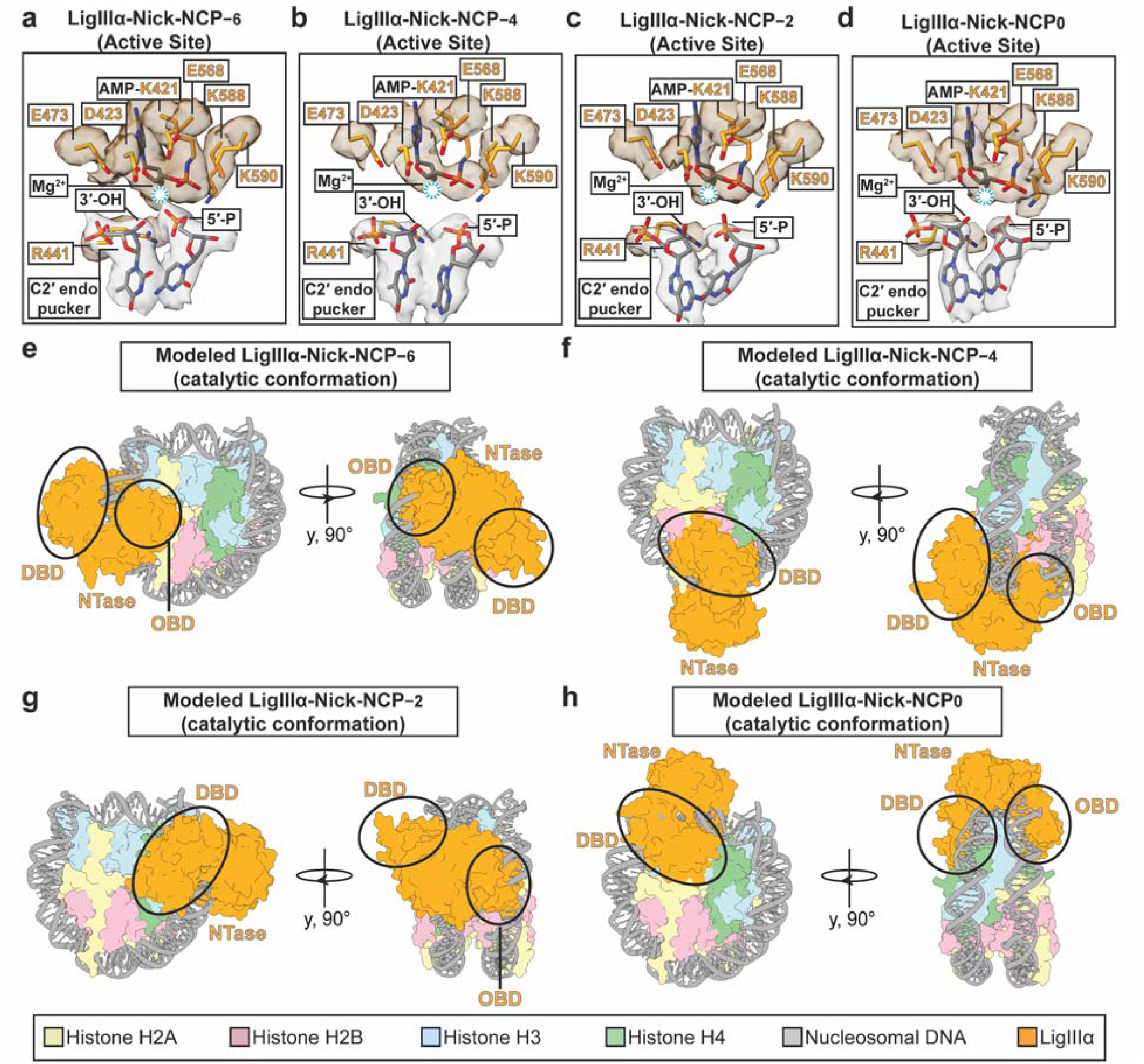
LigIIIα adopts a catalytically inactive conformation during nick recognition in the nucleosome. **a-d,** Focused views of the LigIIIα active site in the LigIIIα-Nick-NCP−6, LigIIIα-Nick-NCP−4, LigIIIα-Nick-NCP−2, and LigIIIα-Nick-NCP0 complexes. The segmented cryo-EM density for LigIIIα active site residues and the nucleosomal DNA are shown as a transparent yellow or gray surface, respectively. **e-g,** Structural models of LigIIIα bound to Nick-NCP−6, Nick-NCP−4, Nick-NCP−2, and Nick-NCP0 in a catalytic conformation. Substantial clashes between the LigIIIα OBD/DBD and the nucleosome are circled in black.

### XRCC1 has minimal impact on LigIIIα nick binding and ligation in the nucleosome

LigIIIα forms a heterodimer with the scaffolding protein XRCC1^61-64^, which is important for LigIIIα stability^62,65-67^ and recruitment of LigIIIα to sites of DNA damage in cellular chromatin^68,69^. We reasoned that XRCC1 may stimulate the ability of LigIIIα to bind and/or ligate nicks in the nucleosome. To test this possibility, we generated an XRCC1-LigIIIα complex from recombinant XRCC1 and adenylylated LigIIIα purified from *E. coli* (Supplementary Fig. 1c). We then performed single-turnover kinetic analysis with the XRCC1-LigIIIα complex and a 50 bp Nick-DNA, Nick-NCP−6, Nick-NCP−4, Nick-NCP−2, and Nick-NCP0 substrates to determine whether XRCC1 could stimulate the ligation efficiency of LigIIIα in the nucleosome (Fig. 6b, Supplementary Fig. 21, and Supplementary Table 1). Similar to the analysis with LigIIIα alone, the single-turnover kinetics of the XRCC1-LigIIIα complex on Nick-NCP−6 and Nick-NCP−4 revealed biphasic behavior indicating two kinetic phases of ligation at both positions in the nucleosomal DNA (Fig. 6b). The ligation rates (*k*_obs_) of XRCC1-LigIIIα on Nick-NCP−6 were 0.034 ± 1.9x10^−3^ s^−1^ (fast phase) and 2.1x10^−3^ ± 1.1x10^−3^ s^−1^ (slow phase), and the ligation rates (*k*_obs_) of XRCC1-LigIIIα on Nick-NCP−4 were 7.5x10^−3^ ± 0.8x10^−3^ s^−1^ (fast phase) and 6.9x10^−4^ ± 2.1x10^−4^ s^−1^ (slow phase). Importantly, these ligation rates for the XRCC1-LigIIIα complex on Nick-NCP−6 and Nick-NCP−4 are strikingly similar to those determined for LigIIIα alone (compare Fig. 1c and Fig. 6b), indicating that XRCC1 does not substantially enhance LigIIIα nick ligation at translational positions near the nucleosome entry/exit site. Notably, we also observed minimal product formation during the XRCC1-LigIIIα kinetic time course on Nick-NCP−2 and Nick-NCP0 (Fig. 6b and Supplementary Fig. 21), consistent with the observations for LigIIIα alone (Fig. 1c and Supplementary Fig. 2), indicating that XRCC1 does not substantially enhance LigIIIα nick ligation at translational positions in proximity to the nucleosome dyad.

**Fig. 6:**
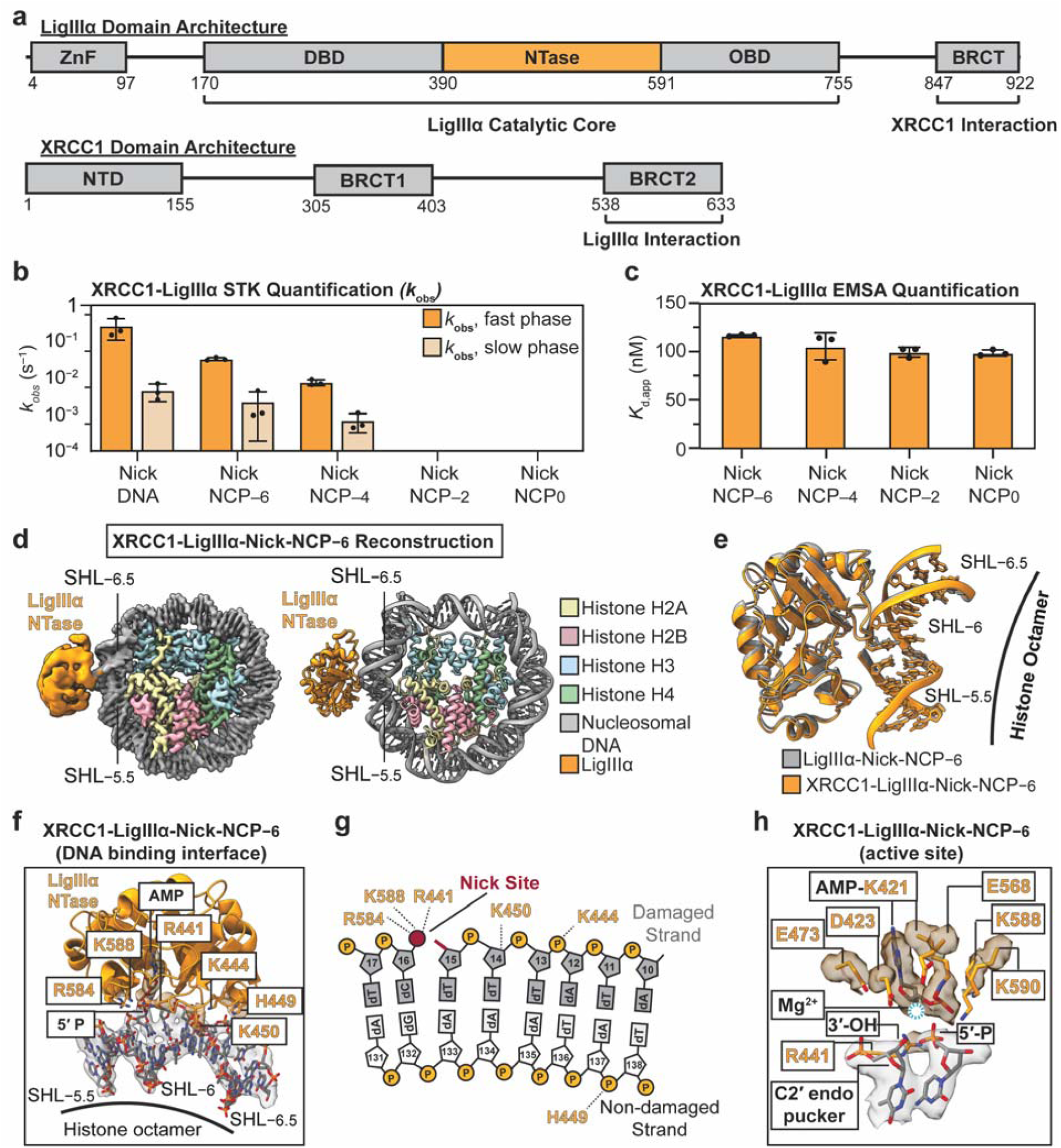
XRCC1 has minimal impact on LigIIIα nick binding and ligation in the nucleosome. **a,** Diagram showing the domain architecture of LigIIIα and XRCC1. **b,** XRCC1-LigIIIα ligation rates (*k*_obs_) for Nick-DNA, Nick-NCP−6, Nick-NCP−4, Nick-NCP−2, and Nick-NCP0 obtained from the single-turnover kinetic analysis. Ligation rates were not determined (ND) for Nick-NCP−2 and Nick-NCP0 due to minimal product formation (>10%) during the kinetic time course. The data points represent the mean ± standard error of the mean from three independent replicate experiments. See Supplementary Fig. 21 for associated data. **c,** Apparent binding affinities (*K*_d,app_) for the interaction of XRCC1-LigIIIα with Nick-NCP−6, Nick-NCP−4, Nick-NCP−2, and Nick-NCP0 obtained from the EMSA analysis. The data points represent the mean ± standard deviation from three independent replicate experiments. See Supplementary Fig. 22 for associated data. **d,** Composite cryo-EM map and model of the XRCC1-LigIIIα-Nick-NCP−6 complex. **e**, Structural comparison of the LigIIIα/nucleosomal DNA binding interface in the LigIIIα-Nick-NCP−6 (gray) and LigIIIα-Nick-NCP−6 (orange) complexes. **f,** A model of the XRCC1-LigIIIα-Nick-NCP−6 complex with a focused view of the LigIIIα-nucleosomal DNA binding interface from SHL−5.5 to SHL−6.5 shown as an inset. Segmented cryo-EM density for LigIIIα and the nucleosomal DNA are shown as a transparent gray surface. **g,** Contact map showing the interaction between LigIIIα and the nucleosomal DNA at SHL−6 in the XRCC1-LigIIIα-Nick-NCP−6. **h,** Focused view of the LigIIIα active site in the XRCC1-LigIIIα-Nick-NCP−6 complex. All source data in this figure are provided as a Source data file.

To define whether XRCC1 enhances the ability of LigIIIα to recognize nicks in the nucleosome, we performed EMSAs to determine the apparent binding affinity (*K*_d,app_) of the XRCC1-LigIIIα complex for Nick-NCP−6, Nick-NCP−4, Nick-NCP−2, and Nick-NCP0 (Fig. 6c, Supplementary Fig. 22, and Supplementary Table 1). The ESMA analysis revealed XRCC1-LigIIIα binds Nick-NCP−6 with a *K*_d,app_ of 99 ± 3 nM, Nick-NCP−4 with a *K*_d,app_ of 106 ± 14 nM, Nick-NCP−2 with a *K*_d,app_ of 100 ± 5 nM, and Nick-NCP0 with a *K*_d,app_ of 125 ± 15 nM. Notably, the apparent binding affinities of the XRCC1-LigIIIα complex for the Nick-NCPs are strikingly similar to that observed for LigIIIα alone (Compare Fig. 1d and Fig. 6c), indicating that XRCC1 does not substantially enhance LigIIIα nick binding within the nucleosome.

To determine if XRCC1 alters the structural mechanism used by LigIIIα to engage nicks in the nucleosome, we obtained a cryo-EM reconstruction of the XRCC1-LigIIIα complex bound to a NCP containing a nick at SHL−6 (Fig. 6d). This was accomplished by generating a complex of adenylylated XRCC1-LigIIIα bound to Nick-NCP−6 that was stabilized via glutaraldehyde crosslinking. The complex was then subjected to single particle analysis, which resulted in a of 2.9 Å reconstruction of the XRCC1-LigIIIα-Nick-NCP−6 complex (Fig. 6d, Supplementary Fig. 23-24, and Supplementary Table 5). Similar to the LigIIIα-Nick-NCP−6 reconstruction, we only observed clear density for the LigIIIα NTase domain in the XRCC1-LigIIIα-Nick-NCP−6 reconstruction (Fig. 6d), and the remaining catalytic core of LigIIIα (OBD and DBD), the ZnF, and the BRCT domain were unresolved. We were also unable to resolve XRCC1 in the reconstruction, including the known interaction interface between the BRCT domain of LigIIIα and the BRCT2 domain of XRCC1^35-37,64,70^. Closer inspection of the XRCC1-LigIIIα-Nick-NCP−6 structure revealed several striking similarities to the LigIIIα-Nick-NCP−6 structure (Fig. 6e). In the XRCC1-LigIIIα-Nick-NCP−6 structure, the LigIIIα NTase engages the nucleosomal DNA with an ∼6 bp footprint and directly coordinates the nick using the same general mechanism observed for LigIIIα alone (Fig. 6f,g). Importantly, LigIIIα remains unable to encircle the nucleosomal DNA, transition the nucleosomal DNA on the 3′ end of the nick to A-form, nor shift the 3′ nucleotide into a C3′ endo conformation (Fig. 6h), indicating the presence of XRCC1 does not enable LigIIIα to adopt a ligation-competent conformation on the Nick-NCP. Collectively, these biochemical and structural observations suggest that XRCC1 does not play a substantial role in facilitating the ability of LigIIIα to recognize or ligate nicks in the nucleosome.

## Discussion

Our work has established the molecular basis of nick ligation in the nucleosome by LigIIIα, providing fundamental insight into the terminal step of SSBR/BER in chromatin. Our kinetic analysis revealed that the ligation kinetics of LigIIIα in the nucleosome are highly dependent on the nick position within the nucleosomal DNA. Nicks near the nucleosome entry/exit site are ligated by LigIIIα with moderate ligation rates, whereas nicks near the nucleosome dyad are refractory to ligation. Despite this strong positional preference for ligation, our nucleosome binding assays demonstrate that LigIIIα robustly binds nicks at both the nucleosome entry/exit site and the nucleosome dyad, indicating that the position-dependent differences in ligation kinetics are not the result of large differences in substrate binding affinity. Our cryo-EM structures of LigIIIα bound to solvent-exposed nicks at four distinct translational positions provide a structural rationale for these observations. These structures reveal the LigIIIα NTase domain readily engages solvent-exposed nicks at all four translational positions in the nucleosome. However, the inability of the DBD and OBD to encircle and kink the nucleosomal nick prevents LigIIIα from adopting a catalytic conformation compatible with nick ligation. While these structures reveal why LigIIIα is generally unable to efficiently ligate nicks at translational positions adjacent to the nucleosome dyad, they do not directly explain how LigIIIα catalyzes nick ligation near the nucleosome entry/exit site. We hypothesize that spontaneous unwrapping of the nucleosomal DNA at the entry/exit site^58-60^ may transiently relieve the steric constraints imposed by the histone octamer, providing an opportunity for LigIIIα to encircle and kink the nucleosomal DNA to facilitate nick ligation. Though we were unable to capture a structure of LigIIIα bound to the nucleosome in this configuration, the ligation rates determined for LigIIIα near the nucleosome entry/exit site (Supplementary Table 1) are less than the rate constant determined for spontaneous unwrapping of the nucleosomal DNA from the histone octamer at the entry/exit site^58,59^. This suggests that intrinsic nucleosome dynamics at the entry/exit site may represent the rate limiting step of LigIIIα nick ligation in the nucleosome at these locations.

Over the past several years, structural studies have defined how the SSBR/BER enzymes that act upstream of LigIIIα, including multiple DNA glycosylases^5,71,72^, APE1^73^, and DNA Polymerase β^74^, recognize and process DNA lesions in the nucleosome. These structural studies demonstrated that SSBR/BER enzymes use localized or global DNA sculpting mechanisms (i.e., enzyme-induced deformations in the nucleosomal DNA) to reposition their cognate DNA lesions into the active site for catalysis^47^. In contrast to the upstream SSBR/BER, our structures reveal that LigIIIα is unable to perform the nucleosomal DNA sculpting required for catalysis, which includes encircling the DNA and introducing a sharp kink at the nick site to properly position the nick into the active site. Instead, LigIIIα likely relies on intrinsic nucleosome dynamics to transiently expose the nick to facilitate ligation. This fundamental difference between LigIIIα and upstream SSBR/BER enzymes likely explains why nick ligation represents the rate-limiting step of BER at most positions in the nucleosome^39,43^. Of note, it is interesting to speculate whether the nucleosome barrier to ligation may be partially overcome through substrate channeling^75-79^, the process where sequential BER enzymes act in a coordinated fashion to prevent the accumulation of toxic repair intermediates^80-82^. Given the requirement that LigIIIα encircle and sharply kink the nucleosomal DNA for catalysis^29^, it is possible that the stepwise DNA sculpting induced by upstream SSBR/BER enzymes progressively displaces the nucleosomal DNA from the histone octamer to facilitate LigIIIα binding in a catalytically competent conformation^47^. While recent work has begun to probe substrate channeling between BER enzymes in the nucleosome^43,83^, future work will be needed to unambiguously determine whether substrate channeling can occur in the nucleosome and decipher the impact this coordination has on the ability of LigIIIα to catalyze nick ligation in the nucleosome.

XRCC1 is a central scaffold in the BER and SSBR pathways that forms a heterodimer with LigIIIα^61-63^, which is important for LigIIIα stability^62,65-67^ and recruitment of LigIIIα to sites of DNA damage in cellular chromatin^68,69^. Prior work identified that XRCC1 modestly stimulates the ability of LigIIIα to catalyze nick ligation in the nucleosome^40^ by partially disrupting nucleosome structure^39,40^, suggesting that XRCC1 may help LigIIIα partially overcome the nucleosome barrier to repair. In contrast to these prior observations, our biochemical assays demonstrate that XRCC1 does not substantially impact LigIIIα nick recognition or nick ligation in the nucleosome. Consistent with our biochemical observations, our structural analysis of nick recognition in the nucleosome by the XRCC1-LigIIIα complex revealed that XRCC1 does not directly engage the nucleosome, does not disrupt overall nucleosome structure, and does not promote a LigIIIα conformation compatible with catalysis. The basis of these conflicting results is unclear, though one notable difference is that the prior work utilized XRCC1 purified from insect cells^40^, whereas our study used XRCC1 purified from *E. coli* cells. It is possible that the XRCC1 purified from insect cells carries post-translational modifications^84-89^ that enable XRCC1 to modulate nucleosome structure and/or stimulate LigIIIα nick ligation in the nucleosome, which are not present in XRCC1 purified from *E. coli* cells. Nevertheless, our data supports the idea that the primary role of XRCC1 in chromatin-based SSBR/BER is to recruit LigIIIα to sites of DNA damage^68,69^ and/or to coordinate substrate channeling between upstream SSBR/BER enzymes and LigIIIα within chromatin^68,90-92^, rather than directly facilitating the ability of LigIIIα to catalyze nick ligation in the nucleosome.

Our work, in conjunction with prior biochemical studies strongly suggests that LigIIIα is incapable of efficiently catalyzing nick ligation in the nucleosome at nick positions outside of the nucleosome entry/exit site^39,40,43^. This raises the important question of how LigIIIα overcomes the nucleosome barrier to ligation at these positions in order to successfully complete chromatin-based SSBR/BER. We hypothesize that the nucleosome barrier to ligation is overcome through a combination of stochastic and DNA damage-dependent chromatin remodeling, both of which could generate a chromatin environment permissive for LigIIIα to catalyze nick ligation. Stochastic changes in chromatin structure (i.e., chromatin remodeling) that arise during nuclear processes such as transcription and DNA replication may randomly reposition nicks to more accessible locations in the nucleosome, displace nicks into linker DNA regions, or completely disassemble the histone octamer, all of which would enhance LigIIIα nick ligation. In addition to stochastic chromatin remodeling, DNA damage-dependent chromatin remodeling may also play a key role in facilitating nick ligation in the nucleosome. For example, the chromatin signaling cascade involving PARP1/2, HPF1, and downstream ATP-dependent chromatin remodeling enzymes has been implicated in facilitating BER and SSBR within cellular chromatin^93-110^. Consistent with this idea, the recruitment of XRCC1-LigIIIα to sites of DNA damage is dependent on ADP-ribosylation^69,94,98,110,111^, and recent work identified that histone ADP-ribosylation enhances XRCC1-LigIIIα nick ligation in the nucleosome by altering nucleosome dynamics^42^. In addition, the multiple ATP-dependent chromatin remodeling enzymes (e.g., ALC1) recruited downstream of PARP1/2 may function to reposition nicks to more accessible locations in the nucleosome (e.g., the nucleosome entry/exit site) and/or displace nicks into linker DNA regions that would also promote productive LigIIIα nick ligation. Our structural observation that LigIIIα can engage nicks in the nucleosome without adopting a ligation competent conformation raises the intriguing possibility that this non-productive binding, or alternative modes of nucleosome binding mediated by the LigIIIα ZnF^112^, may tether LigIIIα at the nucleosomal nick until stochastic or DNA damage-dependent chromatin remodeling occurs. However, future work will be needed to further dissect the impact of stochastic and DNA damage-dependent chromatin remodeling on LigIIIα function during chromatin-based SSBR/BER.

## Methods

### Preparation of oligonucleotides

All oligonucleotides used to generate recombinant nucleosomes containing nicks were synthesized by Integrated DNA Technologies (Coralville, IA). A complete list of the oligonucleotides used for generating recombinant nucleosomes containing nicks can be found in Supplementary Table 6. The oligonucleotides were resuspended to a final concentration of 100 µM in a buffer containing 10 mM Tris (pH 7.5) and 1 mM EDTA. Equimolar amounts of complimentary oligonucleotides (see Supplementary Table 6) were mixed and diluted with a buffer containing 10 mM Tris (pH 7.5) and 1 mM EDTA to a final concentration of 10 µM. The complimentary oligonucleotides were then annealed by heating to 95°C for 2 minutes and cooling to 10 °C at a rate of −5 °C/min. The annealed oligonucleotides were stored short-term at 4 °C prior to nucleosome reconstitution.

### Purification of recombinant human histones

Recombinant *H. sapiens* histone H2A (UniProt identifier: P0C0S8), histone H2B (UniProt identifier: P62807), histone H3 C110A (UniProt identifier Q71DI3), and histone H4 (Uniprot identifier: P62805) proteins were generated from a tagless pet3a expression vector. The plasmids containing histone H2A, H3, and H4 were transformed and expressed in T7 Express lysY/I^q^ competent *E. coli* cells (New England BioLabs). The plasmid containing histone H2B was transformed and expressed in BL21-CodonPlus (DE3)-RIPL *E. coli* cells (Agilent). Th cells were grown in M9 minimal media at 37 °C to an OD600 of 0.4 and histone expression was induced with 0.4 mM (histone H2A, H2B, and H3) IPTG or 0.3 mM (histone H4) for 3 hours (histone H3 and H4) or 4 hours (histone H2A and H2B) at 37 °C. The cells were harvested via centrifugation and cell pellets stored long term at −80 °C. The purification of each individual histone was carried out using well-established methods^113-115^. In brief, the cell pellets were lysed via sonication in a buffer containing 50 mM Tris (pH 7.5), 100 mM NaCl, 1 mM benzamidine, 1 mM DTT, and 1 mM EDTA, and the cell lysate clarified via centrifugation. The resulting pellet was washed two times with a buffer containing 50 mM Tris (pH 7.5), 100 mM NaCl, 1 mM benzamidine, 1 mM DTT, 1 mM EDTA, and 1% Triton X-100, and further washed one time with a buffer containing 50 mM Tris (pH 7.5), 100 mM NaCl, 1 mM benzamidine, 1 mM DTT, and 1 mM EDTA. The histones were then extracted from inclusion bodies under denaturing condition using 6 M Guanidinium-HCl. The extracted histones were then purified under gravity flow using a combination of anion-exchange and cation-exchange chromatography. The purified histones were dialyzed against H_2_O, lyophilized, and stored long term at −20 °C.

### Generation of H2A/H2B dimer and H3/H4 tetramer

H2A/H2B dimers and H3/H4 tetramers were prepared prior to nucleosome reconstitution using established methods^113-115^. To generate H2A/H2B dimers, lyophilized histone H2A and H2B were resuspended in a buffer containing 20 mM Tris (pH 7.5), 6 M Guanidinium-HCl, and 10 mM DTT. The resuspended histone H2A and H2B were then mixed in a 1:1 molar ratio and dialyzed three times against a buffer containing 20 mM Tris (pH 7.5), 2 M NaCl, and 1 mM EDTA at 4 °C. To generate H3/H4 tetramers, lyophilized histone H3 and H4 were resuspended in a buffer containing 20 mM Tris (pH 7.5), 6 M Guanidinium-HCl, and 10 mM DTT. The resuspended histone H3 and H4 proteins were then mixed in a 1:1 molar ratio and dialyzed three times against a buffer containing 20 mM Tris (pH 7.5), 2 M NaCl, and 1 mM EDTA at 4 °C. The refolded H2A/H2B dimer and H3/H4 tetramer were then purified using a HiPrep Sephacryl S-200 16/60 HR gel filtration column (Cytiva) in a buffer containing 20 mM Tris (pH 7.5), 2 M NaCl, and 1 mM EDTA. Fractions containing pure H2A/H2B dimer and H3/H4 tetramer were combined and stored long term at −20 °C in 50% glycerol.

### Nucleosome reconstitution

All nucleosomes were prepared using an established salt-dialysis method ^113-115^. H2A/H2B dimer, H3/H4 tetramer, and annealed Nick-DNA were mixed at a 2.2:1:1 molar ratio, respectively. Nucleosomes were reconstituted by sequentially decreasing the concentration of NaCl from 2 M NaCl to 1.5 M NaCl, to 1.0 M NaCl, to 0.75 M NaCl, to 0.5 M NaCl, to 0.25 M NaCl, and to 0 M NaCl over 24 h. The reconstituted nucleosomes were then purified by sucrose gradient ultracentrifugation (10–40% gradient). Final nucleosome purity and homogeneity was assessed via native PAGE (5%, 59:1 acrylamide:bis-acrylamide ratio) and the nucleosomes stored in a buffer containing 10 mM Tris (pH 7.5) and 1 mM EDTA at 4 °C. Native PAGE gels of the final nucleosomes can be found in Supplementary Fig. 1.

### Nucleosome stability assays

Nucleosome stability assays were performed by incubating each NCP (200 nM) in a reaction buffer containing 20 mM HEPES (pH 7.5), 50 mM NaCl, 1 mM MgCl_2_, 0.1 mM TCEP, and 0.2 mg/ml BSA. Samples were taken immediately after resuspension (0′ time point) and after incubation for 24 hours at 37 °C. Nucleosomes were mixed with 2x sucrose loading dye (10% sucrose, 0.025% bromophenol blue, 0.1 mg/ml BSA, 10 mM Tris-HCl (pH 7.5), and 1 mM EDTA) and nucleosome integrity assessed by native polyacrylamide gel electrophoresis (5%, 59:1 acrylamide:bis-acrylamide). The gels were stained with SYBR Gold (Thermo Fisher Scientific) and the bands corresponding to nucleosome and free DNA quantified using ImageJ^116^.

### Purification of recombinant human LigIIIα

The codon-optimized DNA for *H. sapiens* LigIIIα was cloned into a pET24a bacterial expression vector containing a C-terminal His-tag by GenScript. LigIIIα was expressed and purified using a previously established method^38^ with minor modifications. The pET24a-LigIIIα plasmid was transformed and expressed in BL21 (DE3) *E. coli* cells (Agilent). The transformed cells were grown at 37 °C to an OD600 of 1.0 and LigIIIα expression induced with 0.5 mM IPTG for 16h – 18h at 25 °C. The resulting cells were lysed via sonication in a buffer containing 25 mM HEPES (pH 8.0), 300 mM NaCl, 0.25 mM TCEP, 10 mM imidazole, and a cocktail of protease inhibitor (AEBSF, leupeptin, benzamidine, pepstatin A). The lysate was clarified via centrifugation and loaded onto a HisTrap HP column (Cytiva) equilibrated with 25 mM HEPES (pH 8.0), 300 mM NaCl, 0.25 mM TCEP, 10 mM imidazole, and eluted in a buffer containing 25 mM HEPES (pH 8.0), 300 mM NaCl, 0.25 mM TCEP, 400 mM imidazole (0 – 100% gradient). The LigIIIα protein was then loaded onto a HiTrap Heparin HP column (Cytiva) equilibrated with 25 mM HEPES (pH 7.5), 200 mM NaCl, and 0.1 mM TCEP, and eluted in a buffer containing 25 mM HEPES (pH 7.5), 1000 mM NaCl, and 0.1 mM TCEP (0 – 100% gradient). The LigIIIα protein was further purified using a HiPrep SP HP column (Cytiva) equilibrated with 25 mM HEPES (pH 7.5), 200 mM NaCl, and 0.1 mM TCEP, and eluted in a buffer containing 25 mM HEPES (pH 7.5), 1000 mM NaCl, and 0.1 mM TCEP (0 – 100% gradient). LigIIIα was then adenylylated by incubation with 20 mM MgCl_2_ and 1 mM ATP for 60 mins at 4 °C. The LigIIIα protein was then polished via size exclusion chromatography using a HiPrep Sephacryl S-200 26/60 HR (Cytiva) equilibrated in a buffer containing 20 mM HEPES (pH 7.5), 150 mM NaCl, and 0.1 mM TCEP. The purity of LigIIIα was confirmed by denaturing SDS PAGE and the purified LigIIIα was stored long term at −80 °C. An SDS PAGE gel of the purified LigIIIα protein can be found in Supplementary Fig. 1c.

### Purification of recombinant human XRCC1

The DNA for *H. sapiens* XRCC1 was cloned into a pET-His-GFP bacterial expression vector using GenScript. The pET-His-GFP-XRCC1 plasmid was transformed and expressed in BL21 (DE3)-RIPL *E. coli* cells (Agilent). The transformed cells were grown at 37 °C to an OD600 of 1.0 and XRCC1 expression induced with 0.5 mM IPTG for 16h – 18h at 18 °C. The resulting cells were lysed via sonication in a buffer containing 25 mM HEPES (pH 8.0), 500 mM NaCl, 0.25 mM TCEP, 10 mM imidazole, and a cocktail of protease inhibitor (AEBSF, leupeptin, benzamidine, pepstatin A). The lysate was clarified via centrifugation and loaded onto a gravity flow column containing HisPur Ni-NTA Resin (Thermo Fisher Scientific) equilibrated with 25 mM HEPES (pH 8.0), 500 mM NaCl, 0.25 mM TCEP, 10 mM imidazole, and eluted in a buffer containing 25 mM HEPES (pH 8.0), 300 mM NaCl, 0.25 mM TCEP, 400 mM imidazole. The XRCC1 protein was then liberated from the His-GFP tag via incubation with PreScission Protease, loaded onto a HiTrap Heparin HP column (Cytiva) equilibrated with 25 mM HEPES (pH 7.5), 200 mM NaCl, and 0.1 mM TCEP, and eluted in a buffer containing 25 mM HEPES (pH 7.5), 1000 mM NaCl, and 0.1 mM TCEP (0 – 100% gradient). The XRCC1 protein was further purified using a HiPrep SP HP column (Cytiva) equilibrated with 25 mM HEPES (pH 7.5), 200 mM NaCl, and 0.1 mM TCEP, and eluted in a buffer containing 25 mM HEPES (pH 7.5), 1000 mM NaCl, and 0.1 mM TCEP (0 – 100% gradient). The XRCC1 protein was then polished via size exclusion chromatography using a HiPrep Sephacryl S-200 26/60 HR (Cytiva) equilibrated in a buffer containing 20 mM HEPES (pH 7.5), 100 mM NaCl, and 0.1 mM TCEP. The purity of XRCC1 was confirmed by denaturing SDS PAGE and the purified XRCC1 stored long term at −80 °C. An SDS PAGE gel of the purified XRCC1 protein can be found in Supplementary Fig. 1c.

### XRCC1-LigIIIα complex formation

The XRCC1-LigIIIα complex was generated by mixing purified recombinant XRCC1 and adenylylated LigIIIα at a 1:1 molar ratio in a buffer containing 25 mM HEPES (pH 7.5), 100 mM NaCl, and 0.1 mM TCEP. The reactions were incubated on ice for 2h, and the XRCC1-LigIIIα complex purified via size exclusion chromatography using a Superdex 200 Increase 10/300 GL (Cytiva) equilibrated in a buffer containing 25 mM HEPES (pH 7.5), 100 mM NaCl, and 0.1 mM TCEP. The purity of the XRCC1-LigIIIα complex was confirmed by denaturing SDS PAGE and the purified XRCC1-LigIIIα complex stored long term at −80 °C. An SDS PAGE gel of the purified XRCC1-LigIIIα complex can be found in Supplementary Fig. 1c.

### Single-turnover kinetic analysis of LigIIIα ligation

Single-turnover kinetic reactions were carried out by mixing LigIIIα (2 µM) or XRCC1-LigIIIα (2 µM) with each respective Nick-NCP (100 nM) or Nick-DNA (100 nM) in a buffer containing 50 mM HEPES (pH 7.5), 100 mM NaCl, 1 mM TCEP, and 0.1 mg/ml bovine serum albumin. The reactions were incubated for 10 mins at 37 °C, ligation initiated through the addition of Mg^2+^-ATP (2 mM Mg^2+^, 1 mM ATP), and the ligation reactions quenched at various time points with a solution containing 100 mM EDTA, 80% deionized formamide, 0.25 mg/ml bromophenol blue and 0.25 mg/ml xylene cyanol. For Nick-NCP0, Nick-NCP−2, Nick-NCP−4, the reactions time points were 10 sec, 20 sec, 30 sec, 60 sec, 120 sec, 240 sec, 600 sec, 1200 sec, 1800 sec, 3600 sec, 7200 sec, and 14400 sec. For Nick-DNA and Nick-NCP−6, the reactions time points were 5 sec, 10 sec, 15 sec, 20 sec, 30 sec, 60 sec, 120 sec, 240 sec, 600 sec, 1200 sec, 1800 sec, and 3600 sec. The quenched reactions were incubated at 95 °C for 5 mins and substrate and products bands separated by 8% (Nick-NCP0, Nick-NCP−2, and Nick-NCP−4) or 12% (Nick-DNA and Nick-NCP−6) denaturing urea polyacrylamide gel electrophoresis (29:1 acrylamide:bis-acrylamide). The bands corresponding to substrate and product were visualized by the 6-FAM label using a ChemiDoc MP imaging system (Bio-Rad) The substrate and product bands were quantified using ImageJ^116^ and the data fit to the double exponential equation:

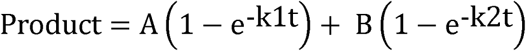

Where *A* and *B* are the amplitude of the fast and slow phase, respectively, and *k1* and *k2* are the ligation rates (*k_obs_*) corresponding to the fast and slow phase, respectively. Each time point represents the average of at least three independent replicate experiments, and the error bars represent the standard deviation of the three independent replicate experiments. The *k*_obs_ is reported as the mean of the three independent replicate experiments ± the standard deviation of the three independent replicate experiments.

### Electrophoretic mobility shift assays

Electrophoretic mobility shift assay reactions were carried out by mixing each respective NCP (20 nM) with increasing concentrations of LigIIIα (0 - 1000 µM) or XRCC1-LigIIIα (0 - 1000 µM) in a buffer containing 50 mM HEPES (pH 7.5), 100 mM NaCl, 1 mM TCEP, 0.1 mg/ml bovine serum albumin, 5% sucrose, and 0.25 % (w/v) bromophenol blue. The reactions were incubated for 10 mins at 4 °C, and the reactions species separated by native polyacrylamide gel electrophoresis (5%, 59:1 acrylamide:bis-acrylamide) for 50 min at 4 °C. The disappearance of the free NCP band was quantified using ImageJ^116^ and fit to a modified Hill equation:

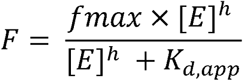

Where *fmax* is the maximal nucleosome bound, *[E]* is the total LigIIIα or XRCC1-LigIIIα concentration, *h* is the Hill coefficient, and *K*_d,app_ is the apparent dissociation constant. Each experimental point represents the average of three independent replicate experiments, and the error bars represent the standard deviation of the three independent replicate experiments. The *K*_d,app_ is reported as the mean of the three independent replicate experiments ± the standard deviation of the three independent replicate experiments.

### Cryo-EM sample and grid preparation

LigIIIα-Nick-NCP complexes were generated by mixing LigIIIα (2.0 μM) with each individual Nick-NCP (1.0 μM) in a buffer containing 25 mM HEPES (pH 7.5), 25 mM NaCl, 1 mM TCEP, and 1 mM EDTA. The XRCC1-LigIIIα-Nick-NCP−6 complex was generated by mixing XRCC1-LigIIIα (2.0 μM) with Nick-NCP−6 (1.0 μM) in a buffer containing 25 mM HEPES (pH 7.5), 25 mM NaCl, 1 mM TCEP, and 1 mM EDTA. The LigIIIα-ND-NCP complex was generated by mixing LigIIIα (2.0 μM) with NCP (1.0 μM) in a buffer containing 25 mM HEPES (pH 7.5), 25 mM NaCl, and 1 mM TCEP. All complexes were incubated for 20 mins at 4 °C, stabilized via glutaraldehyde crosslinking (0.2 %) for 20 min at 4 °C, and immediately purified via a Superdex S200 Increase 10/300 GL (Cytiva) equilibrated in a buffer containing 25 mM HEPES (pH 7.5), 50 mM NaCl, 1 mM TCEP, and 1 mM EDTA. Fractions containing complexes were combined and concentrated to 1.2 μM for LigIIIα-ND-NCP, 1.2 μM for LigIIIα-Nick-NCP0, 0.7 μM for LigIIIα-Nick-NCP−2, 1.2 μM for LigIIIα-Nick-NCP−4, 1.2 μM for LigIIIα-Nick-NCP−6, and 1.2 μM for XRCC1-LigIIIα-Nick-NCP−6. The final sample quality was assessed by native PAGE (5%, 59:1 acrylamide:bis-acrylamide ratio). Native PAGE gels of the final cryo-EM samples can be found in Supplementary Figs. 4-8 and 23. Cryo-EM grids were generated by applying each sample (3 μL) to a glow discharged Quantifoil R2/1 300 mesh copper cryo-EM grid. The cryo-EM grids were blotted for 3.5 – 4s at 4 °C and 95% humidity and plunge-frozen in liquid ethane using a Vitrobot Mark IV (Thermo Fisher Scientific).

### Cryo-EM data collection and processing

All cryo-EM data collections were performed at the University of Virginia Molecular Electron Microscopy Facility on a 300 kV TFS Titan Krios cryo-TEM equipped with a Gatan K3/GIF direct electron detector using EPU. All data collection parameters can be found in Supplementary Tables 1-4. The cryo-EM data was processed with cryoSPARC^117^ using similar data processing workflows. Each individual data processing workflow can be found in Supplementary Figs. 4-8 and 23. In brief, the micrographs for each dataset were initially imported into cryoSPARC and subjected to pre-processing using patch motion correction and patch CTF-estimation. An initial cycle of blob particle picking was performed to generate a training set for Topaz, and Topaz^118^ used to generate the final particle stack. Ab initio models were then generated using the final particle stacks, and heterogeneous refinement(s) and/or 3D classifications were performed to separate the free Nick-NCPs and LigIIIα-containing Nick-NCP complexes. Following classification, the free NCP and LigIIIα-containing Nick-NCP reconstructions were subjected to local CTF refinement and reference-based motion correction prior to a final non-uniform refinement. To improve interpretability of the LigIIIα-Nick-NCP−2, LigIIIα-Nick-NCP−4, LigIIIα-Nick-NCP−6, and XRCC1-LigIIIα-Nick-NCP−6 reconstructions, the maps were further subjected to local refinement without particle subtraction using a mask for LigIIIα and the adjacent nucleosomal DNA. A composite map was then generated by combining the consensus and local refinement maps using PHENIX combine focus maps. The local resolution estimation for each composite map was generated using cryoSPARC local resolution estimation after combining the half maps from the consensus and local refinements in PHENIX.

### Model building and refinement

An initial nucleosome model was generated from a previously determined cryo-EM structure of a nucleosome containing a 1-nt gap (PDB: 9DWF)^74^, and an initial LigIIIα model was generated from a previously determined X-ray crystal structure of LigIIIα bound to a nicked DNA (PDB: 3L2P)^29^. For the ND-NCP and Nick-NCP structures, the initial nucleosome model was rigid-body docked into each respective map using ChimeraX^119^, and the models further refined using PHENIX^120^ and Coot^121^ with protein and DNA secondary structure restraints. For the LigIIIα-Nick-NCP0 structures, the initial nucleosome and LigIIIα models were rigid-body docked into the map using ChimeraX^119^, and the models further refined using PHENIX^120^ and Coot^121^ with protein and DNA secondary structure restraints. For the LigIIIα-Nick-NCP−6, LigIIIα-Nick-NCP−4, LigIIIα-Nick-NCP−2, and XRCC1-LigIIIα-Nick-NCP−6 structures, the initial nucleosome and LigIIIα model (from the LigIIIα-Nick-NCP0 structure) were rigid-body docked into the map using ChimeraX^119^, and the models further refined using PHENIX^120^ and Coot^121^ with protein and DNA secondary structure restraints. Each model was validated using MolProbity^122^ and PHENIX comprehensive cryo-EM validation^120^ prior to deposition into the Protein Data Bank (PDB).

The model coordinates for all NCP, LigIIIα-Nick-NCP, and XRCC1-LigIIIα-Nick-NCP structures were deposited into the PDB under accession number 10XZ for ND-NCP, 10YA for Nick-NCP0, 10YB for Nick-NCP−2, 10YC for Nick-NCP−4, 10YD for Nick-NCP−6, 10YE for LigIIIα-Nick-NCP0, 10YF for LigIIIα-Nick-NCP−2, 10YG for LigIIIα-Nick-NCP−4, 10YH for LigIIIα-Nick-NCP−6, and 10YI for XRCC1-LigIIIα-Nick-NCP−6. The consensus cryo-EM maps for all NCP, LigIIIα-Nick-NCP, and XRCC1-LigIIIα-Nick-NCP structures were deposited into the Electron Microscopy Data Bank (EMDB) under accession number EMD-75521 for ND-NCP, EMD-75522 for Nick-NCP0, EMD-75523 for Nick-NCP−2, EMD-75524 for Nick-NCP−4, and EMD-75525 for Nick-NCP−6, EMD-75526 for LigIIIα-Nick-NCP0, EMD-75531 for LigIIIα-Nick-NCP−2, EMD-75533 for LigIIIα-Nick-NCP−4, EMD-75535 for LigIIIα-Nick-NCP−6, and EMD-75537 for XRCC1-LigIIIα-Nick-NCP−6. The LigIIIα local refine cryo-EM maps for LigIIIα-Nick-NCP and XRCC1-LigIIIα-Nick-NCP structures were deposited into the EMDB under accession number EMD-75532 for LigIIIα-Nick-NCP−2, EMD-75534 for LigIIIα-Nick-NCP−4, EMD-75536 for LigIIIα-Nick-NCP−6, and EMD-75538 for XRCC1-LigIIIα-Nick-NCP−6. The composite cryo-EM maps for LigIIIα-Nick-NCP and XRCC1-LigIIIα-Nick-NCP structures were deposited into the EMDB under accession number EMD-75527 for LigIIIα-Nick-NCP−2, EMD-75528 for LigIIIα-Nick-NCP−4, EMD-75529 for LigIIIα-Nick-NCP−6, and EMD-75530 for XRCC1-LigIIIα-Nick-NCP−6.

### Molecular Dynamics Simulations

All simulations were performed with the AMBER20 suite of programs^123^. The starting models of the ND-NCP, Nick-NCP0, Nick-NCP−2, Nick-NCP−4, and Nick-NCP−6 used in the MD simulations were computationally generated using a previously determined cryo-EM structure of a nucleosome containing a 1-nt gap (PDB: 9DWF)^74^. The ff19SB^124^ and OL21^125^ force fields were used for protein and DNA. Each system was solvated in a truncated octahedron box of OPC water^126^ with a buffer of 12 Å around the solute. Sodium and chloride ions were added to reach a final salt concentration of 0.15 M, resulting in final systems of approximatively 250,000 atoms. Each system was subjected to a 10,000-step minimization, followed by a 10 ps NVT thermalization with 10 kcal/mol restraints on the solute atoms. Two 4 ns NPT equilibration steps with decreasing restraints on the solute backbone atoms were then carried out, followed by a final 40 ns equilibration, and a 1 μs unrestrained production run. Three replicates were performed with random starting velocities for the ND-NCP, Nick-NCP0, Nick-NCP−2, Nick-NCP−4, and Nick-NCP−6 systems. The Particle Mesh Ewald approach was used with an 8 Å cutoff for electrostatics contributions, and a 4 fs timestep enabled by the use of Hydrogen Mass Repartitioning^127^. The Langevin thermostat and Berendsen barostat were used to maintain the temperature and pressure at 300 K and 1 bar, respectively. The per nucleotide contribution to the DNA flexibility was quantified using a PCA-based analysis^128^ previously used on similar systems^129,130^. Contact maps were generated with the cpptraj tool of AMBER20 using default parameters.

## Acknowledgements

This research was supported by startup funds from the University of Virginia School of Medicine and the University of Virginia Comprehensive Cancer Center (T.M.W), the National Institute of Environmental Health Sciences R35ES031638 (B.V.H) and R01ES012512 (A.E.T), the Agence Nationale de la Recherche ANR-24-CE29-2473 (E.B.), and the National Institute of General Medical Sciences R35GM155098 (A.M.W). This work used the 200 kV TFS Glacios cryo-TEM and the 300 kV TFS Titan Krios cryo-TEM within the University of Virginia Molecular Electron Microscopy Core, which is supported by the University of Virginia School of Medicine (RRID:SCR_019031) and the University of Virginia Comprehensive Cancer Center NCI Cancer Center Support Grant P30-CA044579. The Titan Krios (S10-RR025067) and K3/GIF (U24-GM116790) within the Molecular Electron Microscopy Core were purchased in part or in full using designated NIH grants. We also acknowledge Michael Purdy, Ph.D. and David Cooper, Ph.D. from the Molecular Electron Microscopy Core at the University of Virginia School of Medicine for their generous help with cryo-EM screening and data collection. This work also used High Performance Computing resources provided by the EXPLOR center of University of Lorraine (2019CPMXX0983) and by GENCI (A0190716838).

## Author Contributions

D.J.B., N.I.M., and T.M.W conceptualized the experiments and established the general research goals. D.J.B. and N.I.M. generated recombinant nucleosomes and recombinant proteins for biochemical assays and cryo-EM. N.I.M., A.G.M, C.A.K., A.M.W., and T.M.W. performed and analyzed the biochemical experiments. N.G. and E.B performed and analyzed the MD simulations. D.J.B performed cryo-EM sample preparation and validation. D.J.B. and T.M.W. processed and analyzed the cryo-EM datasets. D.J.B. and T.M.W. performed model building and refinement. D.J.B., N.I.M., and T.M.W wrote the manuscript, with input from C.A.K., A.E.T., B.V.H., N.G., E.B., and A.M.W.

## Data Availability Statement

Atomic coordinates for the reported structures have been deposited with the Protein Data Bank under accession numbers <u>10XZ</u>, <u>10YA</u>, <u>10YB</u>, <u>10YC</u>, <u>10YD</u>, <u>10YE</u>, <u>10YF</u>, <u>10YG</u>, <u>10YH</u>, and <u>10YI</u>. All cryo-EM maps are available from the Electron Microscopy Data Bank under accession numbers <u>EMD-75521</u>, <u>EMD-75522</u>, <u>EMD-75523</u>, <u>EMD-75524</u>, <u>EMD-75525</u>, <u>EMD-75526</u>, <u>EMD-75527</u>, <u>EMD-75528</u>, <u>EMD-75529</u>, <u>EMD-75530</u>, <u>EMD-75531</u>, <u>EMD-75532</u>, <u>EMD-75533</u>, <u>EMD-75534</u>, <u>EMD-75535</u>, <u>EMD-75536</u>, <u>EMD-75537</u>, and <u>EMD-75538</u>. Atomic coordinates for the initial nucleosome model and the LigIIIα-Nick-DNA model were obtained from the Protein Data bank under accession numbers 5DWF and 3L2P, respectively. The data from the single-turnover enzyme kinetics and EMSA experiments generated in this study are available in the Supplementary Information file. The raw data from the MD simulations were deposited into the Zenodo repository and are available from DOI:<u>10.5281/zenodo.19235695</u>. Source data are provided with this paper.

