## Supplementary Information File for "Molecular basis of nick ligation in the nucleosome by DNA Ligase IIIα"

**Supplementary Fig. 1: Validation of recombinant NCPs and recombinant proteins**

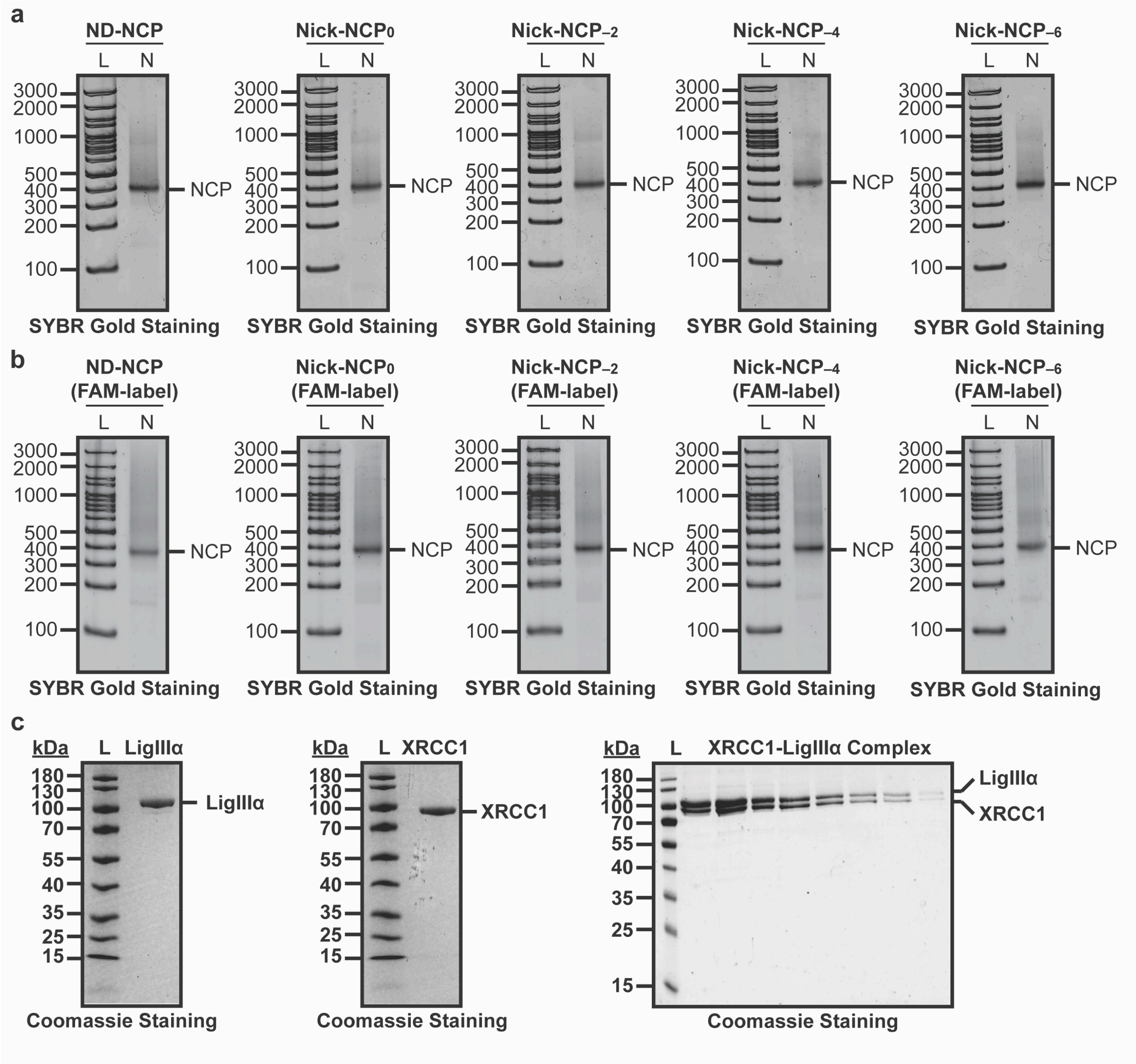

**Supplementary Fig. 1: Validation of recombinant NCPs and recombinant proteins**

**a,** Native PAGE gels confirming nucleosome formation and purity for the non-damaged NCP (ND-NCP), Nick-NCP$_{-6}$, Nick-NCP$_{-4}$, Nick-NCP$_{-2}$, and Nick-NCP$_{0}$ samples. The NCPs were detected using SYBR Gold staining. The lanes corresponding to 100 bp DNA ladder (L) and nucleosome sample (N) are labeled. **b,** Native PAGE gels confirming nucleosome formation and purity for the 6-carboxyfluorescein (FAM)-labeled ND-NCP, Nick-NCP$_{-6}$, Nick-NCP$_{-4}$, Nick-NCP$_{-2}$, and Nick-NCP$_{0}$ samples. The NCPs were detected using SYBR Gold staining. The lanes corresponding to a 100 bp DNA ladder (L) and nucleosome sample (N) are labeled. **c,** SDS-PAGE gels confirming the purity of LigIIIα (left), XRCC1 (middle), and the XRCC1-LigIIIα complex (right). The proteins were detected with Coomassie blue staining. The lanes corresponding to the protein ladder (L) are labeled. All source data in this figure are provided as a Source data file.

**Supplementary Fig. 2: Single-turnover kinetic analysis of LigIIIα nick ligation in the nucleosome**

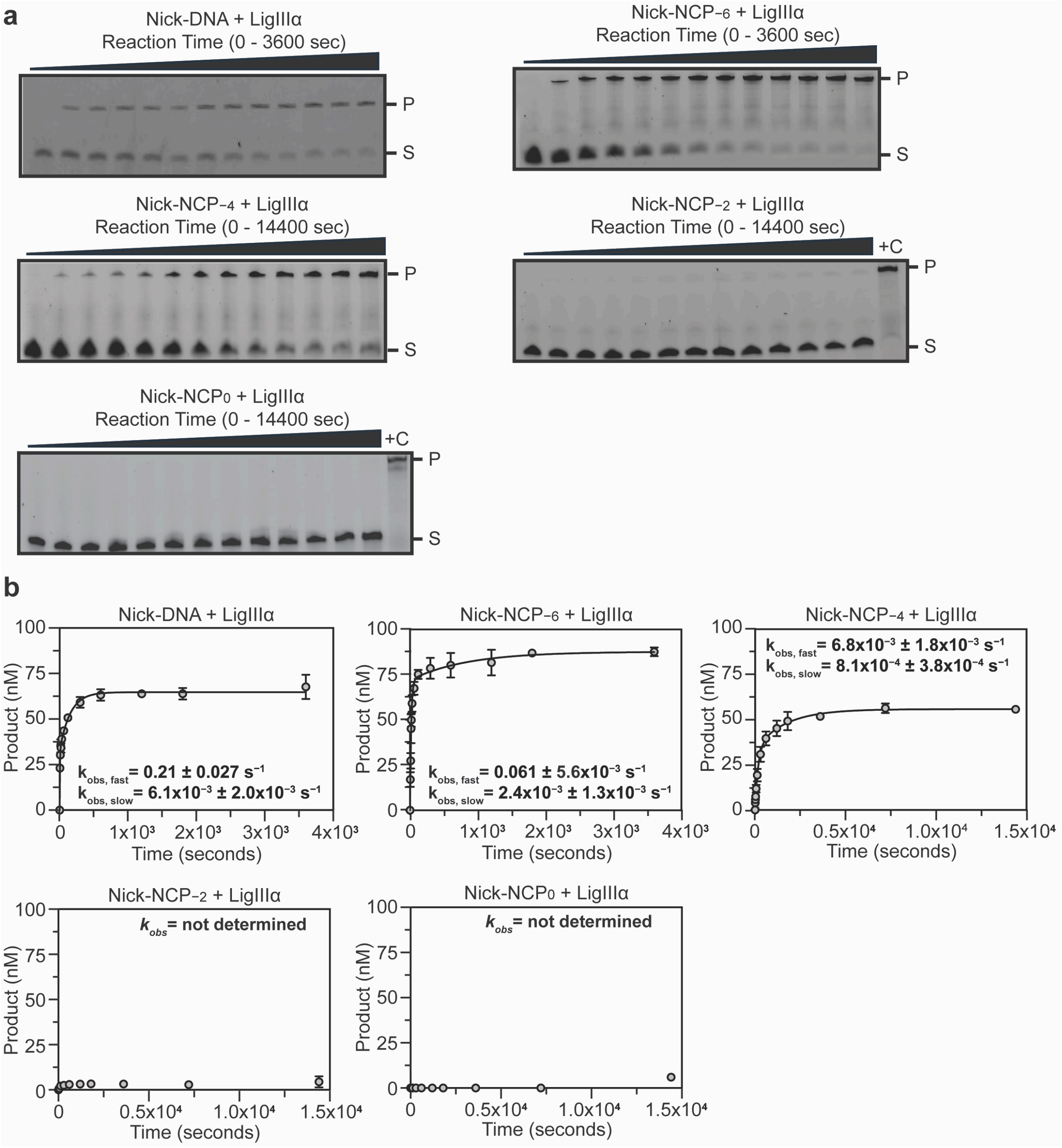

**Supplementary Fig. 2: Single-turnover kinetic analysis of LigIIIα nick ligation in the nucleosome**

**a,** Representative denaturing Urea-PAGE gels from the single-turnover kinetic experiments (STK) for LigIIIα with Nick-DNA, Nick-NCP$_{-6}$, Nick-NCP$_{-4}$, Nick-NCP$_{-2}$, and Nick-NCP$_{0}$. The substrate (S, Nick-DNA or Nick-NCP) and product (P, ligated DNA or NCP) were detected using the 6-FAM label on the nucleosomal DNA of each NCP. A 147 nt single-stranded DNA was loaded as a positive control (+C) for experiments with less than 10% product formed over the kinetic time course. The gels are representative of three independent STK experiments performed for LigIIIα with the Nick-DNA and each Nick-NCP. **b,** Quantification of the STK experiments for LigIIIα with Nick-DNA, Nick-NCP$_{-6}$, Nick-NCP$_{-4}$, Nick-NCP$_{-2}$, and Nick-NCP$_{0}$. The data points represent the mean ± standard deviation from three independent replicate experiments. The error bars are included for all experimental data points, but some error bars are smaller than the circles used to represent data points. The ligation rate ($k_{obs}$) is shown as an inset for each experiment and represents the mean ± standard error of the mean from the three independent replicate experiments. The ligation rate ($k_{obs}$) was not determined for experiments with less than 10% product formed over the kinetic time course. All source data in this figure are provided as a Source Data file.

**Supplementary Fig. 3: EMSA Analysis of the LigIIIα-Nick-NCP interaction**

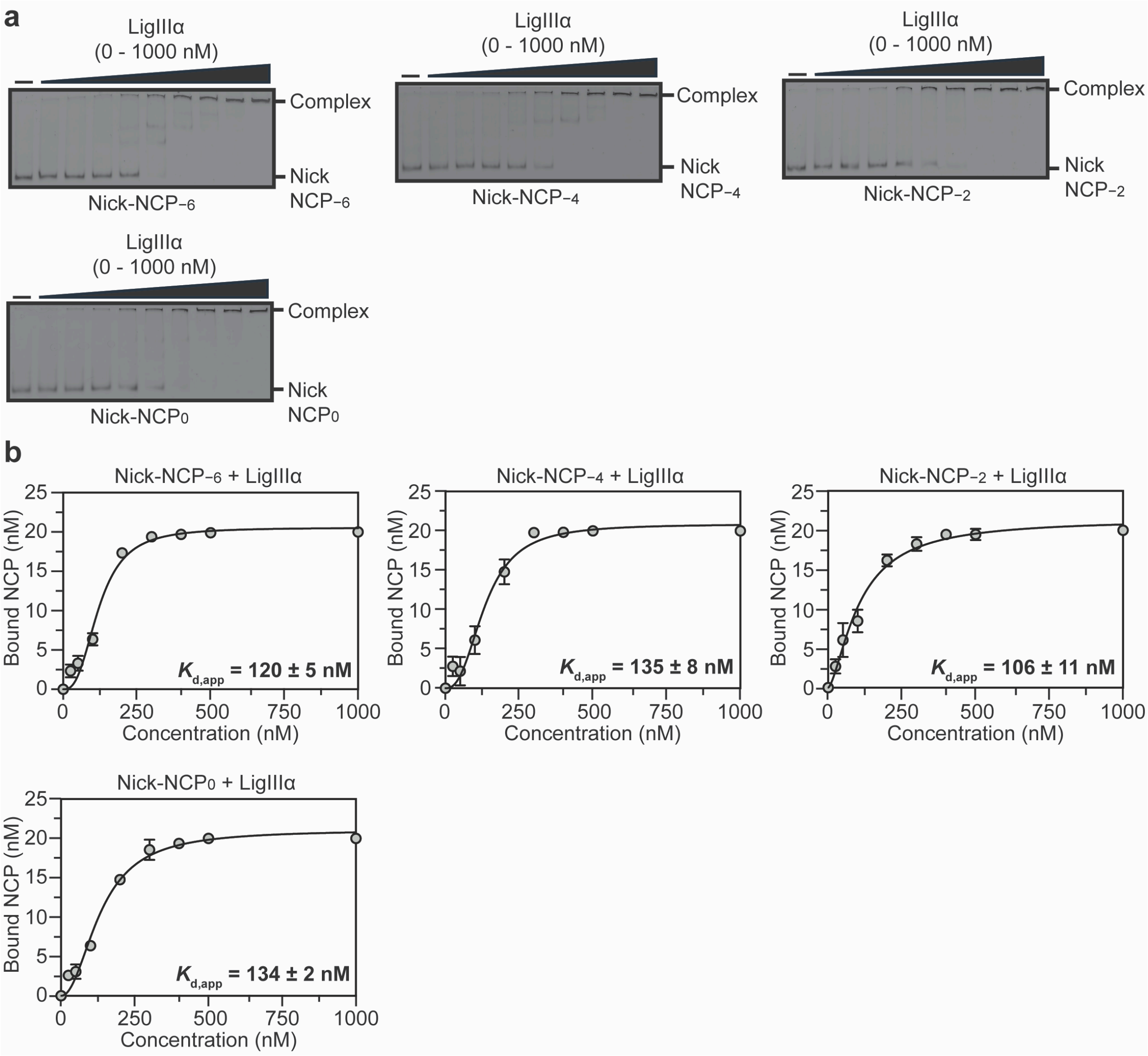

**Supplementary Fig. 3: EMSA Analysis of the LigIIIα-Nick-NCP interaction**

**a,** Representative native PAGE gels from the electrophoretic mobility shift assays (EMSAs) of LigIIIα with Nick-NCP$_{-6}$, Nick-NCP$_{-4}$, Nick-NCP$_{-2}$, and Nick-NCP$_{0}$. The free Nick-NCP and LigIIIα-Nick-NCP complex were detected using the 6-FAM label on the nucleosomal DNA of each NCP. The gels are representative of three independent EMSA experiments performed for LigIIIα and each Nick-NCP. **b,** Quantification of the EMSA experiments for LigIIIα with Nick-NCP$_{-6}$, Nick-NCP$_{-4}$, Nick-NCP$_{-2}$, and Nick-NCP$_{0}$. The data points represent the mean ± standard deviation from the three independent replicate experiments. The error bars are included for all experimental data points, but some error bars are smaller than the circles used to represent data points. The apparent binding affinity ($K_{d,app}$) is shown as an inset for each experiment and represents the mean ± standard deviation from the three independent replicate experiments. All source data in this figure are provided as a Source Data file.

**Supplementary Fig. 4: ND-NCP single particle analysis processing workflow**

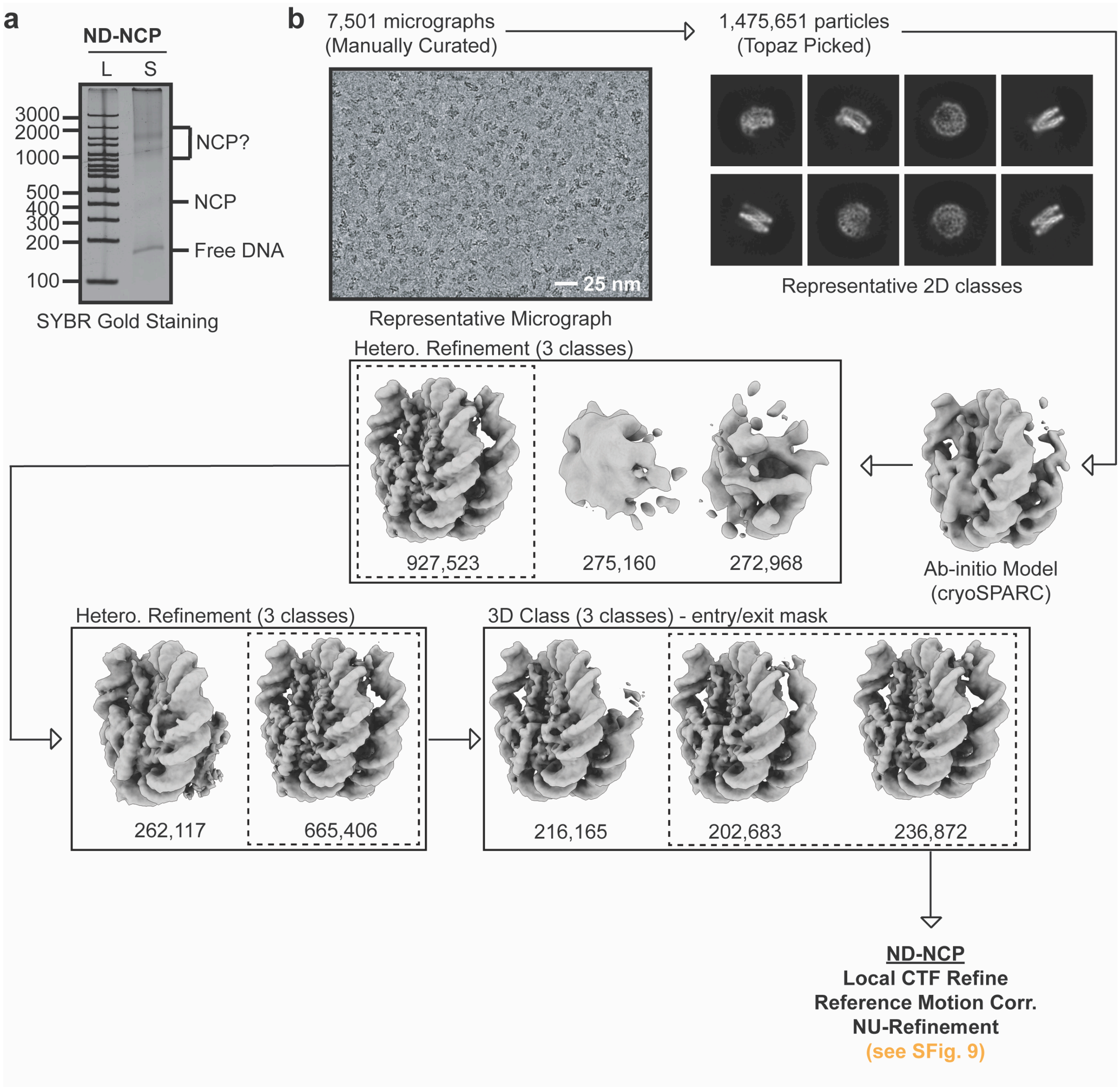

**Supplementary Fig. 4: ND-NCP single particle analysis processing workflow**

**a,** Native PAGE gel of the ND-NCP cryo-EM sample (S) and a 100 bp DNA ladder (L). The ND-NCP complexes were visualized with SYBR gold staining. **b,** Cryo-EM data processing workflow for the ND-NCP dataset. A representative micrograph (n=7,501) and representative 2D classes from the ND-NCP dataset are shown. The maps chosen for further classification and refinement throughout the cryo-EM data processing workflow are boxed. The final map, final model, and quality assessment metrics for ND-NCP can be found in Supplementary Fig. 9. All source data in this figure are provided as a Source data file.

**Supplementary Fig. 5: LigIIIα-Nick-NCP−6 single particle analysis processing workflow**

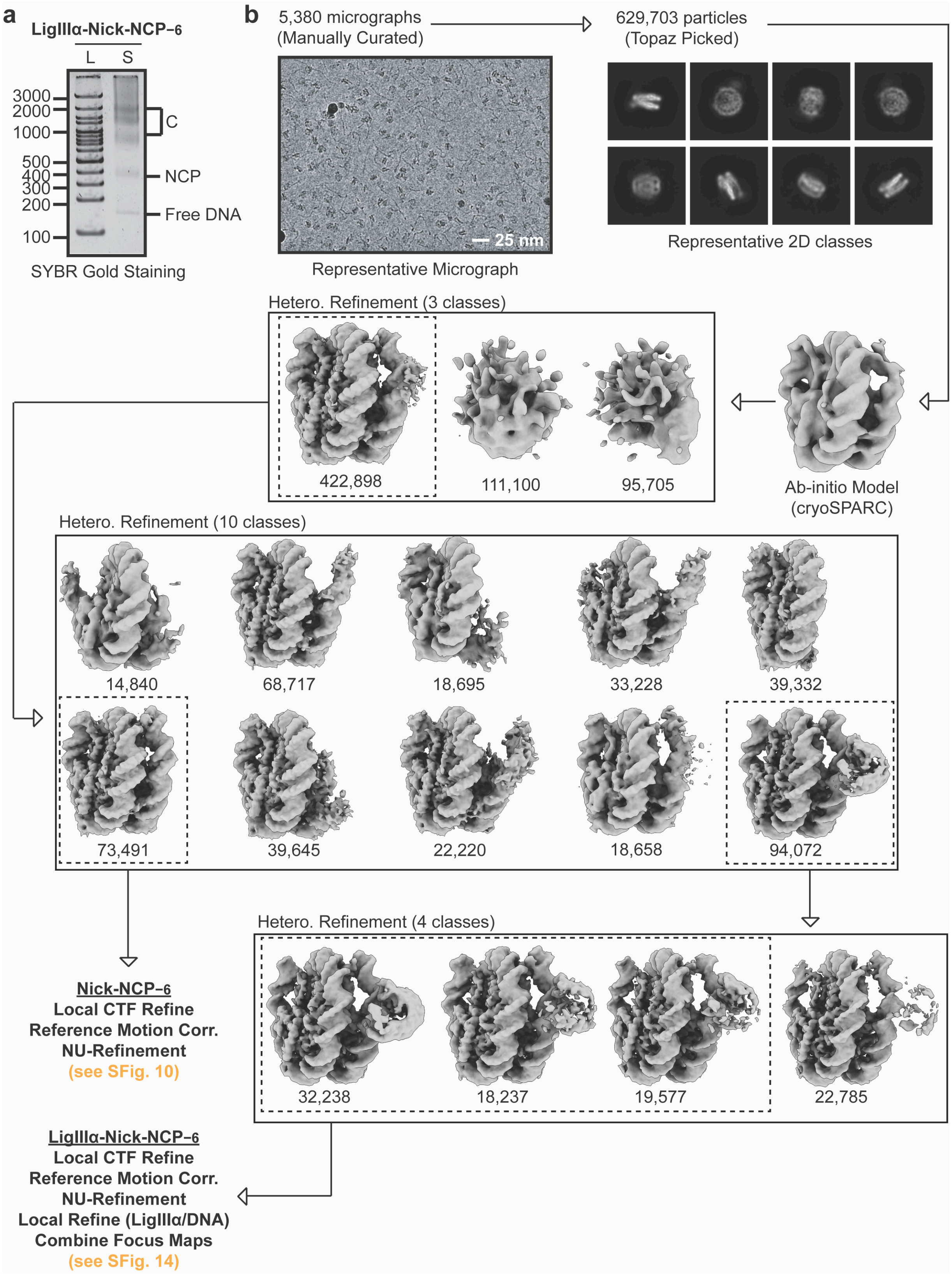

**Supplementary Fig. 5: LigIIIα-Nick-NCP−6 single particle analysis processing workflow**

**a,** Native PAGE gel of the LigIIIα-Nick-NCP−6 cryo-EM sample (S) and a 100 bp DNA ladder (L). The NCP−6 and LigIIIα-Nick-NCP−6 complex were visualized with SYBR gold staining. The bands corresponding to free DNA, the NCP, and LigIIIα-Nick-NCP−6 (C) are labeled. **b,** Cryo-EM data processing workflow for the LigIIIα-Nick-NCP−6 dataset. A representative micrograph (n=5,380) and representative 2D classes from the LigIIIα-Nick-NCP−6 dataset are shown. The maps chosen for further classification and/or refinement throughout the data processing pipeline are boxed. The final maps, final models, and quality assessment metrics for Nick-NCP−6 and LigIIIα-Nick-NCP−6 can be found in Supplementary Fig. 11 and Supplementary Fig. 17, respectively. All source data in this figure are provided as a Source data file.

**Supplementary Fig. 6: LigIIIα-Nick-NCP−4 single particle analysis processing workflow**

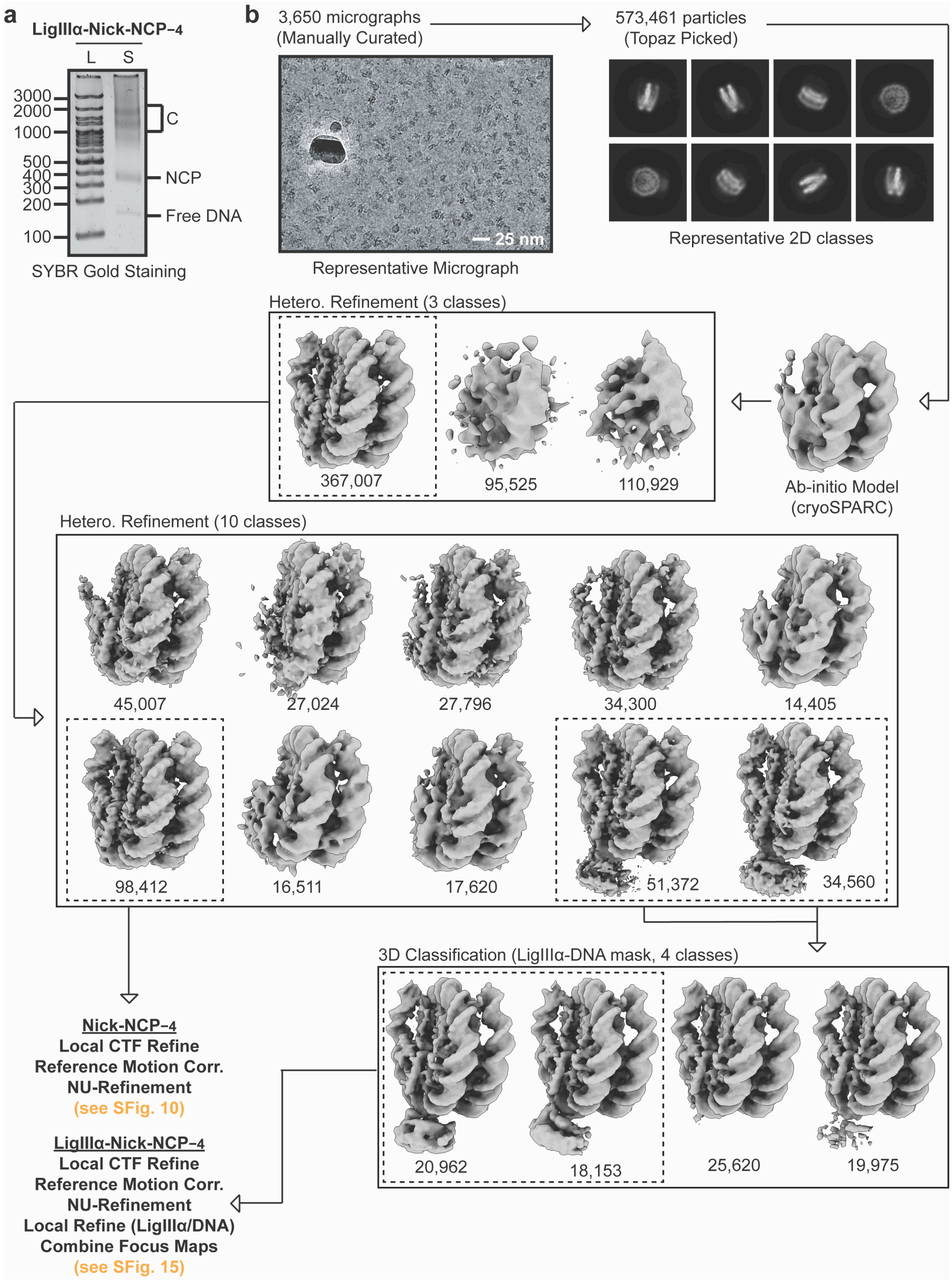

**Supplementary Fig. 6: LigIIIα-Nick-NCP−4 single particle analysis processing workflow**

**a,** Native PAGE gel of the LigIIIα-Nick-NCP−4 cryo-EM sample (S) and a 100 bp DNA ladder (L). The NCP−4 and LigIIIα-Nick-NCP−4 complex were visualized with SYBR gold staining. The bands corresponding to free DNA, the NCP, and LigIIIα-Nick-NCP−4 (C) are labeled. **b,** Cryo-EM data processing workflow for the LigIIIα-Nick-NCP−4 dataset. A representative micrograph (n=3,650) and representative 2D classes from the LigIIIα-Nick-NCP−4 dataset are shown. The maps chosen for further classification and/or refinement throughout the data processing pipeline are boxed. The final maps, final models, and quality assessment metrics for Nick-NCP−4 and LigIIIα-Nick-NCP−4 can be found in Supplementary Fig. 11 and Supplementary Fig. 16, respectively. All source data in this figure are provided as a Source data file.

**Supplementary Fig. 7: LigIIIα-Nick-NCP−2 single particle analysis processing workflow**

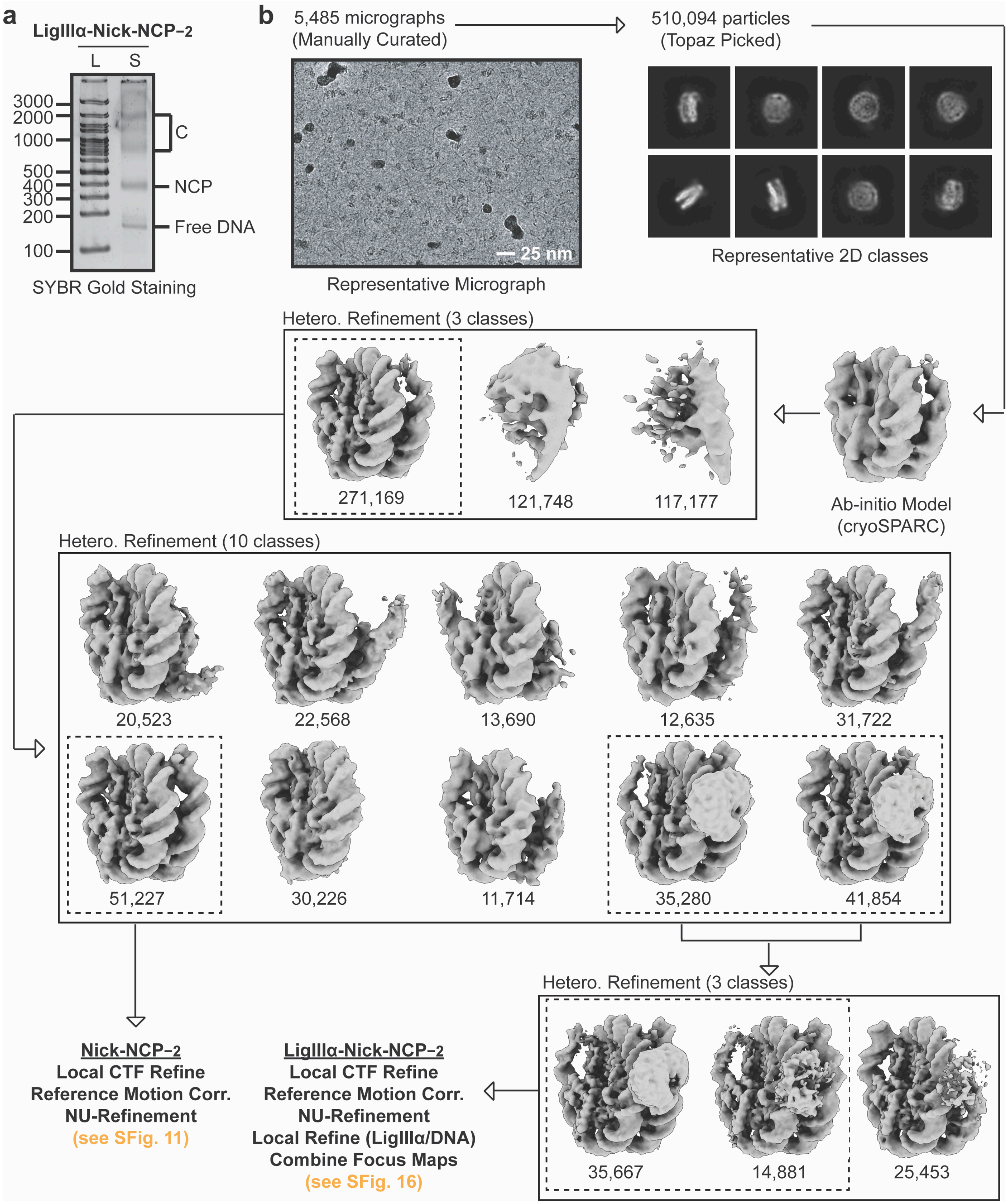

**Supplementary Fig. 7: LigIIIα-Nick-NCP$_{-2}$ single particle analysis processing workflow**

**a,** Native PAGE gel of the LigIIIα-Nick-NCP$_{-2}$ cryo-EM sample (S) and a 100 bp DNA ladder (L). The NCP$_{-2}$ and LigIIIα-Nick-NCP$_{-2}$ complex were visualized with SYBR gold staining. The bands corresponding to free DNA, the NCP, and LigIIIα-Nick-NCP$_{-2}$ (C) are labeled. **b,** Cryo-EM data processing workflow for the LigIIIα-Nick-NCP$_{-2}$ dataset. A representative micrograph (n=5,485) and representative 2D classes from the LigIIIα-Nick-NCP$_{-2}$ dataset are shown. The maps chosen for further classification and/or refinement throughout the data processing pipeline are boxed. The final maps, final models, and quality assessment metrics for Nick-NCP$_{-2}$ and LigIIIα-Nick-NCP$_{-2}$ can be found in Supplementary Fig. 10 and Supplementary Fig. 15, respectively. All source data in this figure are provided as a Source data file.

**Supplementary Fig. 8: LigIIIα-Nick-NCP0 single particle analysis processing workflow**

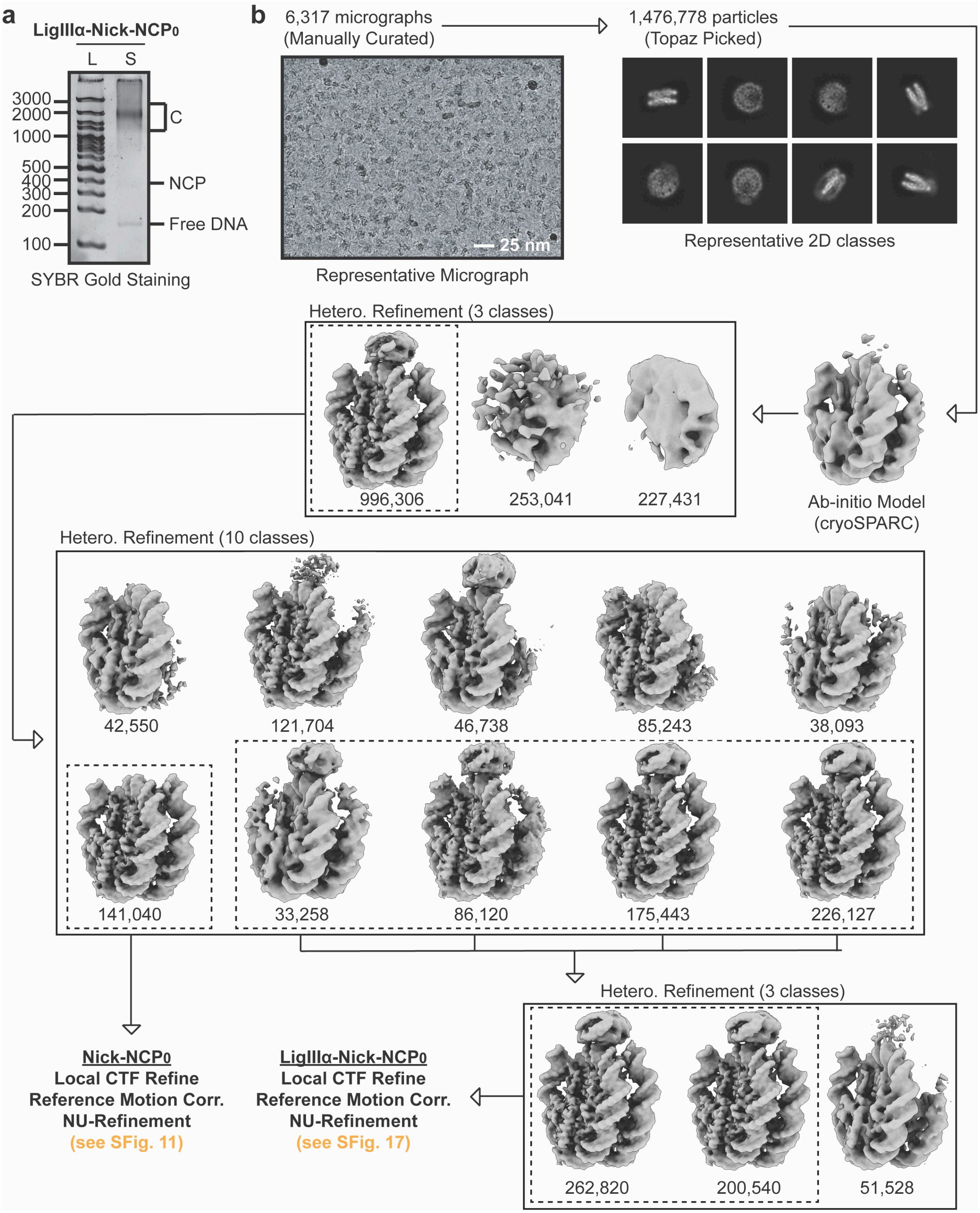

**Supplementary Fig. 8: LigIIIα-Nick-NCP0 single particle analysis processing workflow**

**a,** Native PAGE gel of the LigIIIα-Nick-NCP0 cryo-EM sample (S) and a 100 bp DNA ladder (L). The NCP0 and LigIIIα-Nick-NCP0 complex were visualized with SYBR gold staining. The bands corresponding to free DNA, the NCP, and LigIIIα-Nick-NCP0 (C) are labeled. **b,** Cryo-EM data processing workflow for the LigIIIα-Nick-NCP0 dataset. A representative micrograph (n=6,317) and representative 2D classes from the LigIIIα-Nick-NCP0 dataset are shown. The maps chosen for further classification and/or refinement throughout the data processing pipeline are boxed. The final maps, final models, and quality assessment metrics for Nick-NCP0 and LigIIIα-Nick-NCP0 can be found in Supplementary Fig. 10 and Supplementary Fig. 14, respectively. All source data in this figure are provided as a Source data file.

**Supplementary Fig. 9: ND-NCP map and model quality assessment**

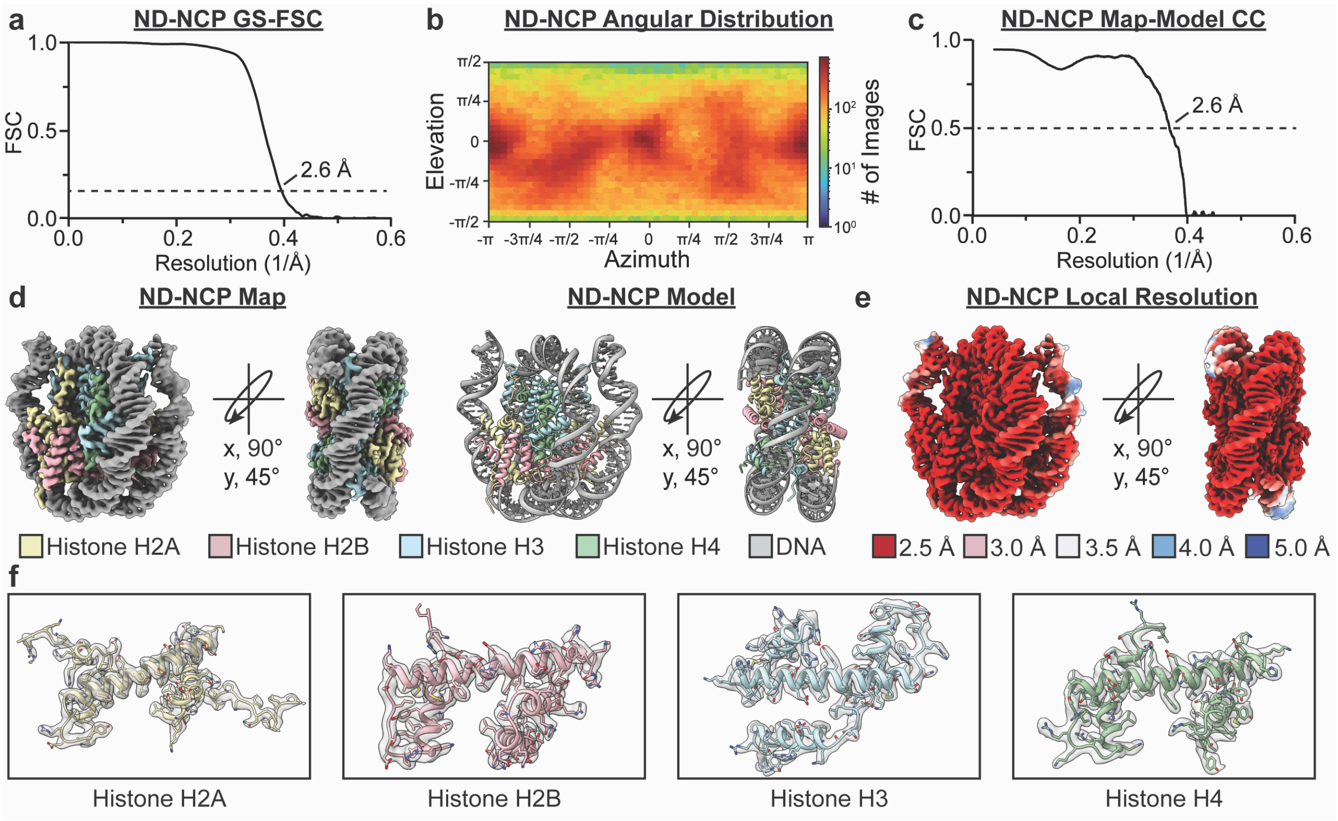

**Supplementary Fig. 9: ND-NCP map and model quality assessment**

**a,** Gold-standard Fourier shell correlation (GS-FSC) curves for the ND-NCP cryo-EM map (solid black line). The dashed line corresponds to GS-FSC - 0.143. **b,** Angular distribution heatmap for the ND-NCP cryo-EM map**. c,** Map-to-model FSC curves for the ND-NCP model and cryo-EM map. The dashed line corresponds to FSC - 0.5. **d,** The final 2.6 Å ND-NCP cryo-EM map and model shown in two different orientations. **e**, Local resolution estimation for the ND-NCP cryo-EM map shown in two different orientations. **f,** Representative segmented densities for histones H2A, H2B, H3, and H4 from the ND-NCP cryo-EM map. The representative segmented densities from the cryo-EM map are shown as transparent gray surfaces. All source data in this figure are provided as a Source data file.

**Supplementary Fig. 10: Nick-NCP–6 and Nick-NCP–4 map and model quality assessment**

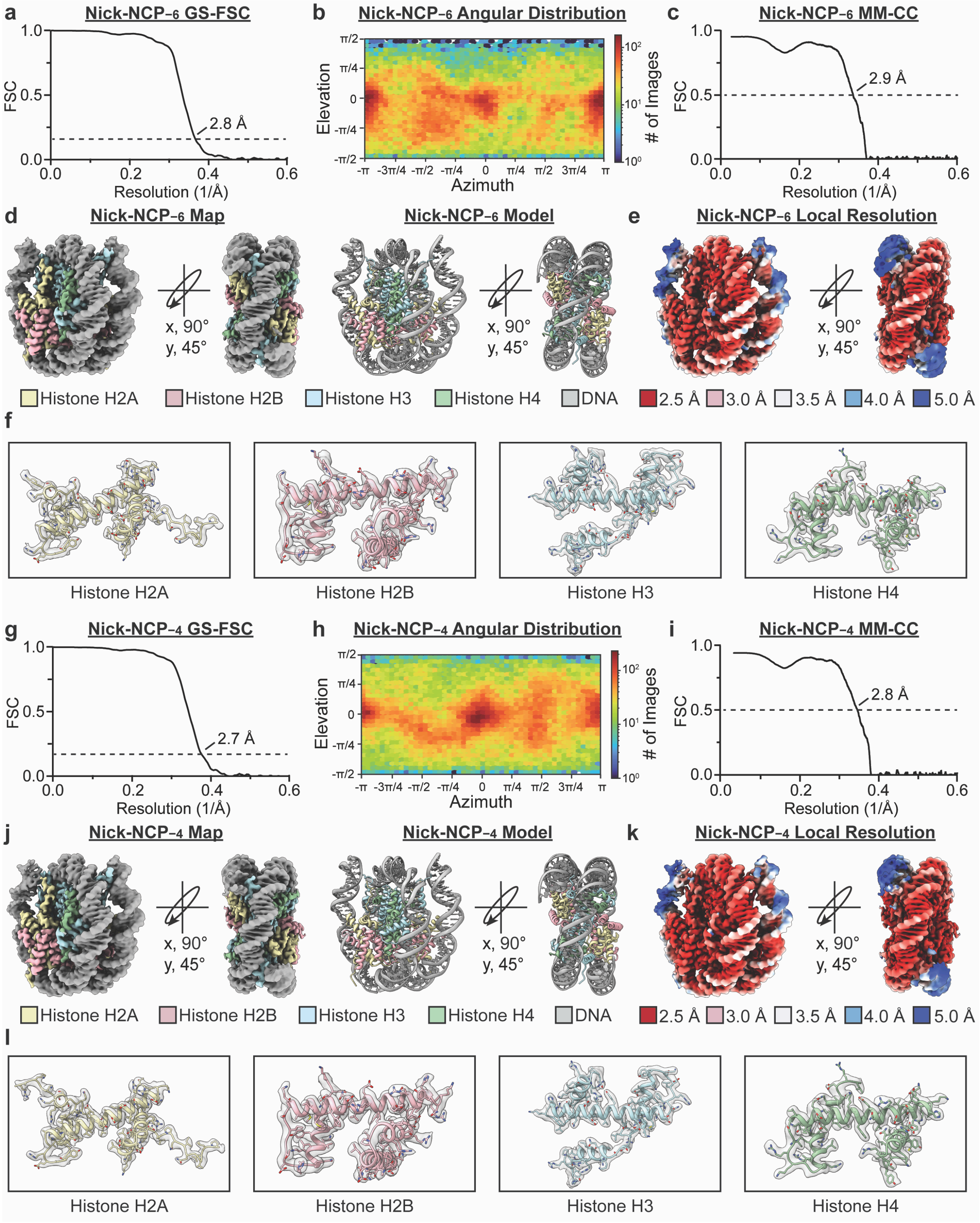

**Supplementary Fig. 10: Nick-NCP$_{-6}$ and Nick-NCP$_{-4}$ map and model quality assessment**

**a,** Gold-standard Fourier shell correlation (GS-FSC) curves for the Nick-NCP$_{-6}$ cryo-EM map (solid black line). The dashed line corresponds to GS-FSC - 0.143. **b,** Angular distribution heatmap for the Nick-NCP$_{-6}$ cryo-EM map**. c,** Map-to-model FSC curves for the Nick-NCP$_{-6}$ model and cryo-EM map. The dashed line corresponds to FSC - 0.5. **d,** The final 2.8 Å Nick-NCP$_{-6}$ cryo-EM map and model shown in two different orientations. **e**, Local resolution estimation for the Nick-NCP$_{-6}$ cryo-EM map shown in two different orientations. **f,** Representative segmented densities for histones H2A, H2B, H3, and H4 from the Nick-NCP$_{-6}$ cryo-EM map. The representative segmented densities from the cryo-EM map are shown as transparent gray surfaces. **g,** Gold-standard Fourier shell correlation (GS-FSC) curves for the Nick-NCP$_{-4}$ cryo-EM map (solid black line). The dashed line corresponds to GS-FSC - 0.143. **h,** Angular distribution heatmap for the Nick-NCP$_{-4}$ cryo-EM map**. i,** Map-to-model FSC curves for the Nick-NCP$_{-4}$ model and cryo-EM map. The dashed line corresponds to FSC - 0.5. **j,** The final 2.7 Å Nick-NCP$_{-4}$ cryo-EM map and model shown in two different orientations. **k**, Local resolution estimation for the Nick-NCP$_{-4}$ cryo-EM map shown in two different orientations. **l,** Representative segmented densities for histones H2A, H2B, H3, and H4 from the Nick-NCP$_{4}$ cryo-EM map. The representative segmented densities from the cryo-EM map are shown as transparent gray surfaces. All source data in this figure are provided as a Source data file.

**Supplementary Fig. 11: Nick-NCP−2 and Nick-NCP0 map and model quality assessment**

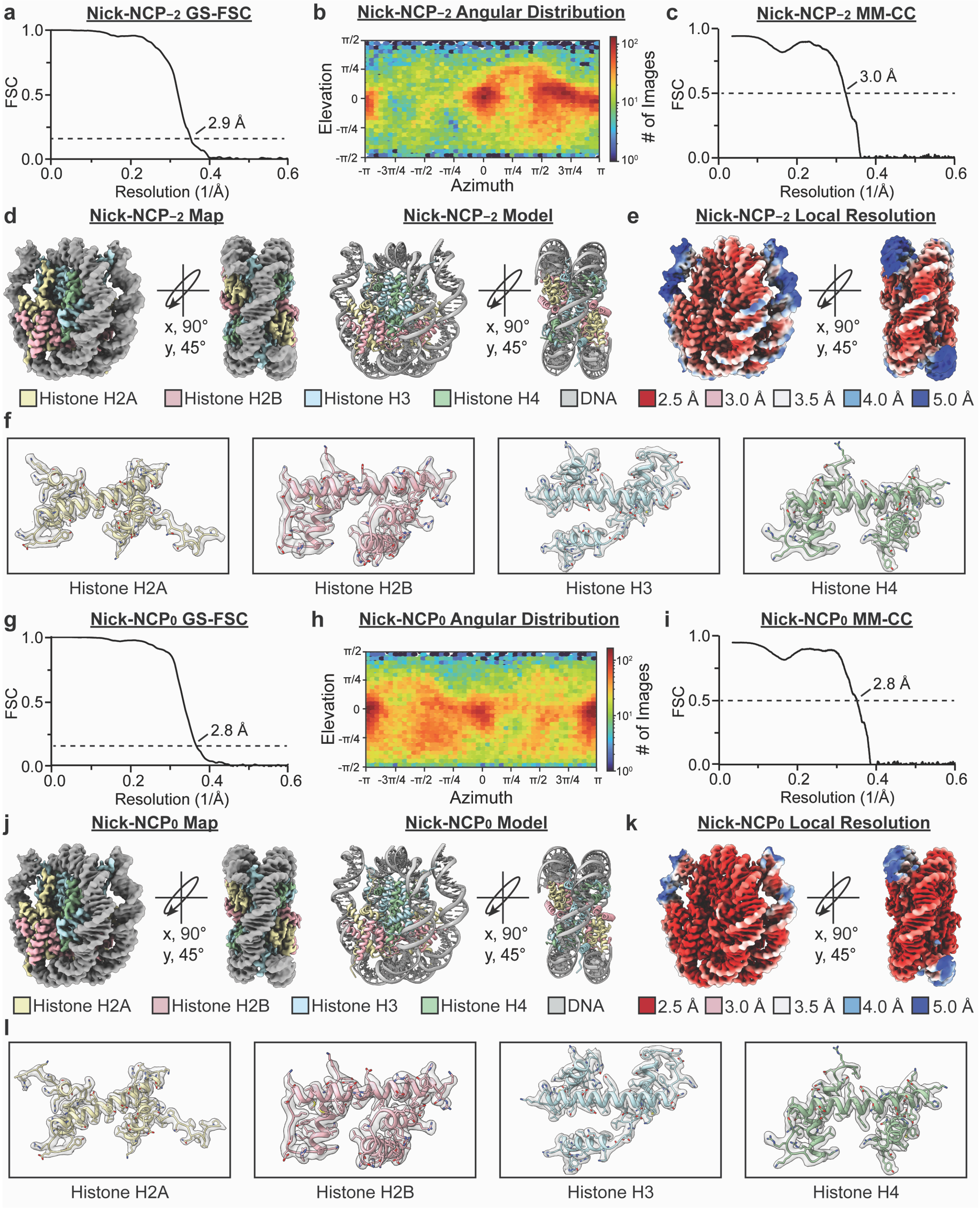

**Supplementary Fig. 11: Nick-NCP$_{-2}$ and Nick-NCP$_0$ map and model quality assessment**

**a,** Gold-standard Fourier shell correlation (GS-FSC) curves for the Nick-NCP$_{-2}$ cryo-EM map (solid black line). The dashed line corresponds to GS-FSC - 0.143. **b,** Angular distribution heatmap for the Nick-NCP$_{-2}$ cryo-EM map**.** **c,** Map-to-model FSC curves for the Nick-NCP$_{-2}$ model and cryo-EM map. The dashed line corresponds to FSC - 0.5. **d,** The final 2.9 Å Nick-NCP$_{-2}$ cryo-EM map and model shown in two different orientations. **e**, The local resolution estimation for the Nick-NCP$_{-2}$ cryo-EM map shown in two different orientations. **f,** Representative segmented densities for histones H2A, H2B, H3, and H4 from the Nick-NCP$_{-2}$ cryo-EM map. The representative segmented densities from the cryo-EM map are shown as transparent gray surfaces. **g,** Gold-standard Fourier shell correlation (GS-FSC) curves for the Nick-NCP$_0$ cryo-EM map (solid black line). The dashed line corresponds to GS-FSC - 0.143. **h,** Angular distribution heatmap for the Nick-NCP$_0$ cryo-EM map**.** **i,** Map-to-model FSC curves for the Nick-NCP$_0$ model and cryo-EM map. The dashed line corresponds to FSC - 0.5. **j,** The final 2.8 Å Nick-NCP$_0$ cryo-EM map and model shown in two different orientations. **k**, Local resolution estimation for the Nick-NCP$_0$ cryo-EM map shown in two different orientations. **l,** Representative segmented densities for histones H2A, H2B, H3, and H4 from the Nick-NCP$_0$ cryo-EM map. The representative segmented densities from the cryo-EM map are shown as transparent gray surfaces. All source data in this figure are provided as a Source data file.

**Supplementary Fig. 12: Nicks have minimal impact on nucleosome stability and structure**

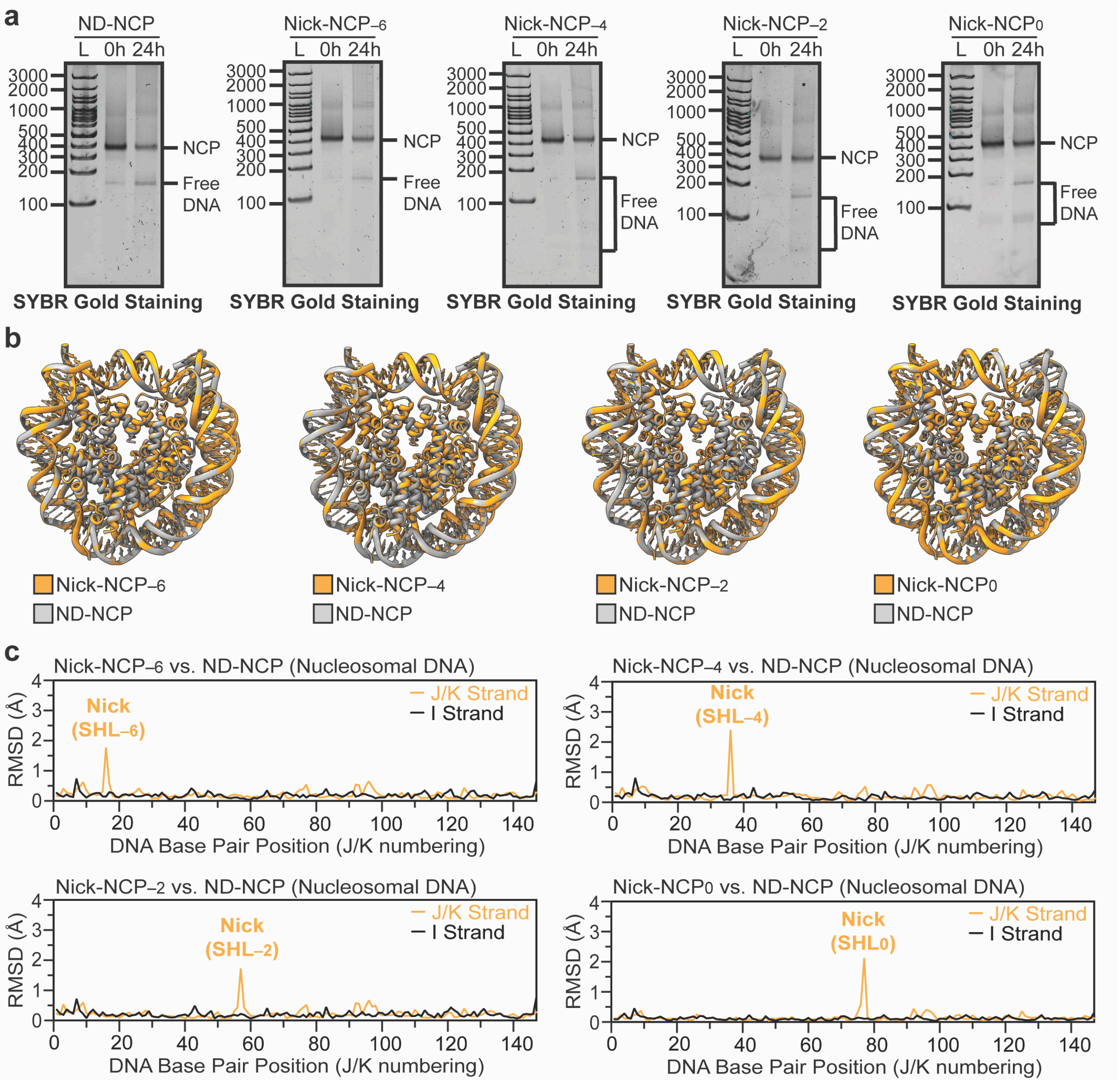

**Supplementary Fig. 12: Nicks have minimal impact on nucleosome stability and structure**

**a,** Representative native PAGE gels from the Nick-NCP$_{-6}$, Nick-NCP$_{-4}$, Nick-NCP$_{-2}$, and Nick-NCP$_{0}$ nucleosome stability assays. The NCP and free DNA species were detected using SYBR Gold staining. The gels are representative of three independent nucleosome stability assays performed for each Nick-NCP. **b,** Structural comparison of the ND-NCP with Nick-NCP$_{-6}$ (left), Nick-NCP$_{-4}$ (left middle), Nick-NCP$_{-2}$ (right middle), and Nick-NCP$_{0}$ (right). **c,** Root mean square deviation (RMSD) analysis comparing the nucleosomal DNA in the ND-NCP with the nucleosomal DNA in Nick-NCP$_{-6}$ (left), Nick-NCP$_{-4}$ (left middle), Nick-NCP$_{-2}$ (right middle), and Nick-NCP$_{0}$ (right). The location of the nick within the nucleosomal DNA is labeled in orange. All source data in this figure are provided as a Source data file.

**Supplementary Fig. 13: Nicks have minimal impact on fast timescale nucleosome dynamics**

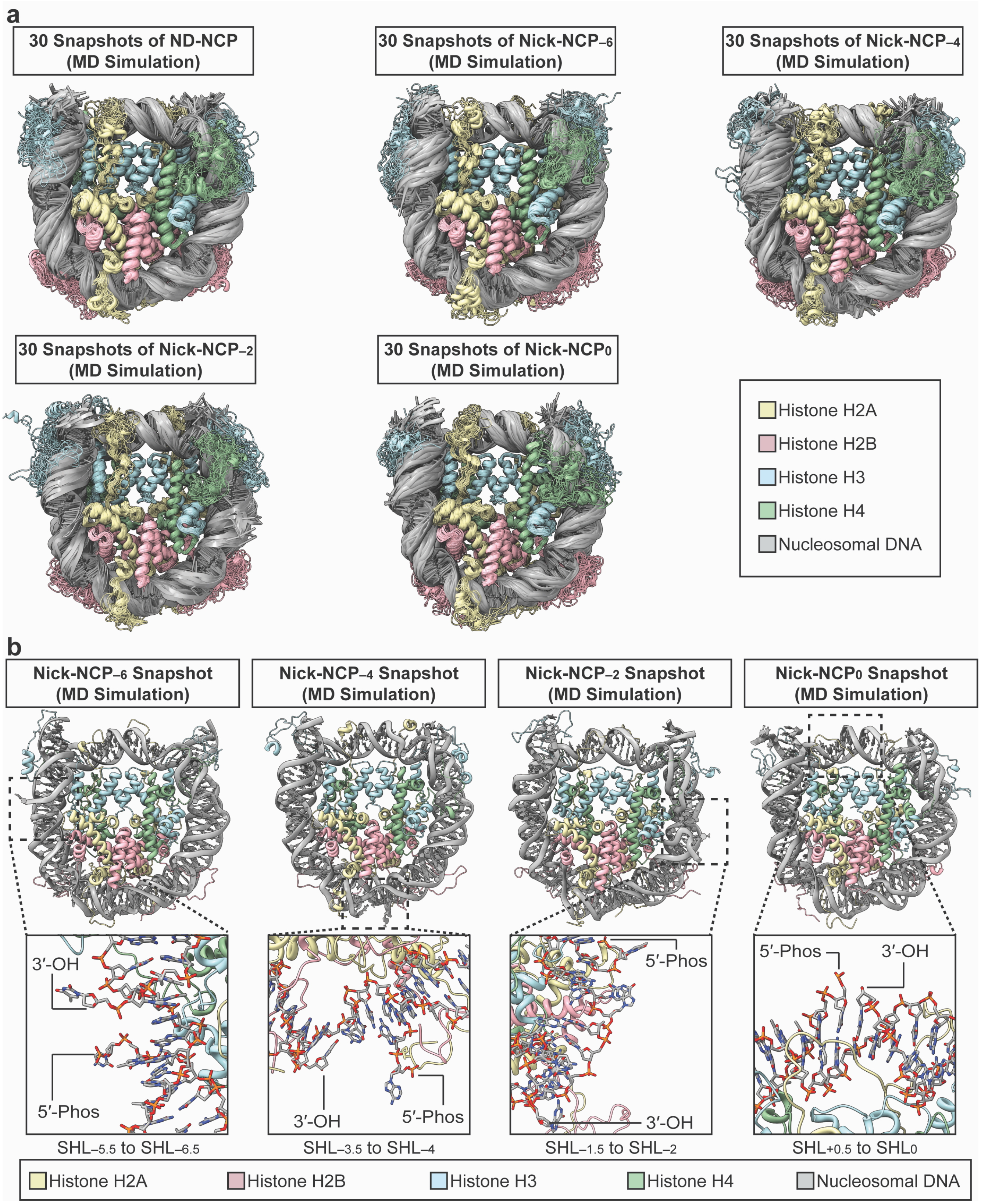

**Supplementary Fig. 13: Nicks have minimal impact on fast timescale nucleosome dynamics**

**a,** Overlay of 30 structural snapshots equally spaced throughout the 1 μs MD simulation for ND-NCP (top left), Nick-NCP$_{-6}$ (top middle), Nick-NCP$_{-4}$ (top right), Nick-NCP$_{-2}$ (bottom left), and Nick-NCP$_{0}$ (bottom right). **b,** A structural snapshot of Nick-NCP$_{-6}$ (left), Nick-NCP$_{-4}$ (middle left), Nick-NCP$_{-2}$ (middle right), and Nick-NCP$_{0}$ (right) from the 1 μs MD simulations with a with a focused view of the nick site shown as an inset. All source data in this figure are provided as a Source data file.

**Supplementary Fig. 14: LigIIIα-Nick-NCP−6 map and model quality assessment**

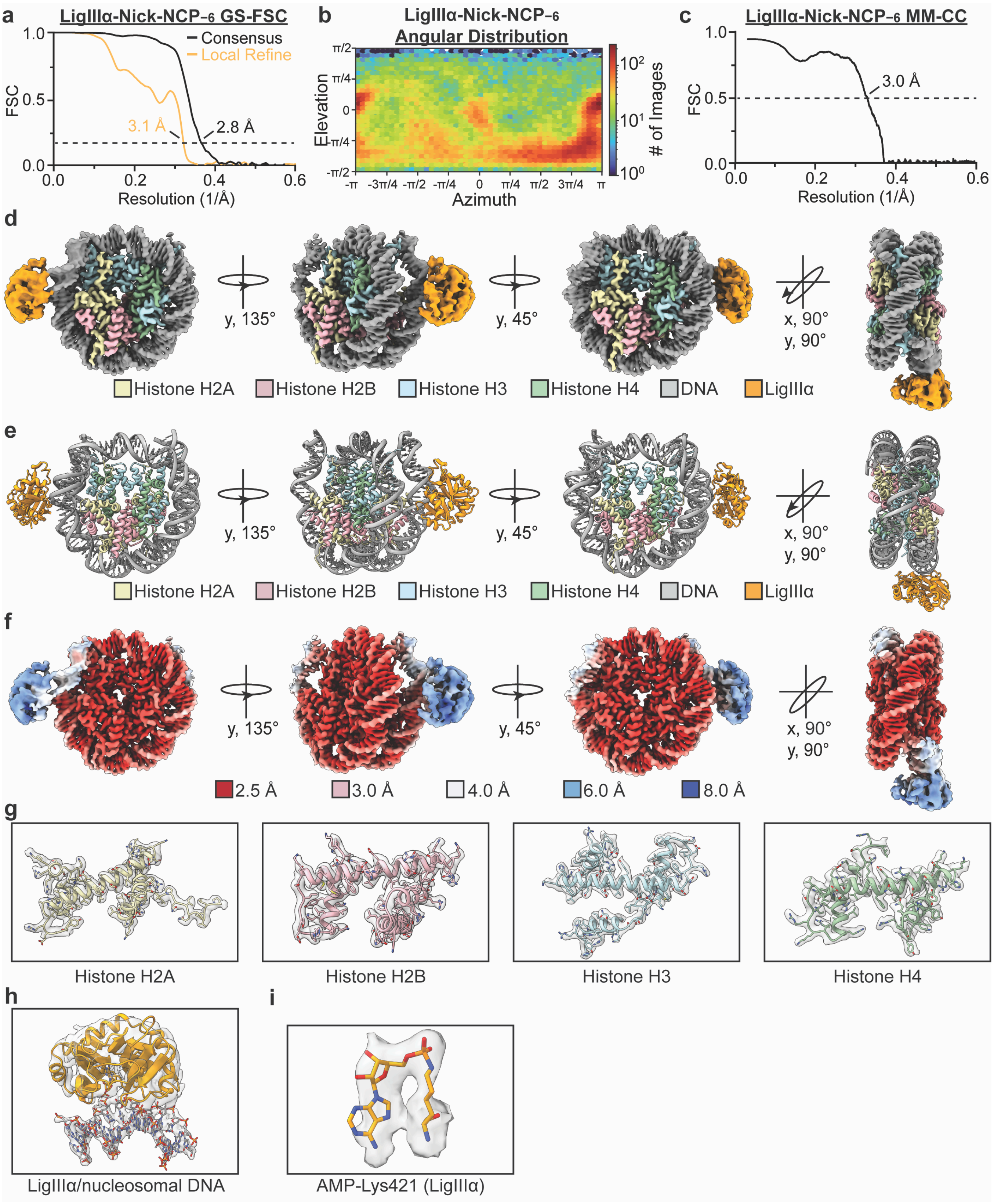

**Supplementary Fig. 14: LigIIIα-Nick-NCP$_{-6}$ map and model quality assessment**

**a,** Gold-standard Fourier shell correlation (GS-FSC) curves for the LigIIIα-Nick-NCP$_{-6}$ consensus cryo-EM map (solid black line) and the LigIIIα/nucleosomal DNA focus cryo-EM map (solid orange line). The dashed line corresponds to GS-FSC - 0.143. **b,** Angular distribution heatmap for the LigIIIα-Nick-NCP$_{-6}$ composite cryo-EM map **c,** Map-to-model FSC curves for the LigIIIα-Nick-NCP$_{-6}$ model and composite cryo-EM map. The dashed line corresponds to FSC - 0.5. **d,** The final 2.8 Å LigIIIα-Nick-NCP$_{-6}$ composite cryo-EM map and model shown in four different orientations. **e**, Local resolution estimation for the LigIIIα-Nick-NCP$_{-6}$ composite cryo-EM map shown in four different orientations. **f,** Representative segmented densities for histones H2A, H2B, H3, H4, the LigIIIα/nucleosomal DNA binding interface, and the adenylylated-K421 from the LigIIIα-Nick-NCP$_{-6}$ composite cryo-EM map. The representative segmented densities from the cryo-EM map are shown as transparent gray surfaces. All source data in this figure are provided as a Source data file.

**Supplementary Fig. 15: LigIIIα-Nick-NCP−4 map and model quality assessment**

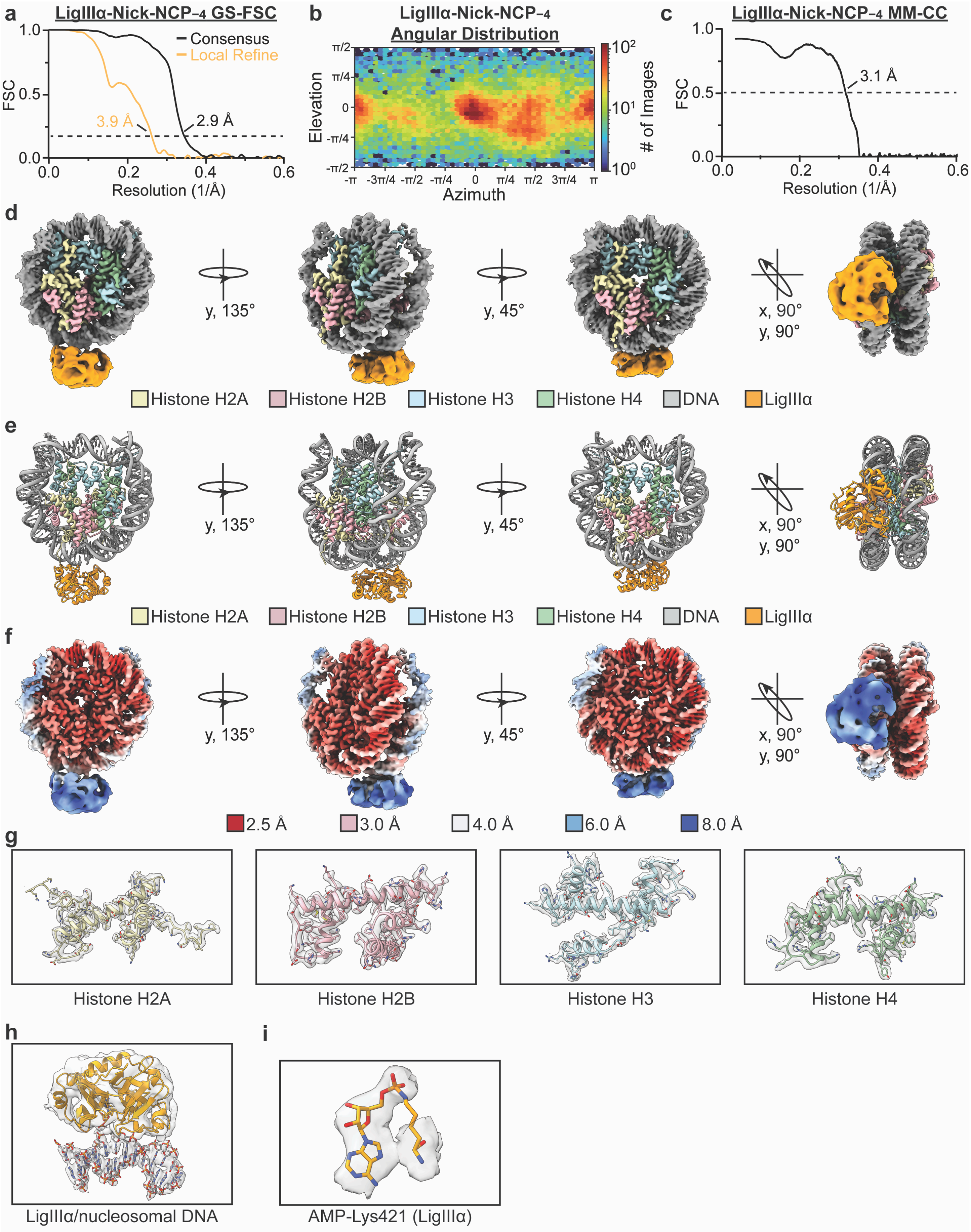

**Supplementary Fig. 15: LigIIIα-Nick-NCP−4 map and model quality assessment**

**a,** Gold-standard Fourier shell correlation (GS-FSC) curves for the LigIIIα-Nick-NCP−4 consensus cryo-EM map (solid black line) and the LigIIIα/nucleosomal DNA focus cryo-EM map (solid orange line). The dashed line corresponds to GS-FSC - 0.143. **b,** Angular distribution heatmap for the LigIIIα-Nick-NCP−4 composite cryo-EM map **c,** Map-to-model FSC curves for the LigIIIα-Nick-NCP−4 model and composite cryo-EM map. The dashed line corresponds to FSC - 0.5. **d,** The final 2.9 Å LigIIIα-Nick-NCP−4 composite cryo-EM map and model shown in four different orientations. **e**, Local resolution estimation for the LigIIIα-Nick-NCP−4 composite cryo-EM map shown in four different orientations. **f,** Representative segmented densities for histones H2A, H2B, H3, H4, the LigIIIα/nucleosomal DNA binding interface, and the adenylylated-K421 from the LigIIIα-Nick-NCP−4 composite cryo-EM map. The representative segmented densities from the cryo-EM map are shown as transparent gray surfaces. All source data in this figure are provided as a Source data file.

**Supplementary Fig. 16: LigIIIα-Nick-NCP−2 map and model quality assessment**

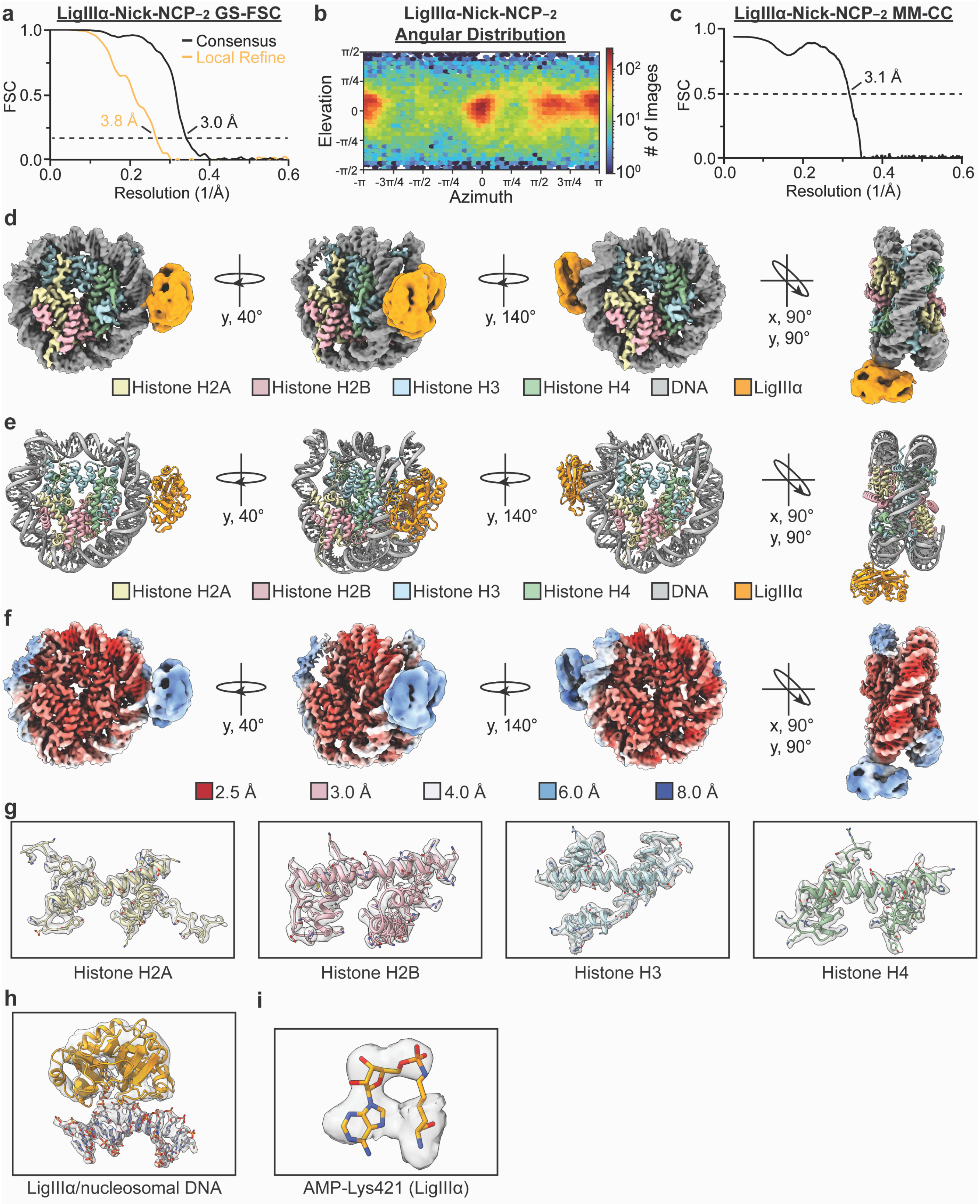

**Supplementary Fig. 16: LigIIIα-Nick-NCP−2 map and model quality assessment**

**a,** Gold-standard Fourier shell correlation (GS-FSC) curves for the LigIIIα-Nick-NCP−2 consensus cryo-EM map (solid black line) and the LigIIIα/nucleosomal DNA focus cryo-EM map (solid orange line). The dashed line corresponds to GS-FSC - 0.143. **b,** Angular distribution heatmap for the LigIIIα-Nick-NCP−2 composite cryo-EM map **c,** Map-to-model FSC curves for the LigIIIα-Nick-NCP−2 model and composite cryo-EM map. The dashed line corresponds to FSC - 0.5. **d,** The final 3.0 Å LigIIIα-Nick-NCP−2 composite cryo-EM map and model shown in four different orientations. **e**, Local resolution estimation for the LigIIIα-Nick-NCP−2 composite cryo-EM map shown in four different orientations. **f,** Representative segmented densities for histones H2A, H2B, H3, H4, the LigIIIα/nucleosomal DNA binding interface, and the adenylylated-K421 from the LigIIIα-Nick-NCP−2 composite cryo-EM map. The representative segmented densities from the cryo-EM map are shown as transparent gray surfaces. All source data in this figure are provided as a Source data file.

**Supplementary Fig. 17: LigIIIα-Nick-NCP0 map and model quality assessment**

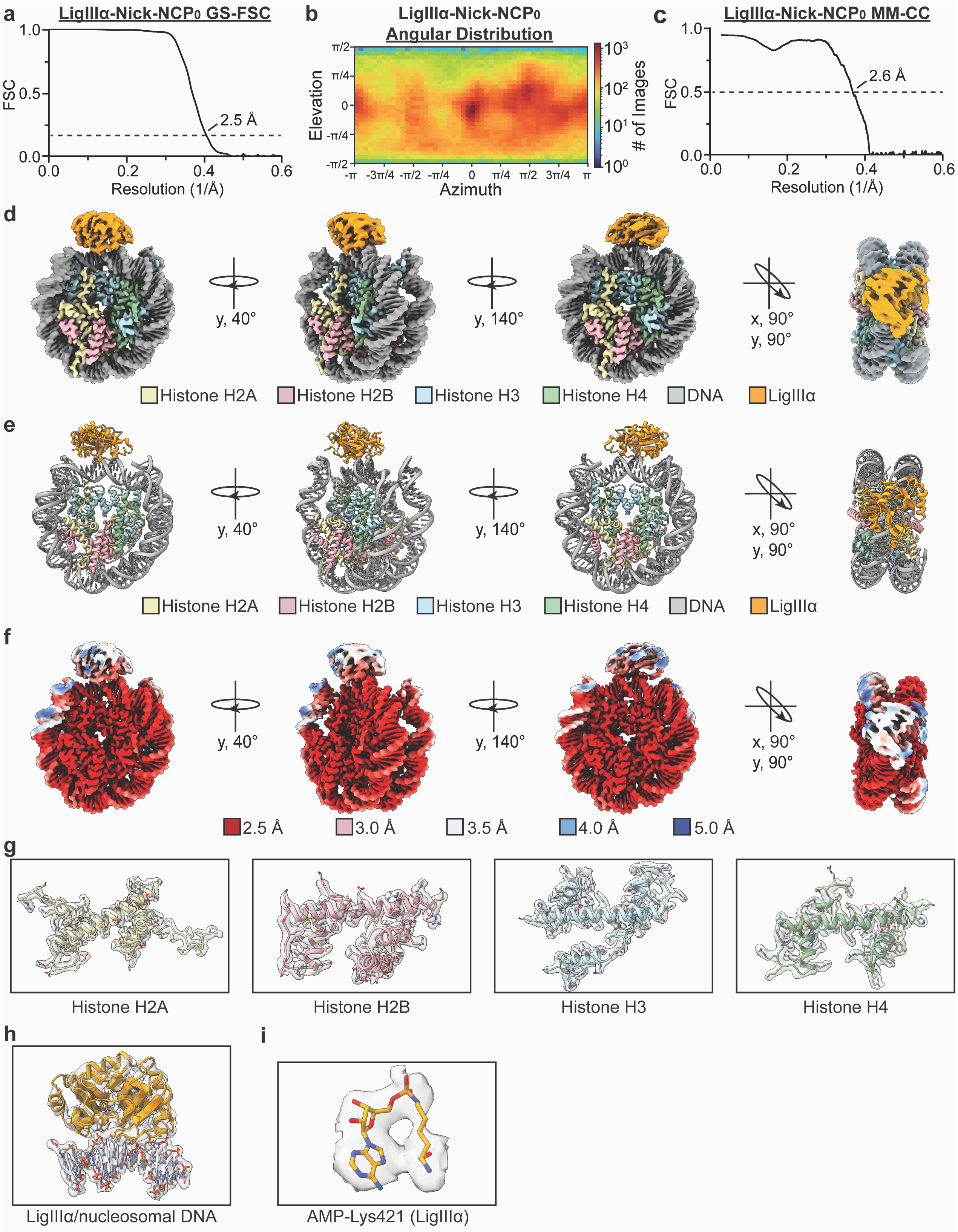

**Supplementary Fig. 17: LigIIIα-Nick-NCP0 map and model quality assessment**

**a,** Gold-standard Fourier shell correlation (GS-FSC) curves for the LigIIIα-Nick-NCP0 cryo-EM map (solid black line). The dashed line corresponds to GS-FSC - 0.143. **b,** Angular distribution heatmap for the LigIIIα-Nick-NCP0 cryo-EM map**. c,** Map-to-model FSC curves for the LigIIIα-Nick-NCP0 model and cryo-EM map. The dashed line corresponds to FSC - 0.5. **d,** The final 2.5 Å LigIIIα-Nick-NCP0 cryo-EM map and model shown in four different orientations. **e**, Local resolution estimation for the LigIIIα-Nick-NCP0 cryo-EM map shown in four different orientations. **f,** Representative segmented densities for histones H2A, H2B, H3, H4, the LigIIIα/nucleosomal DNA binding interface, and the adenylylated-K421 from the LigIIIα-Nick-NCP0 cryo-EM map. The representative segmented densities from the cryo-EM map are shown as transparent gray surfaces. All source data in this figure are provided as a Source data file.

**Supplementary Fig. 18: Unassigned density in the LigIIIα-Nick-NCP cryo-EM reconstructions**

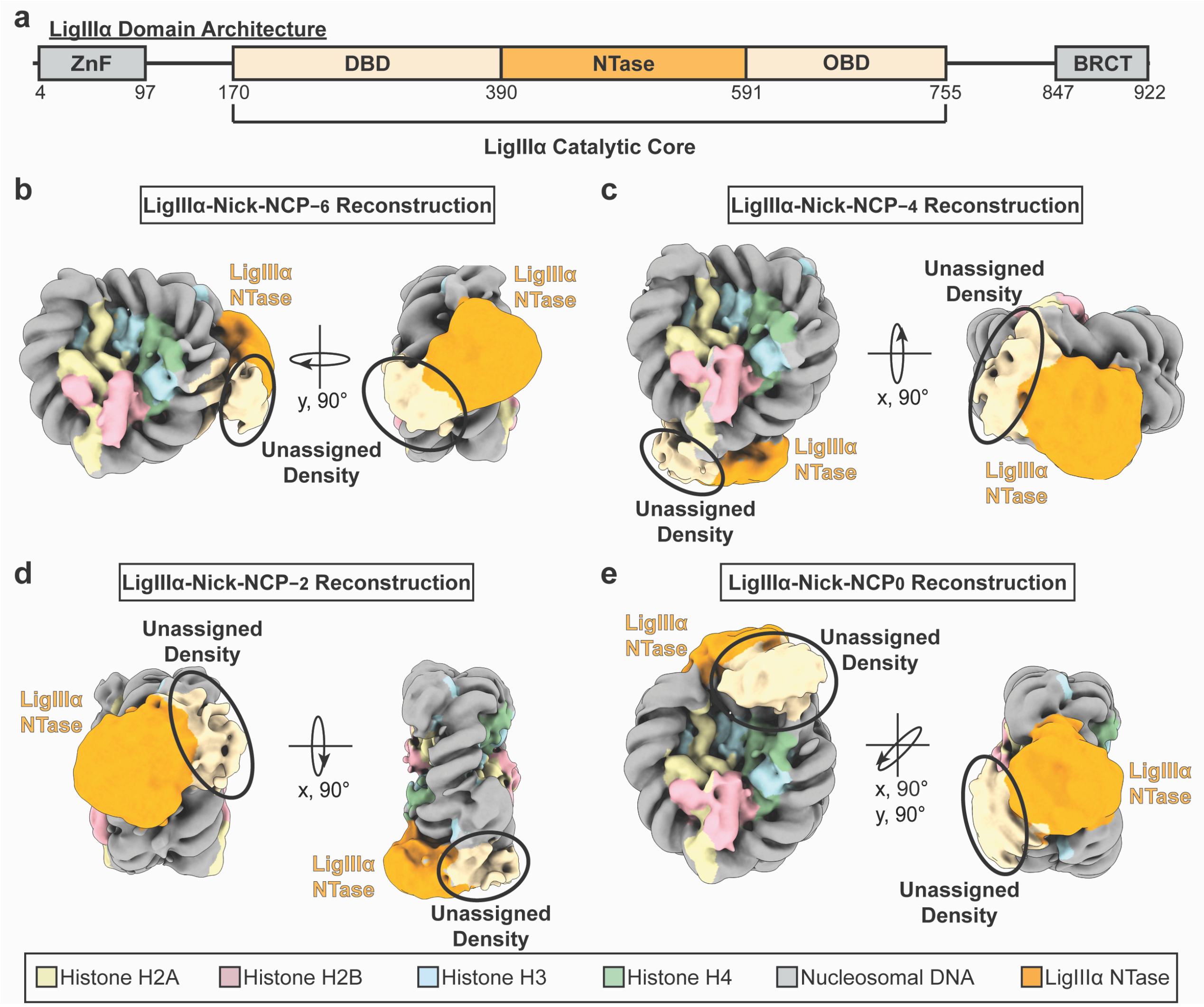

**Supplementary Fig. 18: Unassigned density in the LigIIIα-Nick-NCP cryo-EM reconstructions**

**a,** Diagram showing the domain architecture of LigIIIα. **b,** Low-pass filtered composite cryo-EM map of the LigIIIα-Nick-NCP$_{-6}$ complex. **c,** Low-pass filtered composite cryo-EM map of the LigIIIα-Nick-NCP$_{-4}$ complex. **d,** Low-pass filtered composite cryo-EM map of the LigIIIα-Nick-NCP$_{-2}$ complex. **e,** Low-pass filtered cryo-EM map of the LigIIIα-Nick-NCP$_{0}$ complex. For **b-e**, the density corresponding to the LigIIIα NTase domain is colored orange and the unassigned density is colored light yellow. All source data in this figure are provided as a Source data file.

**Supplementary Fig. 19: Structural comparison of the LigIIIα/nucleosomal DNA binding interface at each nick position in the nucleosome**

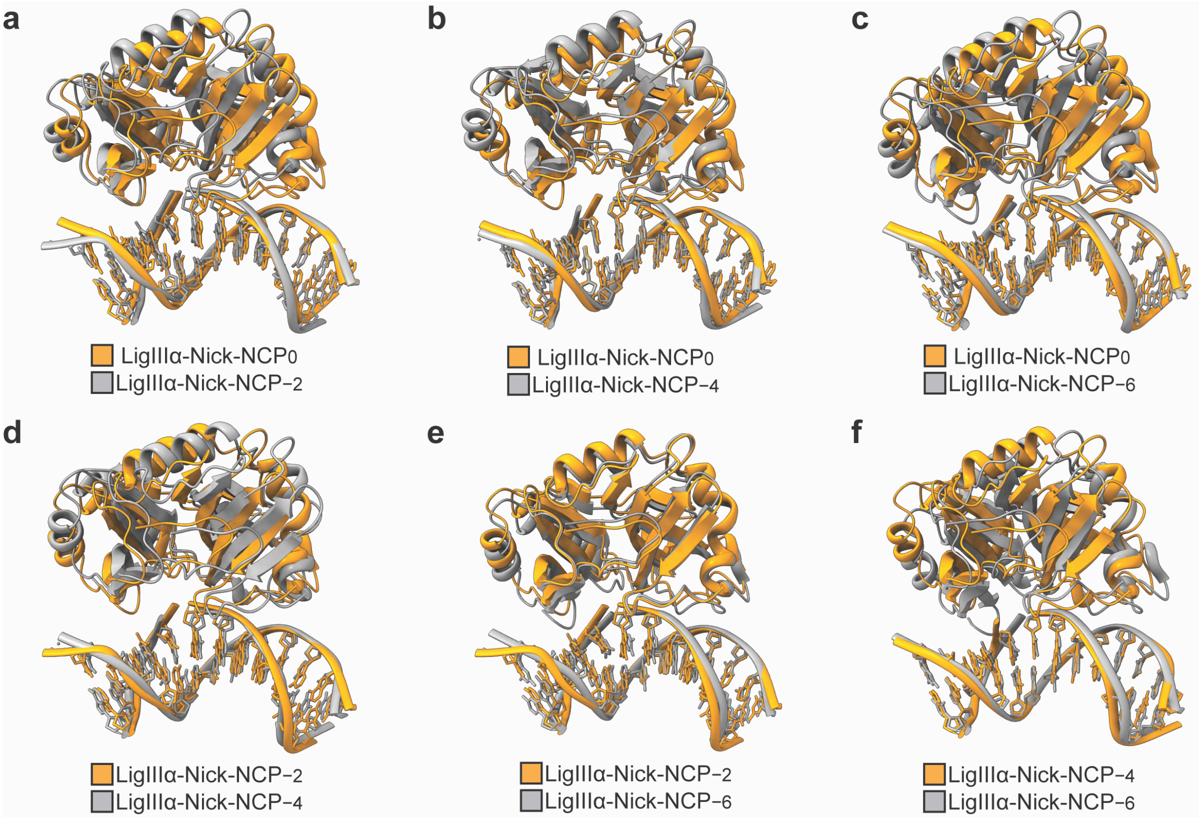

**Supplementary Fig. 19: Structural comparison of the LigIIIα/nucleosomal DNA binding interface at each nick position in the nucleosome**

**a,** Structural comparison of the LigIIIα/nucleosomal DNA binding interface in the LigIIIα-Nick-NCP$_{0}$ (orange) and LigIIIα-Nick-NCP$_{-2}$ (gray) complexes. **b,** Structural comparison of the LigIIIα/nucleosomal DNA binding interface in the LigIIIα-Nick-NCP$_{0}$ (orange) and LigIIIα-Nick-NCP$_{-4}$ (gray) complexes. **c,** Structural comparison of the LigIIIα/nucleosomal DNA binding interface in the LigIIIα-Nick-NCP$_{0}$ (orange) and LigIIIα-Nick-NCP$_{-6}$ (gray) complexes. **d,** Structural comparison of the LigIIIα/nucleosomal DNA binding interface in the LigIIIα-Nick-NCP$_{-2}$ (orange) and LigIIIα-Nick-NCP$_{-4}$ (gray) complexes. **e,** Structural comparison of the LigIIIα/nucleosomal DNA binding interface in the LigIIIα-Nick-NCP$_{-2}$ (orange) and LigIIIα-Nick-NCP$_{-6}$ (gray) complexes. **f,** Structural comparison of the LigIIIα/nucleosomal DNA binding interface in the LigIIIα-Nick-NCP$_{-4}$ (orange) and LigIIIα-Nick-NCP$_{-6}$ (gray) complexes. All source data in this figure are provided as a Source data file.

**Supplementary Fig. 20: LigIIIα adopts a catalytically inactive conformation during nick recognition in the nucleosome.**

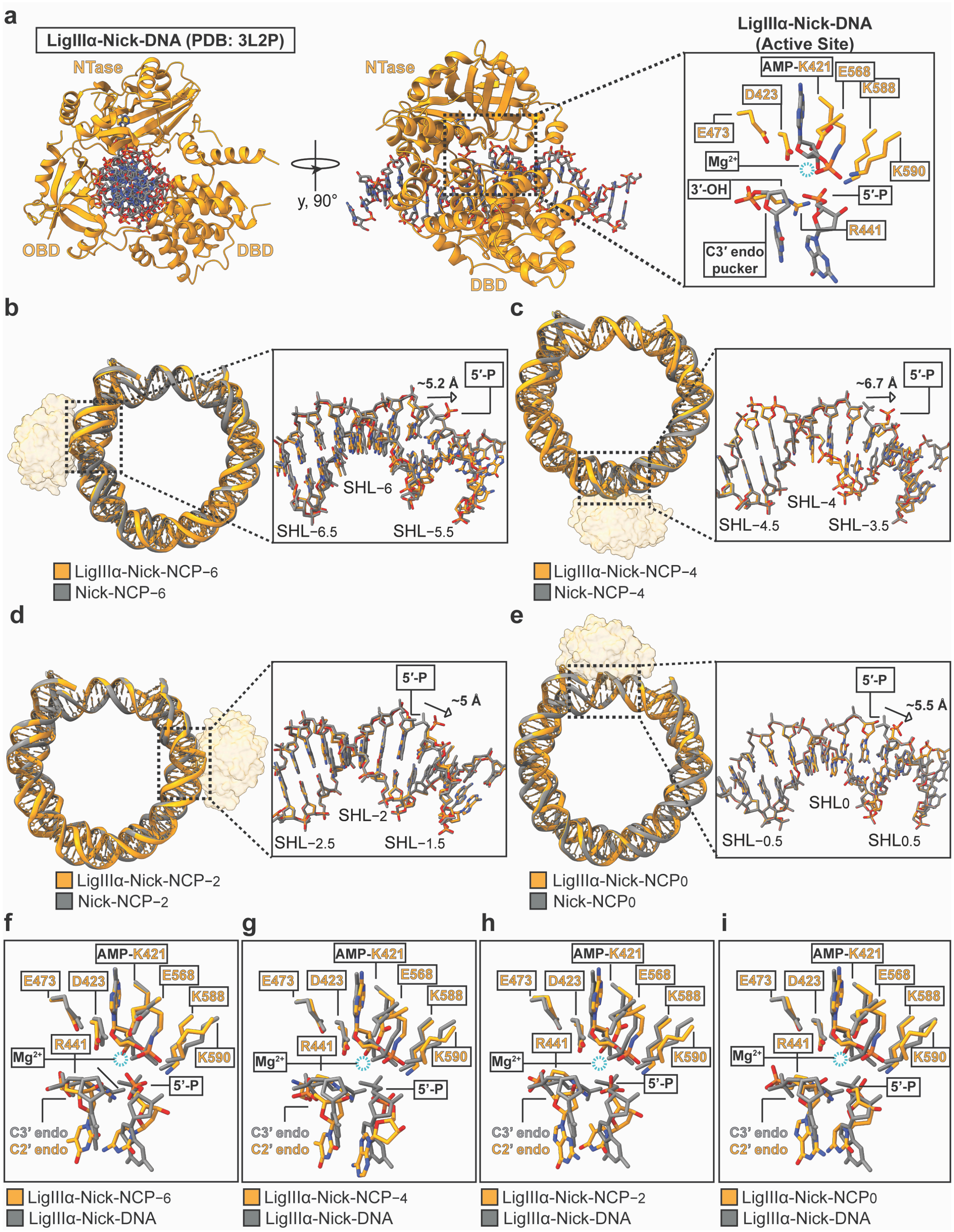

**Supplementary Fig. 20: LigIIIα adopts a catalytically inactive conformation during nick recognition in the nucleosome.**

**a,** A model of the LigIIIα-Nick-DNA complex (PDB: 3L2P) shown in two orientations, with a focused view of the LigIIIα-active site shown as an inset. The unoccupied $Mg^{2+}$ binding site is denoted with a dashed blue circle. **b,** Structural comparison of the nucleosomal DNA in the Nick-NCP$_{-6}$ (gray) and LigIIIα-Nick-NCP$_{-6}$ (orange) structures. The inset is rotated 180°. **c,** Structural comparison of the nucleosomal DNA in the Nick-NCP$_{-4}$ (gray) and LigIIIα-Nick-NCP$_{-4}$ (orange) structures. The inset is rotated 180°. **d,** Structural comparison of the nucleosomal DNA in the Nick-NCP$_{-2}$ (gray) and LigIIIα-Nick-NCP$_{-2}$ (orange) structures. The inset is rotated 180°. **e,** Structural comparison of the nucleosomal DNA in the Nick-NCP$_{0}$ (gray) and LigIIIα-Nick-NCP$_{0}$ (orange) structures. The inset is rotated 180°. In **b-e**, LigIIIα is shown as a transparent orange surface. **f,** Focused view of the LigIIIα active site in the LigIIIα-Nick-DNA (gray) and LigIIIα-Nick-NCP$_{-6}$ complex (orange). **g,** Focused view of the LigIIIα active site in the LigIIIα-Nick-DNA (gray) and LigIIIα-Nick-NCP$_{-4}$ complex (orange). **h,** Focused view of the LigIIIα active site in the LigIIIα-Nick-DNA (gray) and LigIIIα-Nick-NCP$_{-2}$ complex (orange). **i,** Focused view of the LigIIIα active site in the LigIIIα-Nick-DNA (gray) and LigIIIα-Nick-NCP$_{0}$ complex (orange). In **f-i**, the unoccupied $Mg^{2+}$ binding site is denoted with a dashed blue circle. All source data in this figure are provided as a Source data file.

**Supplementary Fig. 21: Single-turnover kinetic analysis of XRCC1-LigIIIα nick ligation in the nucleosome**

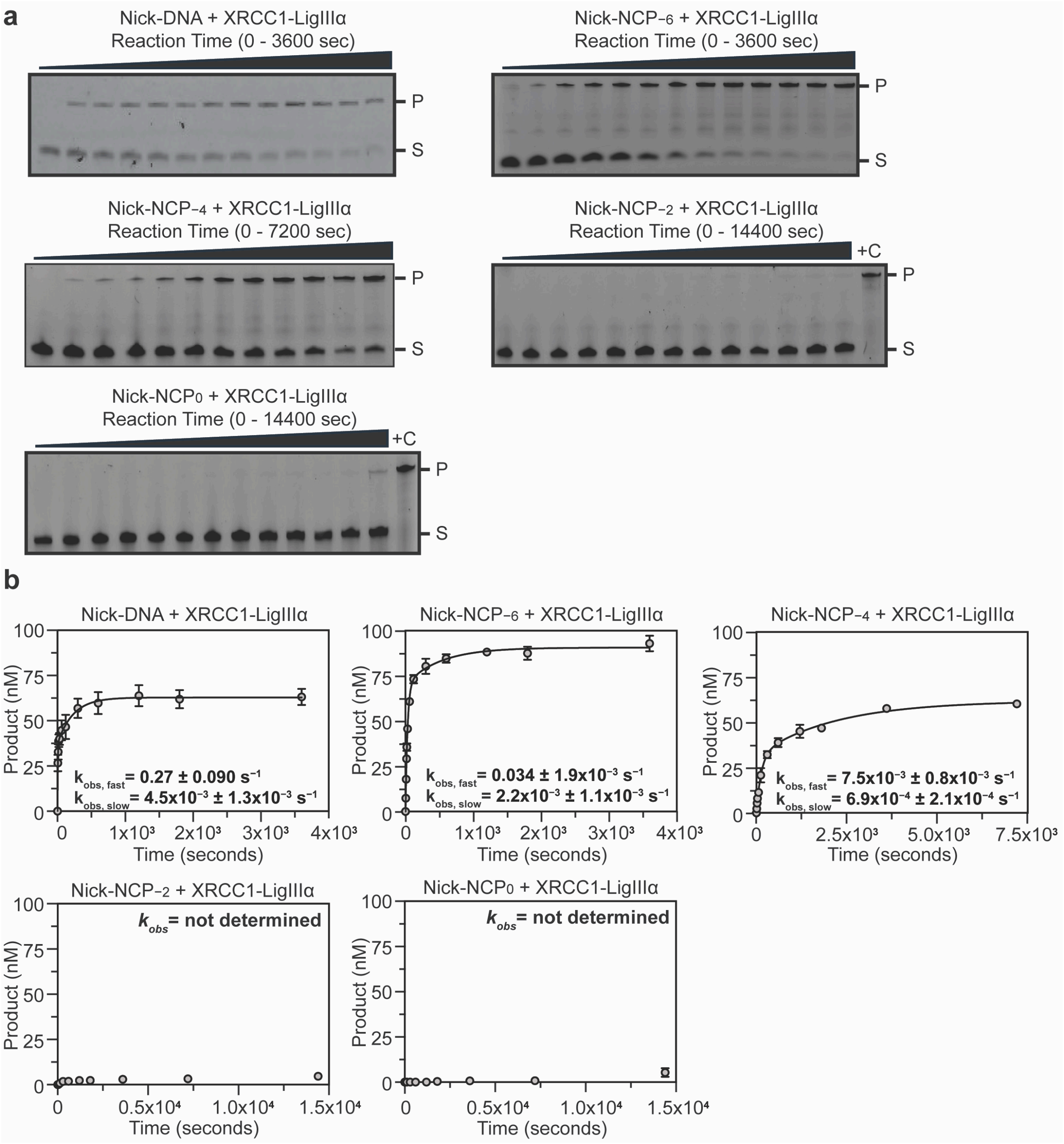

**Supplementary Fig. 21: Single-turnover kinetic analysis of XRCC1-LigIIIα nick ligation in the nucleosome**

**a,** Representative denaturing Urea-PAGE gels from the single-turnover kinetic experiments (STK) for XRCC1-LigIIIα with Nick-DNA, Nick-NCP$_0$, Nick-NCP$_{-2}$, Nick-NCP$_{-4}$, and Nick-NCP$_{-6}$. The substrate (S, Nick-DNA or Nick-NCP) and product (P, ligated DNA or NCP) were detected using the 6-FAM label on the nucleosomal DNA of each NCP. A 147 nt single-stranded DNA was loaded as a positive control (+C) for experiments with less than 10% product formed over the kinetic time course. The gels are representative of three independent STK experiments performed for XRCC1-LigIIIα with the Nick-DNA and each Nick-NCP. **b,** Quantification of the STK experiments for XRCC1-LigIIIα with Nick-DNA, Nick-NCP$_{-6}$, Nick-NCP$_{-4}$, Nick-NCP$_{-2}$, and Nick-NCP$_0$. The data points represent the mean ± standard deviation from three independent replicate experiments. The error bars are included for all experimental data points, but some error bars are smaller than the circles used to represent data points. The ligation rate ($k_{obs}$) is shown as an inset for each experiment and represents the mean ± standard error of the mean from the three independent replicate experiments. The ligation rate ($k_{obs}$) was not determined for experiments with less than 10% product formed over the kinetic time course. All source data in this figure are provided as a Source Data file.

**Supplementary Fig. 22: EMSA Analysis of the XRCC1-LigIIIα-Nick-NCP interaction**

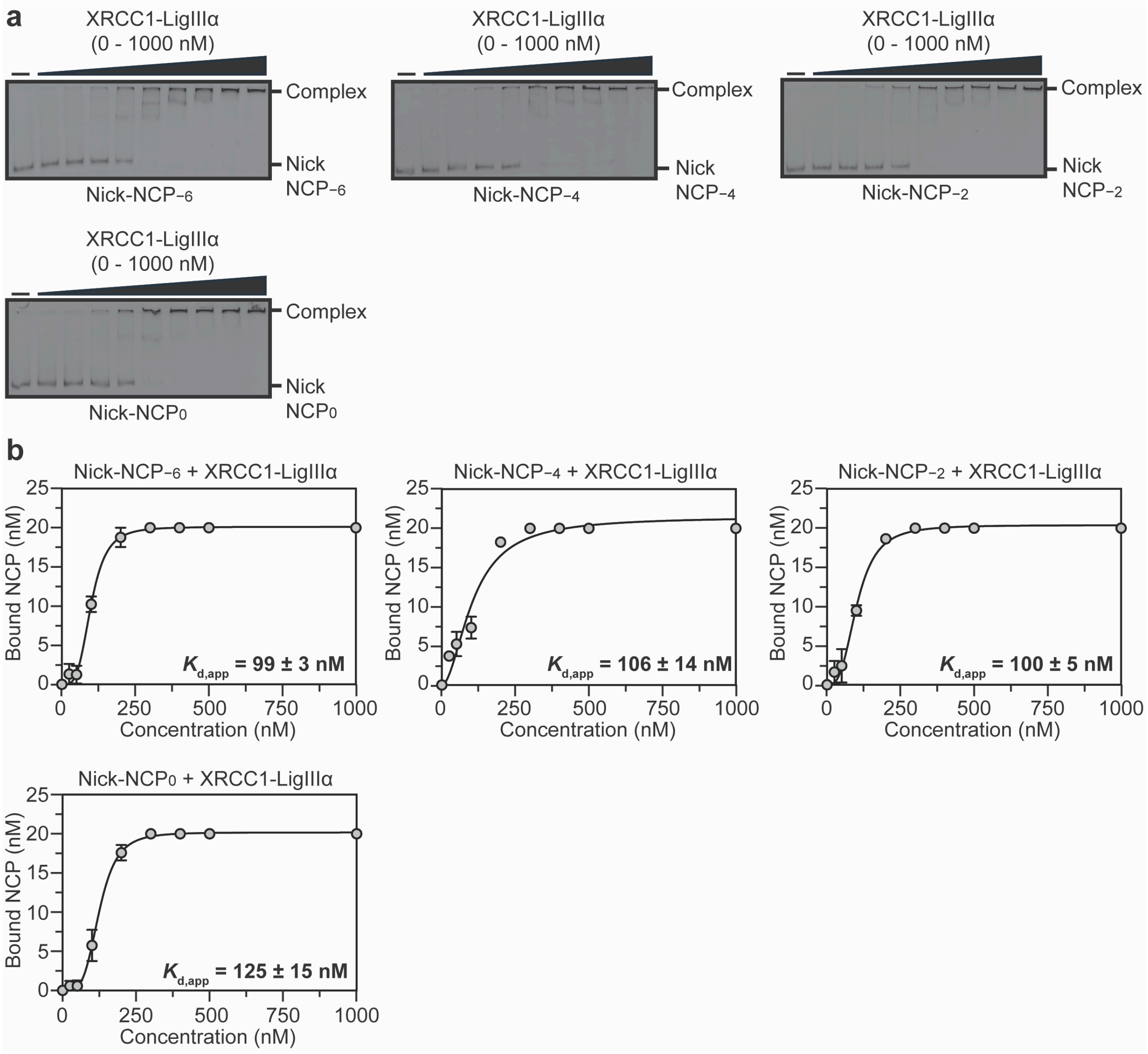

**Supplementary Fig. 22: EMSA Analysis of the XRCC1-LigIIIα-Nick-NCP interaction**

**a,** Representative native PAGE gels from the electrophoretic mobility shift assays (EMSAs) of XRCC1-LigIIIα with Nick-NCP$_{-6}$, Nick-NCP$_{-4}$, Nick-NCP$_{-2}$, and Nick-NCP$_{0}$. The free Nick-NCP and XRCC1-LigIIIα-Nick-NCP complex were detected using the 6-FAM label on the nucleosomal DNA of each NCP. The gels are representative of three independent EMSA experiments performed for XRCC1-LigIIIα and each Nick-NCP. **b,** Quantification of the EMSA experiments for XRCC1-LigIIIα with Nick-NCP$_{0}$, Nick-NCP$_{-2}$, Nick-NCP$_{-4}$, and Nick-NCP$_{-6}$. The data points represent the mean ± standard deviation from the three independent replicate experiments. The error bars are included for all experimental data points, but some error bars are smaller than the circles used to represent data points. The apparent binding affinity ($K_{d,app}$) is shown as an inset for each experiment and represents the mean ± standard deviation from the three independent replicate experiments. All source data in this figure are provided as a Source Data file.

**Supplementary Fig. 23: XRCC1-LigIIIα-Nick-NCP-6 single particle analysis processing workflow**

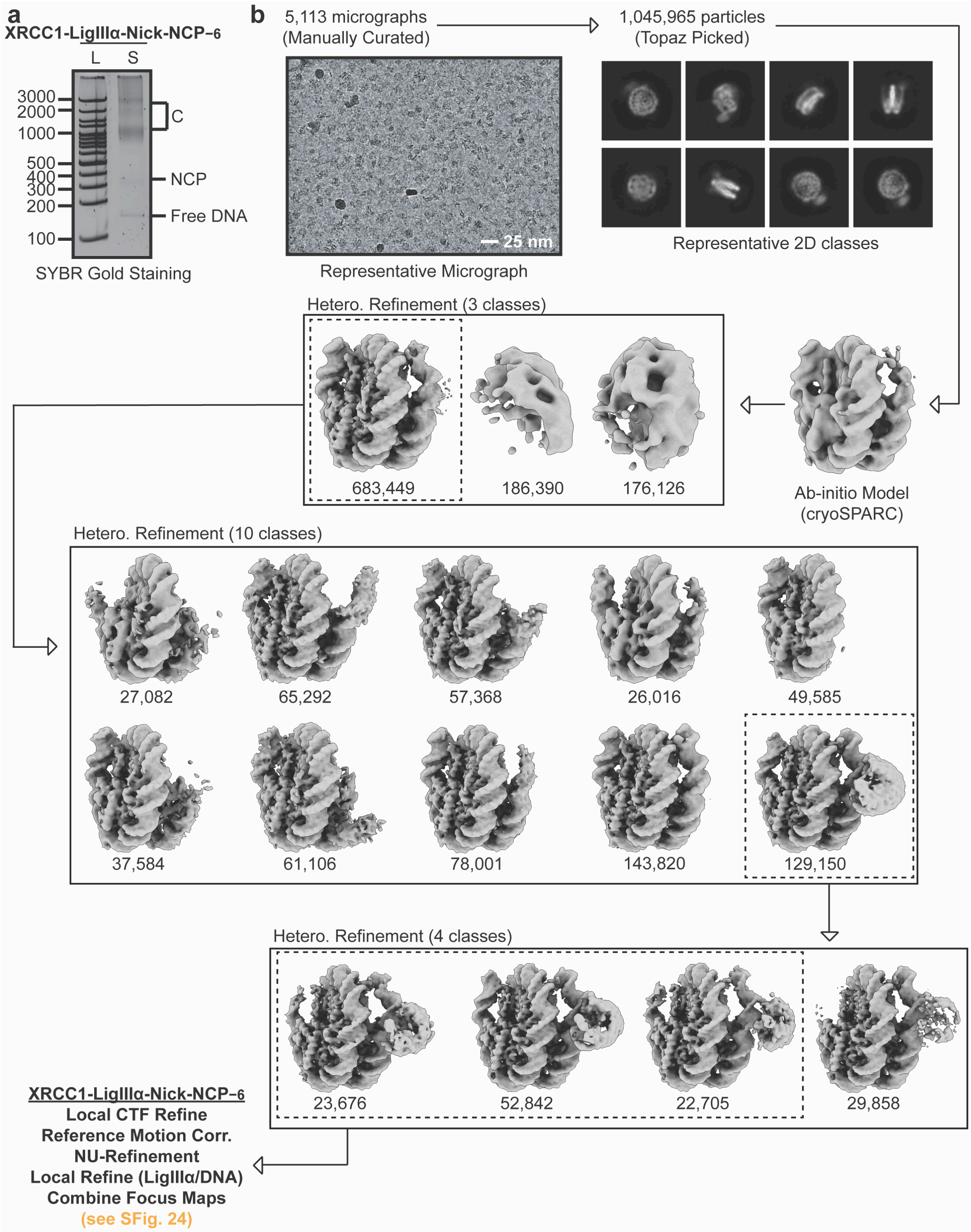

**Supplementary Fig. 23: XRCC1-LigIIIα-Nick-NCP$_{-6}$ single particle analysis processing workflow**

**a,** Native PAGE gel of the XRCC1-LigIIIα-Nick-NCP$_{-6}$ cryo-EM sample (S) and a 100 bp DNA ladder (L). The NCP$_{-6}$ and XRCC1-LigIIIα-Nick-NCP$_{-6}$ complex were visualized with SYBR gold staining. The bands corresponding to free DNA, the NCP, and XRCC1-LigIIIα-Nick-NCP$_{-6}$ (C) are labeled. **b,** Cryo-EM data processing workflow for the XRCC1-LigIIIα-Nick-NCP$_{-6}$ dataset. A representative micrograph (n=5,113) and representative 2D classes from the XRCC1-LigIIIα-Nick-NCP$_{-6}$ dataset are shown. The maps chosen for further classification and/or refinement throughout the data processing pipeline are boxed. The final map, final model, and quality assessment metrics for XRCC1-LigIIIα-Nick-NCP$_{-6}$ can be found in Supplementary Fig. 24. All source data in this figure are provided as a Source data file.

### Supplementary Fig. 24: XRCC1-LigIIIα-Nick-NCP−6 map and model quality assessment

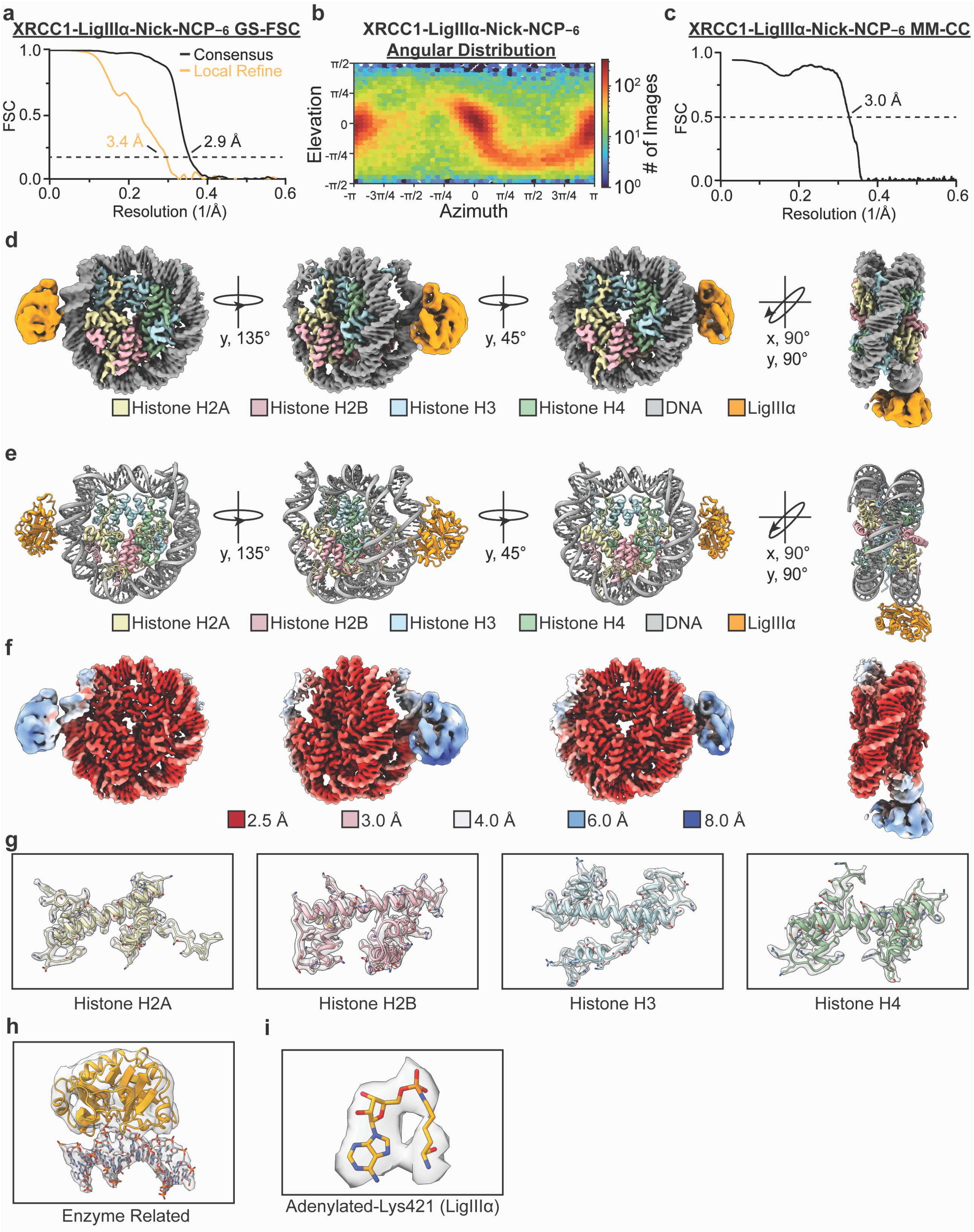

**Supplementary Fig. 24: XRCC1-LigIIIα-Nick-NCP$_{-6}$ map and model quality assessment**

**a,** Gold-standard Fourier shell correlation (GS-FSC) curves for the XRCC1-LigIIIα-Nick-NCP$_{-6}$ consensus cryo-EM map (solid black line) and the LigIIIα/nucleosomal DNA focus cryo-EM map (solid orange line). The dashed line corresponds to GS-FSC - 0.143. **b,** Angular distribution heatmap for the XRCC1-LigIIIα-Nick-NCP$_{-6}$ composite cryo-EM map **c,** Map-to-model FSC curves for the XRCC1-LigIIIα-Nick-NCP$_{-6}$ model and composite cryo-EM map. The dashed line corresponds to FSC - 0.5. **d,** The final 2.9 Å XRCC1-LigIIIα-Nick-NCP$_{-6}$ composite cryo-EM map and model shown in four different orientations. **e**, Local resolution estimation for the XRCC1-LigIIIα-Nick-NCP$_{-6}$ composite cryo-EM map shown in four different orientations. **f,** Representative segmented densities for histones H2A, H2B, H3, H4, the LigIIIα/nucleosomal DNA binding interface, and the adenylylated-K421 from the XRCC1-LigIIIα-Nick-NCP$_{-6}$ composite cryo-EM map. The representative segmented densities from the cryo-EM map are shown as transparent gray surfaces. All source data in this figure are provided as a Source data file.

**Supplementary Table 1: LigIIIα and XRCC1-LigIIIα apparent nucleosome binding affinities and ligation rates**

| Protein | Substrate | $k_{obs}$ (fast, $s^{-1}$) | Fold Change (Nick-DNA) | $k_{obs}$ (slow, $s^{-1}$) | Fold Change (Nick-DNA) | $K_{d,app}$ (nM) | Hill Coeff. |
|---|---|---|---|---|---|---|---|
| LigIIIα | Nick-DNA | 0.21 ± 0.027 | - | $6.1\times10^{-3} \pm 2.0\times10^{-3}$ | - | - | - |
| | Nick-NCP$_{-6}$ | $0.061 \pm 5.6\times10^{-3}$ | 3.4 | $2.4\times10^{-3} \pm 1.3\times10^{-3}$ | 2.5 | 120 ± 5 | 2.7 |
| | Nick-NCP$_{-4}$ | $6.8\times10^{-3} \pm 1.8\times10^{-3}$ | 31 | $8.1\times10^{-4} \pm 3.8\times10^{-4}$ | 7.5 | 135 ± 8 | 2.6 |
| | Nick-NCP$_{-2}$ | ND | - | ND | - | 106 ± 11 | 1.5 |
| | Nick-NCP$_{0}$ | ND | - | ND | - | 134 ± 2 | 2.1 |
| XRCC1-LigIIIα | Nick-DNA | 0.27 ± 0.090 | - | $4.5\times10^{-3} \pm 1.3\times10^{-3}$ | - | - | - |
| | Nick-NCP$_{-6}$ | $0.034 \pm 1.9\times10^{-3}$ | 7.9 | $2.2\times10^{-3} \pm 1.1\times10^{-3}$ | 2 | 99 ± 3 | 3.8 |
| | Nick-NCP$_{-4}$ | $7.5\times10^{-3} \pm 0.8\times10^{-3}$ | 36 | $6.9\times10^{-4} \pm 2.1\times10^{-4}$ | 6.5 | 106 ± 14 | 1.8 |
| | Nick-NCP$_{-2}$ | ND | - | ND | - | 100 ± 5 | 3.0 |
| | Nick-NCP$_{0}$ | ND | - | ND | - | 125 ± 15 | 4.0 |

ND – Not determined

**Supplementary Table 2: Cryo-EM data collection, refinement, and validation**

| **Data collection and processing** | |
|---|---|
| **Dataset** | **ND-NCP** |
| Magnification | 130,000x |
| Voltage (kV) | 300 |
| Electron exposure ($e^{-}/Å^2$) | 50 |
| Defocus range (μm) | −1.2 to −1.8 |
| Pixel size (Å) | 0.652 |
| Symmetry imposed | C1 |
| Initial particle number | 1,475,651 |
| **Structure** | **ND-NCP** |
| Final particle number | 439,555 |
| Map resolution (Å) | 2.6 |
| FSC threshold | 0.143 |
| PDB accession | **10XZ** |
| EMDB accession | **EMD-75521** |
| **Refinement** | |
| Initial model used (PDB ID) | 9DWF |
| Model resolution (Å) | 2.6 |
| FSC threshold | 0.5 |
| **Model composition** | |
| Nonhydrogen atoms | 12,043 |
| Protein residues | 756 |
| Nucleotide | 294 |
| **B factors ($Å^2$)** | |
| Protein | 87.05 |
| Nucleotide | 135.10 |
| **r.m.s. deviations** | |
| Bond Length (Å) (# > 4σ) | 0.005 (0) |
| Bond Angles (°) (# > 4σ) | 0.803 (0) |
| **Validation** | |
| MolProbity score | 1.17 |
| Clashscore | 3.78 |
| Rotamer Outliers (%) | 0.32 |
| **Ramachandran plot** | |
| Favored (%) | 98.51 |
| Allowed (%) | 1.49 |
| Disallowed (%) | 0.00 |

**Supplementary Table 3: Cryo-EM data collection, refinement, and validation**

| **Data collection and processing** | | | | |
|---|---|---|---|---|
| **Dataset** | **Nick-NCP$_0$** | | **Nick-NCP$_{-2}$** | |
| Magnification | 130,000x | | 130,000x | |
| Voltage (kV) | 300 | | 300 | |
| Electron exposure (e$^-$/Å$^2$) | 50 | | 50 | |
| Defocus range (μm) | −1.2 to −1.8 | | −1.2 to −1.8 | |
| Pixel size (Å) | 0.652 | | 0.652 | |
| Symmetry imposed | C1 | | C1 | |
| Initial particle number | 1,476,778 | | 510,094 | |
| **Structure** | **Nick-NCP$_0$** | **LigIIIα-Nick-NCP$_0$** | **Nick-NCP$_{-2}$** | **LigIIIα-Nick-NCP$_{-2}$** |
| Final particle number | 141,040 | 463,243 | 50,482 | 50,548 |
| Map resolution (Å) | 2.8 | 2.5 | 2.9 | 3.0 |
| FSC threshold | 0.143 | 0.143 | 0.143 | 0.143 |
| PDB accession | **10YA** | **10YE** | **10YB** | **10YF** |
| EMDB accession | **EMD-75522** | **EMD-75526** | **EMD-75523** | **EMD-75527**<br>**EMD-75531**<br>**EMD-75532** |
| **Refinement** | | | | |
| Initial model used (PDB ID) | 9DWF | 9DWF, 3L2P | 9DWF | 9DWF, 3L2P |
| Model resolution (Å) | 2.8 | 2.6 | 3.0 | 3.1 |
| FSC threshold | 0.5 | 0.5 | 0.5 | 0.5 |
| **Model composition** | | | | |
| Nonhydrogen atoms | 11,998 | 13,681 | 11,904 | 13,644 |
| Protein residues | 753 | 961 | 742 | 957 |
| Nucleotide | 294 | 294 | 294 | 294 |
| **B factors (Å$^2$ )** | | | | |
| Protein | 59.75 | 110.23 | 87.64 | 99.00 |
| Nucleotide | 105.09 | 129.17 | 154.24 | 151.30 |
| **r.m.s. deviations** | | | | |
| Bond Length (Å) (# > 4σ) | 0.004 (0) | 0.005 (0) | 0.005 (0) | 0.007 (0) |
| Bond Angles (°) (# > 4σ) | 0.772 (0) | 0.802 (0) | 0.810 (1) | 0.813 (0) |
| **Validation** | | | | |
| MolProbity score | 1.19 | 1.17 | 1.16 | 1.17 |
| Clashscore | 4.08 | 3.81 | 3.69 | 3.82 |
| Rotamer Outliers (%) | 0.16 | 0.25 | 0.49 | 0.37 |
| **Ramachandran plot** | | | | |
| Favored (%) | 98.91 | 98.41 | 98.62 | 98.08 |
| Allowed (%) | 1.09 | 1.59 | 1.38 | 1.92 |
| Disallowed (%) | 0.00 | 0.00 | 0.00 | 0.00 |

**Supplementary Table 4: Cryo-EM data collection, refinement, and validation**

| **Data collection and processing** | | | | |
|---|---|---|---|---|
| **Dataset** | **Nick-NCP$_{-4}$** | | **Nick-NCP$_{-6}$** | |
| Magnification | 130,000x | | 130,000x | |
| Voltage (kV) | 300 | | 300 | |
| Electron exposure ($e^{-}$/Å$^2$) | 50 | | 50 | |
| Defocus range (μm) | −1.2 to −1.8 | | −1.2 to −1.8 | |
| Pixel size (Å) | 0.652 | | 0.652 | |
| Symmetry imposed | C1 | | C1 | |
| Initial particle number | 573,461 | | 629,703 | |
| **Structure** | **Nick-NCP$_{-4}$** | **LigIIIα-Nick-NCP$_{-4}$** | **Nick-NCP$_{-6}$** | **LigIIIα-Nick-NCP$_{-6}$** |
| Final particle number | 97,160 | 39,115 | 72,441 | 74,600 |
| Map resolution (Å) | 2.7 | 2.9 | 2.8 | 2.8 |
| FSC threshold | 0.143 | 0.143 | 0.143 | 0.143 |
| PDB accession | **10YC** | **10YG** | **10YD** | **10YH** |
| EMDB accession | **EMD-75524** | **EMD-75528**<br>**EMD-75533**<br>**EMD-75534** | **EMD-75525** | **EMD-75529**<br>**EMD-75535**<br>**EMD-75536** |
| **Refinement** | | | | |
| Initial model used (PDB ID) | 9DWF | 9DWF, 3L2P | 9DWF | 9DWF, 3L2P |
| Model resolution (Å) | 2.8 | 3.1 | 2.9 | 3.0 |
| FSC threshold | 0.5 | 0.5 | 0.5 | 0.5 |
| **Model composition** | | | | |
| Nonhydrogen atoms | 11,998 | 13,681 | 11,959 | 13,642 |
| Protein residues | 753 | 961 | 748 | 956 |
| Nucleotide | 294 | 294 | 294 | 294 |
| **B factors (Å$^2$ )** | | | | |
| Protein | 89.08 | 109.39 | 83.94 | 112.74 |
| Nucleotide | 151.07 | 159.02 | 138.59 | 142.41 |
| **r.m.s. deviations** | | | | |
| Bond Length (Å) (# > 4σ) | 0.004 (0) | 0.005 (0) | 0.005 (0) | 0.008 (0) |
| Bond Angles (°) (# > 4σ) | 0.778 (0) | 0.812 (0) | 0.785 (1) | 0.828 (0) |
| **Validation** | | | | |
| MolProbity score | 1.12 | 1.17 | 1.19 | 1.20 |
| Clashscore | 3.24 | 3.77 | 4.00 | 3.38 |
| Rotamer Outliers (%) | 0.16 | 0.37 | 0.32 | 0.25 |
| **Ramachandran plot** | | | | |
| Favored (%) | 98.24 | 97.99 | 98.22 | 97.65 |
| Allowed (%) | 1.76 | 2.01 | 1.78 | 2.35 |
| Disallowed (%) | 0.00 | 0.00 | 0.00 | 0.00 |

**Supplementary Table 5: Cryo-EM data collection, refinement, and validation**

| | |
|---|---|
| **Data collection and processing** | |
| **Dataset** | **XRCC1-LigIIIα-Nick-NCP$_{-6}$** |
| Magnification | 130,000x |
| Voltage (kV) | 300 |
| Electron exposure ($e^-$/Å$^2$) | 50 |
| Defocus range (μm) | −1.2 to −1.8 |
| Pixel size (Å) | 0.652 |
| Symmetry imposed | C1 |
| Initial particle number | 599,935 |
| **Structure** | **XRCC1-LigIIIα-Nick-NCP$_{-6}$** |
| Final particle number | 99,223 |
| Map resolution (Å) | 2.9 |
| FSC threshold | 0.143 |
| PDB accession | **10YI** |
| EMDB accession | **EMD-75530**<br>**EMD-75537**<br>**EMD-75538** |
| **Refinement** | |
| Initial model used (PDB ID) | 9DWF, 3L2P |
| Model resolution (Å) | 3.0 |
| FSC threshold | 0.5 |
| **Model composition** | |
| Nonhydrogen atoms | 13642 |
| Protein residues | 956 |
| Nucleotide | 294 |
| **B factors (Å$^2$ )** | |
| Protein | 95.17 |
| Nucleotide | 137.49 |
| **r.m.s. deviations** | |
| Bond Length (Å) (# > 4σ) | 0.005 (0) |
| Bond Angles (°) (# > 4σ) | 0.791 (0) |
| **Validation** | |
| MolProbity score | 1.25 |
| Clashscore | 3.34 |
| Rotamer Outliers (%) | 0.37 |
| **Ramachandran plot** | |
| Favored (%) | 97.33 |
| Allowed (%) | 2.67 |
| Disallowed (%) | 0.00 |

### Supplementary Table 6: Oligonucleotides for generating recombinant nucleosomes

| Oligo | Sequence (5′ – 3′) |
|---|---|
| **ND-NCP** | |
| Oligo 1 | ATCGAGAATCCCGGTGCCGAGGCCGCTCAATTGGTCGTAGACAGCTCTAGCACCG<br>CTTAAACGCACGTACGCGCTGTCCCCCGCGTTTTAACCGCCAAGGGGATTACTCCC<br>TAGTCTCCAGGCACGTGTCAGATATATACATCCGAT |
| Oligo 2 | /6FAM/ATCGGATGTATATATCTGACACGTGCCTGGAGACTAGGGAGTAATCCCCTT<br>GGCGGTTAAAACGCGGGGGACAGCGCGTACGTGCGTTTAAGCGGTGCTAGAGCTG<br>TCTACGACCAATTGAGCGGCCTCGGCACCGGGATTCTCGAT |
| **Nick-NCP$_{-6}$** | |
| Oligo 1 | ATCGAGAATCCCGGTGCCGAGGCCGCTCAATTGGTCGTAGACAGCTCTAGCACCG<br>CTTAAACGCACGTACGCGCTGTCCCCCGCGTTTTAACCGCCAAGGGGATTACTCCC<br>TAGTCTCCAGGCACGTGTCAGATATATACATCCGAT |
| Oligo 2 | /6FAM/ATCGGATGTATATAT |
| Oligo 3 | /5Phos/CTGACACGTGCCTGGAGACTAGGGAGTAATCCCCTTGGCGGTTAAAACGC<br>GGGGGACAGCGCGTACGTGCGTTTAAGCGGTGCTAGAGCTGTCTACGACCAATTG<br>AGCGGCCTCGGCACCGGGATTCTCGAT |
| **Nick-NCP$_{-4}$** | |
| Oligo 1 | ATCGAGAATCCCGGTGCCGAGGCCGCTCAATTGGTCGTAGACAGCTCTAGCACCG<br>CTTAAACGCACGTACGCGCTGTCCCCCGCGTTTTAACCGCCAAGGGGATTACTCCC<br>TAGTCTCCAGGCACGTGTCAGATATATACATCCGAT |
| Oligo 2 | /6FAM/ATCGGATGTATATATCTGACACGTGCCTGGAGACT |
| Oligo 3 | /5Phos/AGGGAGTAATCCCCTTGGCGGTTAAAACGCGGGGGACAGCGCGTACGTGC<br>GTTTAAGCGGTGCTAGAGCTGTCTACGACCAATTGAGCGGCCTCGGCACCGGGAT<br>TCTCGAT |
| **Nick-NCP$_{-2}$** | |
| Oligo 1 | ATCGAGAATCCCGGTGCCGAGGCCGCTCAATTGGTCGTAGACAGCTCTAGCACCG<br>CTTAAACGCACGTACGCGCTGTCCCCCGCGTTTTAACCGCCAAGGGGATTACTCCC<br>TAGTCTCCAGGCACGTGTCAGATATATACATCCGAT |
| Oligo 2 | /6FAM/ATCGGATGTATATATCTGACACGTGCCTGGAGACTAGGGAGTAATCCCCTT<br>GGCGG |
| Oligo 3 | /5Phos/TTAAAACGCGGGGGACAGCGCGTACGTGCGTTTAAGCGGTGCTAGAGCTG<br>TCTACGACCAATTGAGCGGCCTCGGCACCGGGATTCTCGAT |
| **Nick-NCP$_{0}$** | |
| Oligo 1 | ATCGAGAATCCCGGTGCCGAGGCCGCTCAATTGGTCGTAGACAGCTCTAGCACCG<br>CTTAAACGCACGTACGCGCTGTCCCCCGCGTTTTAACCGCCAAGGGGATTACTCCC<br>TAGTCTCCAGGCACGTGTCAGATATATACATCCGAT |
| Oligo 2 | /6FAM/ATCGGATGTATATATCTGACACGTGCCTGGAGACTAGGGAGTAATCCCCTT<br>GGCGGTTAAAACGCGGGGGACAGCG |
| Oligo 3 | /5Phos/CGTACGTGCGTTTAAGCGGTGCTAGAGCTGTCTACGACCAATTGAGCGGCC<br>TCGGCACCGGGATTCTCGAT |

*Nucleosomes used for cryo-EM were generated with oligonucleotides lacking the 6FAM label.
